# A CRISPR-drug perturbational map for identifying new compounds to combine with commonly used chemotherapeutics

**DOI:** 10.1101/2023.04.12.536612

**Authors:** Hyeong-Min Lee, William C. Wright, Min Pan, Jonathan Low, Duane Currier, Jie Fang, Shivendra Singh, Stephanie Nance, Ian Delahunty, Yuna Kim, Richard H. Chapple, Yinwen Zhang, Xueying Liu, Jacob A. Steele, Jun Qi, Shondra M. Pruett-Miller, John Easton, Taosheng Chen, Jun Yang, Adam D. Durbin, Paul Geeleher

**Affiliations:** Department of Computational Biology, St. Jude Children’s Research Hospital, Memphis, TN 38105, USA; Department of Chemical Biology, St. Jude Children’s Research Hospital, Memphis, TN 38105, USA; Department of Surgery, St. Jude Children’s Research Hospital, Memphis, TN 38105, USA; Division of Molecular Oncology, Department of Oncology, St. Jude Children’s Research Hospital, Memphis, TN 38105, USA; Center for Advanced Genome Engineering, St. Jude Children’s Research Hospital, Memphis, TN 38105, USA; Department of Cell and Molecular Biology, St. Jude Children’s Research Hospital, Memphis, TN 38105, USA; Department of Cancer Biology, Dana-Farber Cancer Institute, Boston, Massachusetts, USA; Department of Medicine, Harvard Medical School, Boston, Massachusetts, USA; Department of Pathology and Laboratory Medicine, College of Medicine, The University of Tennessee Health Science Center, Memphis, TN 38163, USA

**Keywords:** drug combinations, CRISPR, neuroblastoma

## Abstract

Combination chemotherapy is crucial for achieving durable cancer cures, however, developing safe and effective drug combinations has been a significant challenge. To improve this process, we conducted large-scale targeted CRISPR knockout screens in drug-treated cells, creating a genetic map of druggable genes that sensitize cells to commonly used chemotherapeutics. We prioritized neuroblastoma, the most common pediatric solid tumor, where 50% of high-risk patients do not survive. Our screen examined all druggable gene knockouts in 18 cell lines (10 neuroblastoma, 8 others) treated with 8 widely used drugs, resulting in 94,320 unique combination-cell line perturbations, which is comparable to the largest drug combination screens ever reported. Remarkably, using dense drug-drug rescreening, we found that the top CRISPR-nominated drug combinations were far more synergistic than standard-of-care combinations, suggesting existing combinations could be improved. As proof of principle, we discovered that inhibition of PRKDC, a component of the non-homologous end-joining pathway, sensitizes high-risk neuroblastoma cells to the standard-of-care drug doxorubicin *in vitro* and *in vivo* using PDX models. Our findings provide a valuable resource for the development of improved chemotherapeutic strategies and demonstrate the feasibility of using targeted CRISPR knockout to discover new combinations with common chemotherapeutics, a methodology with application across all cancers.

## INTRODUCTION

Almost all curative cancer treatments result from combinations of multiple chemotherapeutic agents. However, existing drug combinations are often ineffective and can cause severe side effects. The development of improved combinations faces several challenges. Firstly, the drug combinatorial search space is astronomical, with, for example, all possible 2 drug combinations of only 600 drugs yielding 179,700 combinations (given by 600^2^/2 - 600/2). All possible 3 drug combinations of 600 drugs yields approximately 100 million unique combinations, far beyond what could be screened using conventional approaches, without even considering variable compound dosage and timing. This suggests that new high-throughput strategies are needed to capture this search space. Secondly, navigating drug approval for two new drugs simultaneously presents additional regulatory barriers and safety considerations over single-agent approval, which is already a difficult process^1, 2^. This suggests that the development of combinations with compounds already in clinical use should be prioritized as this will present the fewest hurdles to achieving rapid clinical impact. Combinations with standard-of-care drugs could also improve patient outcomes by mitigating toxicities if drug synergies mean that similar anti-tumor activity could be maintained at a lower exposure to broadly cytotoxic chemotherapeutics^3, 4^.

Pediatric neuroblastoma represents a particularly pressing clinical need. Neuroblastoma is the most common pediatric solid tumor^5^, and despite intense study, survival in high-risk patients has remained close to 50%^6^. In recent years, numerous clinical trials have been conducted testing new single-agent targeted chemotherapeutics. Virtually all these clinical trials have failed^6^, typically due to limited tumor response. This has been true even when preclinical and mechanistic evidence has been very convincing; for example, for ALK inhibitors in ALK gain-of-function neuroblastoma^7^, or IGF1R inhibitors in IGF1R overexpressing tumors^8, 9^. These examples are among the few recurrent druggable oncogene aberrations in this disease, which only occur in a small fraction of patients. Further, the disappointing results of even these trials suggests that combinations of drugs eliciting synergistic effects will need to be considered, particularly in the relapsed setting, if there is to be any chance of making clinical progress in this hyper-aggressive disease. This approach has demonstrated promise in other highly aggressive pediatric cancers, with for example the synergistic combination of PARP inhibitors with DNA-damaging agents in Ewing’s sarcoma achieving complete responses in some relapsed patients^10^. Despite this, large-scale drug combination screening studies in pediatric cancers have never been reported.

Here, we have exploited recent observations that CRISPR knockout of most druggable genes mimics pharmacological inhibition of the protein encoded by that gene^11^. Considering this, we designed a CRISPR knockout library targeting 655 known druggable genes. We screened this library to identify druggable gene knockouts that sensitize cell lines to commonly used cancer drugs, providing a large increase in throughput over conventional drug-drug combination screening approaches. Leveraging the resulting dataset, we propose novel therapeutics to combine with doxorubicin, topotecan, cisplatin, and the experimental proteolysis-targeted chimaera (PROTAC) agent JQAD1, which we show are effective using *in vitro* and (for doxorubicin) *in vivo* experiments. Overall, this resource provides a map for the discovery of new chemotherapeutic combinations and demonstrates clinical potential in a disease of extremely high need.

## RESULTS

### Design of CRISPR-drug perturbational screen and selection of cell lines

We first designed a targeted CRISPR gene knockout library (Extended Data Table 1) against 655 druggable genes, targeting each gene with 6 unique gRNAs (see Methods). Our list of druggable genes was based on Behan *et al*.^11^ (Extended Data Table 1). Using DepMap data^12^ as a reference, we removed genes with an expression of < 0.1 log_2_(TPM+1) across the 10 neuroblastoma cell lines used in our screen. We also removed most genes that were universally lethal when knocked out in DepMap cell lines, as it will not be possible to find drug-sensitizing effects for genes whose knockout already kills all cells (note: DepMap does not screen drug-treated cells).

We then set out to screen this gRNA library against a panel of 18 cell lines treated separately with each of 8 different drugs, or vehicle-treated control. The resulting relative abundance of gRNAs targeting each of these 655 genes in the drug-treated vs vehicle-treated cells provides a readout of whether target gene knockout sensitizes a cell line to a drug (Fig. 1a; Extended Data Figure 1a; see Methods for details). This indicates that pharmacological inhibition of this gene product could also present a viable combination with the anchor drug^13, 14^. In total, this experimental design yielded 18 cell lines × 8 drugs × 655 gene knockouts = 94,320 total unique combination-cell line pairs. The specific 8 drugs were doxorubicin, cisplatin, phosphoramide mustard (PM, the active metabolite of cyclophosphamide), etoposide, topotecan, vincristine, and all-trans retinoic acid (standard-of-care neuroblastoma drugs that are all also used broadly to treat many cancers), and the PROTAC JQAD1^15^, which is an EP300 degrader in preclinical development. We screened our gRNA library in 18 Cas9 stably expressing cell lines, 10 of which were neuroblastoma cell lines, and 8 of which were non-neuroblastoma cell lines (Fig. 1b, Extended Data Table 2). These included 4 cancer cell lines (from melanoma, Ewing sarcoma, rhabdomyosarcoma, and colon cancer) and 4 cell lines generated from normal tissues, specifically GM12878, a lymphoblastoid cell line, AC16, a cardiomyocyte cell line, BJ-TERT immortalized fibroblasts and HEK293T cells^16^. The use of non-neuroblastoma cell lines in our experimental design provides valuable information in its own right, but also serves as a statistical outgroup to evaluate the specificity of drug combinations. This provides a baseline to understand whether drug-CRISPR combinations are selectively lethal to neuroblastoma cells, or are simply broadly cytotoxic. Broad cytotoxic combinations would be less desirable clinically, since this would likely reduce the chance of achieving a therapeutic window in patients. Neuroblastoma cell lines were chosen that already had prior exome-wide CRISPR-cas9 screening in addition to dense genomic and perturbational data available in the Cancer Cell Line Encyclopedia and DepMap. We used this information to nominate cell lines that cover the highest clinical need, including 5 cell lines with *TP53* mutations, which are enriched at relapse^17, 18^, and 4 mesenchymal-like cell lines, characterized by a gene expression program associated with chemotherapeutic resistance in neuroblastoma^19–21^ (Fig. 1b).

**Figure 1:**
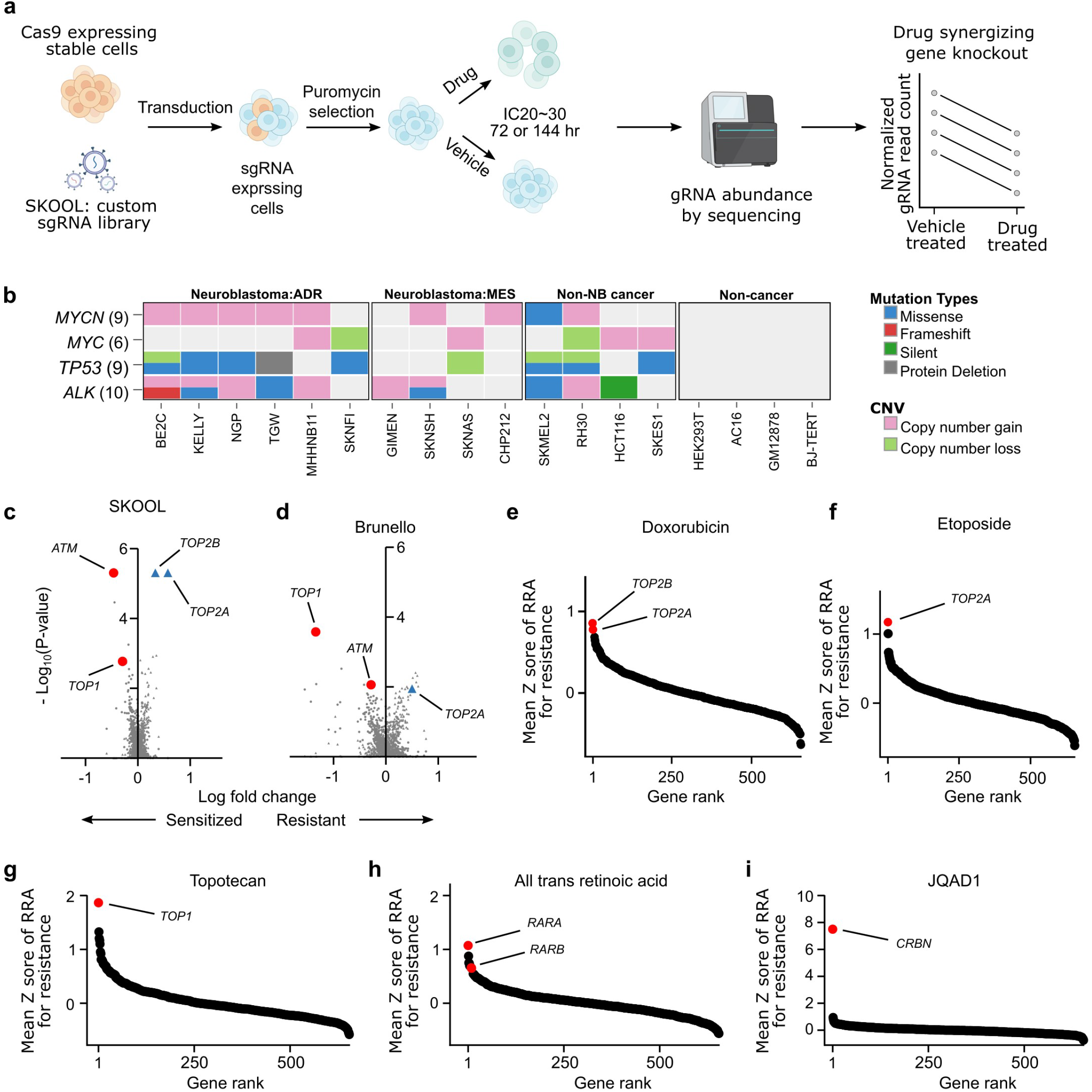
A CRISPR knockout library, targeted to druggable genes, is a viable strategy to prioritize potential synergies with commonly used chemotherapeutics. (a) Simplified schematic of CRISPR-sensitizer screens (detailed schematic available in Extended Data Fig. 1a). (b) Summary of genomics features of cell lines used in the screen. “ADR” refers to predominantly adrenergic neuroblastoma cell lines and “MES” to predominantly mesenchymal cell lines^19, 21, 27^. Mutation profiles were obtained from DepMap. (c) Volcano plot showing the log normalized gRNA fold change (LFC; x axis) and *P*-values (y axis) for each gene knockout in a CX-5461 vs DMSO control treated CHP-134 neuroblastoma cell line. Results were obtained using our 655 gene reduced representation library “SJ KnockOut nOn-Lethal (SKOOL)”. Positive controls, the known sensitizing knockouts (*ATM* and *TOP1*) are highlighted as red circles and known resistance knockouts (*TOP2A* and *TOP2B*) are highlighted as blue triangles. (d) Like (c) but for results obtained for the same genes using the genome-wide Brunello library. (e-h) Waterfall plots showing the gene rank for resistance of the direct drug protein targets in our dataset. Genes are ranked by the mean Z scores across all 18 cell lines of their RRA score for resistance.

### Knockout of known drug-target genes are the top hits for drug resistance

We were first interested in assessing the validity of our 655 gene reduced representation gRNA library. Thus, we performed a targeted CRISPR screen using this gRNA library in CHP-134 neuroblastoma cells treated with the chemotherapeutic CX-5461 and DMSO-treated controls, an experiment we have previously performed using the Brunello genome-wide gRNA library^22^. Our previous genome-wide screen identified that CX-5461 is a topoisomerase inhibitor, with a high affinity for TOP2B. Encouragingly, in this re-screen using our reduced representation library, the top hit for CX-5461 resistance was the primary drug target *TOP2B* (*RRA* = 2.48 × 10^-9^; Note: robust ranking aggregation (*RRA*) scores are the default statistical measure from MAGeCK, the *de facto* computational tool for CRISPR data analysis, see Methods), followed by the secondary drug target *TOP2A* (*RRA* = 8.46 × 10^-9^; Fig. 1c-d; Extended Data Table 3). We also recovered the top sensitizing knockouts, *ATM* and *TOP1* (Fig. 1c-d; Extended Data Table 3). Thus, our reduced representation targeted CRISPR gRNA library delivers results consistent with a widely used genome-wide library. Five of the eight drugs that we screened using the targeted library have known direct protein targets, which can be treated as built-in positive controls to assess the validity of our results. Specifically, doxorubicin and etoposide directly target TOP2^23, 24^; all-trans retinoic acid directly interacts with the retinoic acid and retinoid receptor family (RARA, RARB, RARG, RXRA, RXRB, and RXRG)^25^; topotecan targets TOP1^26^ and JQAD1 is a PROTAC that degrades EP300 by selective recruitment of the E3 ligase receptor cereblon (CRBN)^15^. Cisplatin, PM, and vincristine act on DNA or microtubules and were not included in this analysis. Thus, for each of these 5 drugs, we calculated the mean resistance RRA scores (Z score normalized) across the screens performed in all 18 cell lines. Encouragingly, in all cases, loss of the known protein target of each of these drugs was identified among the top resistance mechanisms, and in the cases of retinoic acid, JQAD1 and topotecan were ranked #1 (Fig. 1e-i, see Extended Data Tables 4 for all RRA scores, positive and negative, across the entire screen). These resistance mechanisms were recovered despite our screens being performed in IC_20_-IC_30_ drug concentrations (Extended Data Figure 1b-c), a dose range suited to identifying drug-sensitizing knockouts, rather than resistance mechanisms (see Methods). Overall, these analyses of known positive controls in the data provide strong evidence of the validity of the screening results.

### Systematic trends in the CRISPR screening data

While these screens recapitulate expected resistance mechanisms and identify new promising combinations (next subsections), the size of this dataset also provides an opportunity to systematically study genetic perturbational effects in cancer cells at a scale beyond what has previously been possible in the context of commonly used chemotherapeutics. We used t-SNE^28^ and UMAP^29^ to cluster the data, which revealed an interesting trend, showing that the experiments primarily clustered by cell line, rather than by drug or outgroup status (Fig. 2a; Extended Data Figure 2a-d; UMAP parameters were selected using an unbiased/automated Monte Carlo approach, see Methods). This is suggestive that a significant proportion of the signal in the data may be cell line specific (NOTE: *this does not mean that all of the signal in the data is cell line specific* (see subsequent sections)). We investigated this by a simple and interpretable orthogonal analysis, where we calculated the number of cell lines crossing a nominally significant RRA < 0.05 for each drug (Fig. 2b) and for each cell line (Fig. 2c; Extended Data Figure 2e). While there were no instances of a gene knockout sensitizing more than 7 of the 10 neuroblastoma cell lines to any given drug (even at a nominal statistical threshold; Fig. 2b), there were several examples of CRISPR knockouts that sensitize a cell line to all drugs screened. For example, knockout of the anti-apoptotic gene *MCL1* sensitized SKMEL2 melanoma cells to all 8 drugs, and knockout of glycine receptor gene *GLRA1* sensitized MHHNB11 neuroblastoma cells to all drugs, which strikingly had little effect in any other cell line (Extended Data Table 4). These behaviors are consistent with previous observations that the effect of pairs of gene knockouts are often cell line specific^30, 31^, and that targeted drug combination effects are highly specific to cellular context^11, 32^. Our results extend these observations to commonly used chemotherapeutics and suggest that caution should be exercised when extrapolating the results of drug combinatorial screens that have used small numbers of biological replicates (see Discussion).

**Fig. 2:**
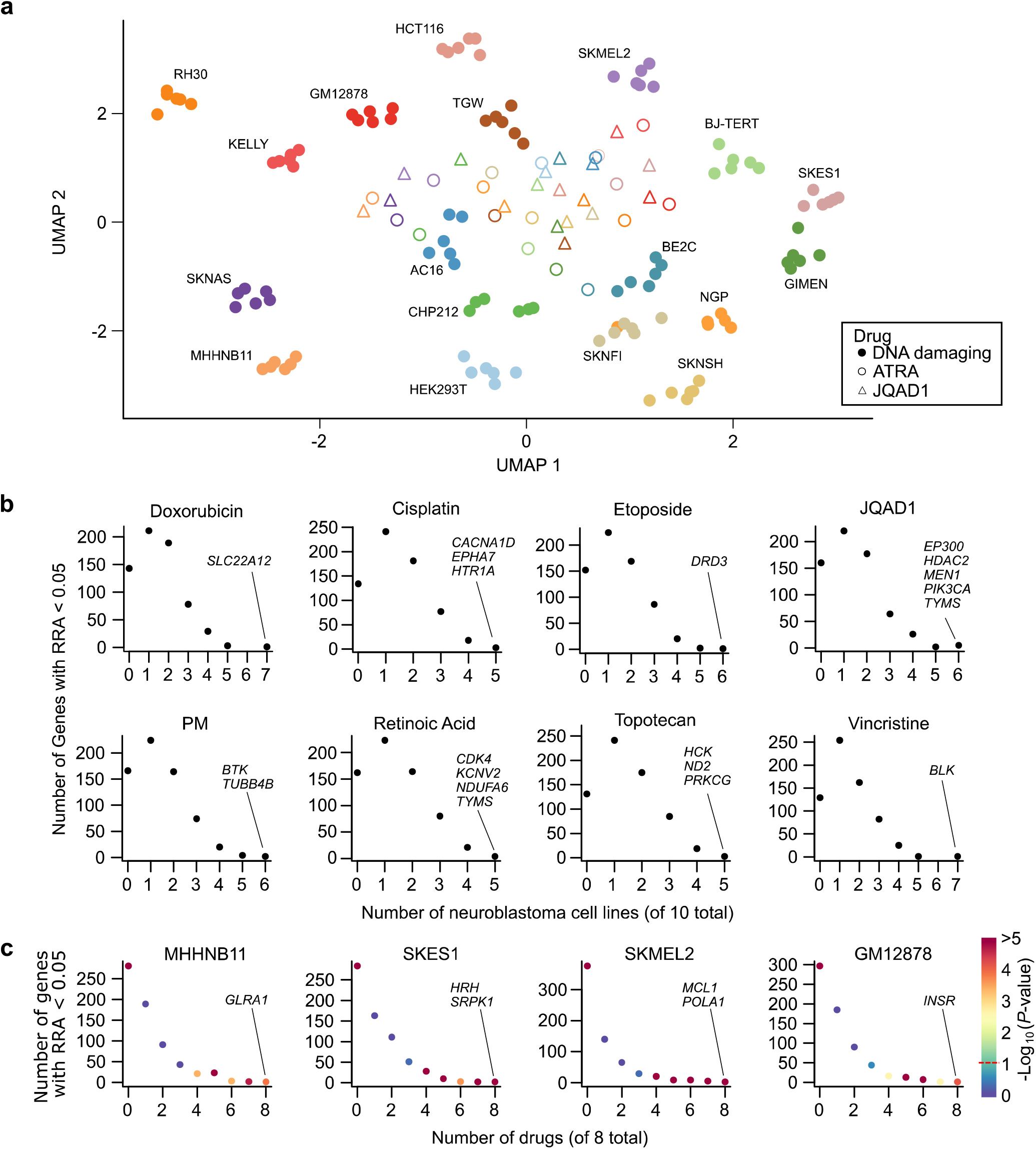
Systematic examination of the screens identifies cell line-specific effects. (a) UMAP representation of all 18 cell lines × 8 drugs = 144 total screens performed. Data are clustered based on fold-change of normalized read counts following each of the 655 gene knockouts profiled. Data points are colored by cell line and the symbol represents whether the cell line was treated with one of the 6 DNA damaging agents (solid circles; cisplatin, doxorubicin, etoposide, phosphoramide mustard, topotecan, or vincristine), all-trans retinoic acid (open circles), or JQAD1 (triangles). (b) Scatterplot showing the number of neuroblastoma cell lines (x axis) that are sensitized to some gene knockout (at RRA < 0.05; y axis) for each of the 8 drugs screened. Genes sensitizing the maximum number of cell lines to any given drug have been highlighted. (c) Scatterplot showing the number of drugs (x axis) that are sensitized to some gene knockout (RRA < 0.05; y axis) for each of the 4 different cell lines screened (all 18 cell lines are shown in Extended Data Figure 2e). Genes sensitizing each of these 4 cell lines to all 8 drugs screened are highlighted. *P*-values were calculated by permutation of all RRA scores.

### A hierarchical Bayesian model can identify gene knockouts that robustly sensitize multiple cell lines to standard-of-care chemotherapeutics

Because of our unique experimental design, and to maximize power to overcome the cell line-specific effects, we developed a set of Bayesian hierarchical models to analyze these data—methods that can be easily implemented in future similar screens. These models have been engineered to account for the uncertainty associated with drug-sensitization fold-changes, estimated from the differences in normalized gRNA read counts in drug-treated vs vehicle-treated control cell lines, which we treat as the model’s outcome variable (see Methods).

To identify the most potent drug-sensitizing gene knockouts across the entire dataset, we first applied this model across all 18 cell lines. Firstly, the resistance hits identified using this model were similar to those identified from RRA scores (Extended Data Table 5) and were again consistent with the known drug targets, with the direct protein targets of etoposide, JQAD1, retinoic acid, and topotecan (*TOP2A*, *CRBN*, *RARA,* and *TOP1* respectively) all recovered as the #1 resistance knockouts when ranked by fold-change (Fig. 3a-h; Extended Data Table 5). Importantly, the model also revealed multiple gene knockouts that drug-sensitize many cell lines in our cohort. Many of the top sensitizing knockouts had a clear biological rationale and several have strong existing experimental evidence: These include *PARP1*, which was the #1 ranked hit (by fold change) for sensitizing cells to the TOP1 inhibitor topotecan (Fig. 3g; Extended Data Tables 5). The synergistic interaction between TOP1 and PARP1 inhibitors has been validated in preclinical models^33^ and represents one of the only standard-of-care drug synergies under active study in pediatric clinical trials^10^. This combination was originally investigated based on biological rationale because TOP1 causes single-stranded DNA breaks, which cannot be effectively repaired in the absence of *PARP1*, but in our data, this was identified without any prior biological knowledge. EP300 knockout was identified as the #6 ranked hit for sensitizing cells to the EP300 degrader JQAD1 (Fig. 3d). PRKDC knockout was identified as the most potent sensitizer to doxorubicin (Fig. 3a), an association recently also reported in hepatoblastoma^34^ and others^35, 36^, and a hit which we explore further in a subsequent subsection. *BCL2L1* knockout was the top-ranked sensitizer for cisplatin (Fig. 3b), #4 for phosphoramide mustard (Fig. 3e), #8 for vincristine (Fig. 3h), and #2 for topotecan (Fig. 3g). *BCL2L1* is an anti-apoptotic protein targetable by navitoclax, which has shown convincing synergy with several chemotherapeutic agents^37^. Combinations of navitoclax with cyclophosphamide^38^, as well as regimens containing doxorubicin and vincristine^39^, are in active clinical investigation^37^. The #2 cisplatin hit *DHFR* (Fig. 3b) is targeted by methotrexate, which is core therapy in osteosarcoma, combined in sequence with cisplatin^40^, and #4 ranked cisplatin hit *CDC7* (Fig. 3b) is supported by existing *in vitro* results^41^. Doxorubicin is synthetic lethal in combination with inhibitors of its #7 ranked hit *CDK1* (Fig. 3a)^42^. *MET* knockout was ranked #2 for sensitization to topoisomerase II inhibitors doxorubicin (Fig. 3a) and etoposide (Fig. 3c), and was ranked #7 for sensitization to cisplatin (Fig. 3b). MET overexpression has been widely implicated in chemotherapy resistance^43^ and MET inhibition has already been reported to sensitize various cancer cells to doxorubicin^44, 45^ and cisplatin^46, 47^. There is also evidence that the screens successfully identified other known resistance mechanisms beyond direct protein targets. For example, it was recently shown that mTOR inhibition represents a general chemoresistance mechanism^48^, and this was indeed identified as a top resistance hit for several of our screened chemotherapeutics (Extended Data Table 5).

**Fig. 3:**
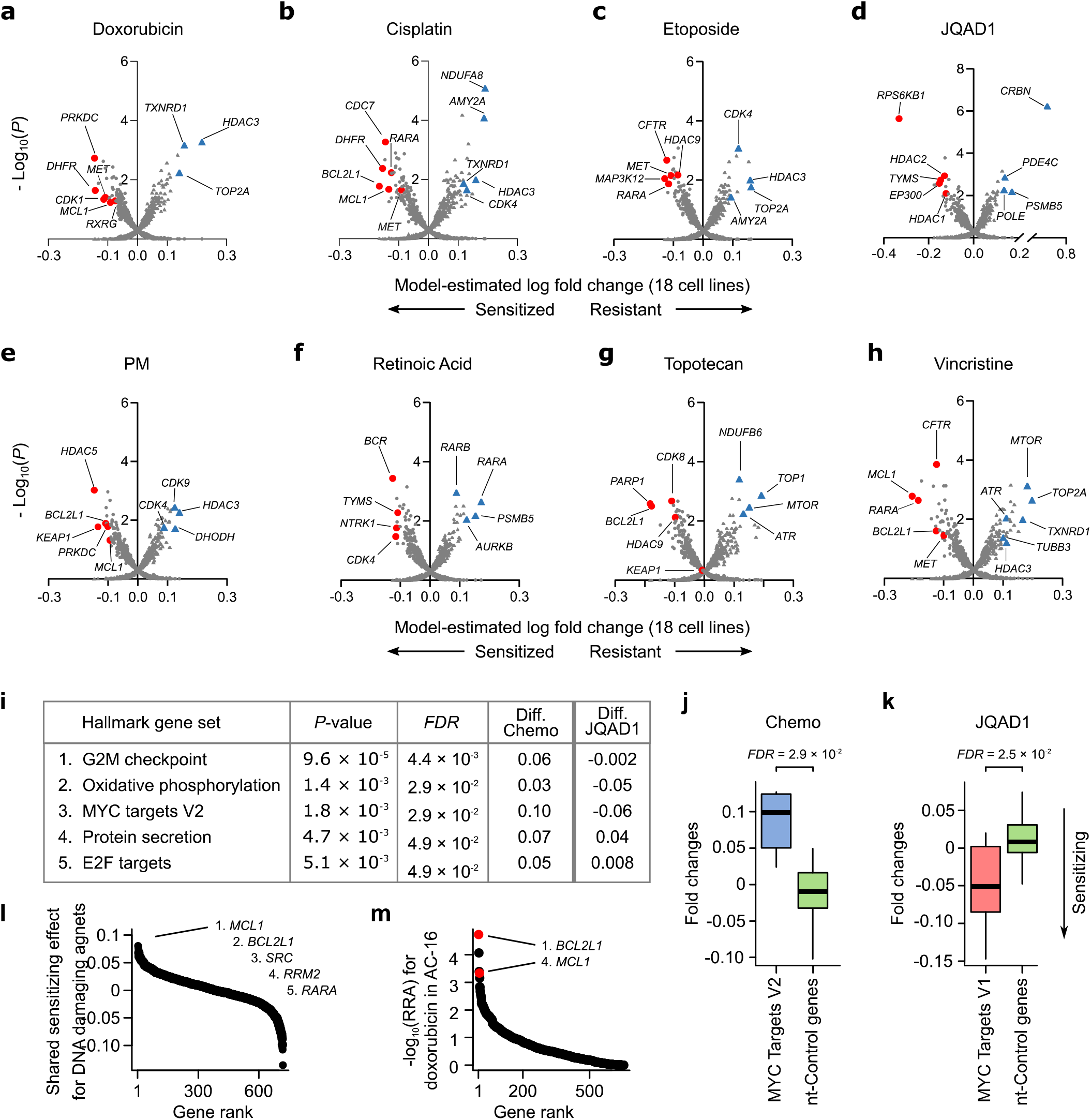
Summary of drug sensitizing hits identified across the entire dataset. (a-h) Volcano plots for each drug showing the estimated mean drug vs mock-treated control normalized gRNA log fold change across all 18 cell lines screened (x axis) and the posterior probability this fold change value is different from 0 (y axis). These values were estimated by fitting our Bayesian Hierarchical model (see Methods) to the 18 fold-change values, and associated measurement uncertainty estimate, obtained for each gene knockout, in each drug. Lower fold change values imply a gene knockout has caused drug-sensitization, with top hits highlighted as red circles. Key resistance knockouts have been highlighted as blue triangles. (i) Top enriched mSigDB “Hallmark” gene sets from a joint model fit across the 6 DNA damaging agents screened. The values in the columns labeled “Diff.” are the difference in fold change for the median gene in the gene set versus the median of the non-targeting control genes; negative values in these “Diff.” columns imply knockouts of genes from this gene set are associated with drug sensitization. The final column shows the directionality of these hits for JQAD1, which are opposite of the 6 DNA damaging agents for the top 3 gene sets. (j) Boxplot showing mean estimated drug vs DMSO normalized gRNA log fold changes (y axis) for genes annotated to the mSigDB’s Hallmark MYC targets gene set (blue box) vs non-targeting control genes (green box). For each gene, these mean fold change estimates were calculated across all 18 cell lines screened using a joint model that considered the 6 DNA damaging agents screened. (k) Like (j), but for JQAD1. (l) Waterfall plot ranking gene knockouts (x axis) by their shared sensitizing log fold change effect (y axis) across the 6 DNA damaging agents, estimated using our Bayesian hierarchical model. (m) Waterfall plot ranking gene knockouts (x axis) by -log_10_ sensitizing RRA scores (y axis) in the AC-16 outgroup cell line. *BCL2L1* and *MCL1*, which are the top broad sensitizer genes in panel (l), are highlighted in red and sensitize this cardiomyocyte cell line to doxorubicin.

Overall, these results show that CRISPR-drug screens can recover known synergies, supporting their utility in identifying synergies with common chemotherapeutics in very high throughput (Extended Data Table 6).

### Gene set functional analysis reveals general chemoresistance mechanisms

We next wondered whether further insights could be gleaned from our data by assessing functional relationships among the hits in the screen. To test this, we developed a gene set analysis approach tailored for these data, which compares the distributions of fold-changes from randomly grouped non-targeting control gRNAs to the groups of functionally related genes (see Methods). We applied this approach to the drug vs vehicle-treated gene-level gRNA fold-change estimates for each drug individually and for the shared effect of the 6 DNA damaging agents considered jointly, performing these functional enrichment analyses at the level of (i) gene families, (ii) genes targeted by the same drug, and (iii) the mSigDB “Hallmarks” gene set, which represents a curated list of well-defined biological processes/pathways. For a combined model assessing the 6 DNA damaging agents, groups of genes whose knockout is likely to slow cell proliferation were most clearly implicated in drug resistance (Fig. 3i; Extended Data Tables 7). This included Hallmark gene sets such as G2M checkpoint (*FDR* = 4.4 × 10^-3^) and MYC targets V2 (*FDR* = 2.9 × 10^-2^; Fig. 3j; Extended Data Table 6). It is known that most common chemotherapeutics are more effective in fast-growing cells because DNA damage is much more likely to be induced during the cell cycle, and these results support the validity of the analyses. Interestingly however, the EP300 degrader JQAD1 provides a compelling exception to this trend, with knockout of genes that support cell cycle surprisingly having the opposite effect and conferring drug sensitivity (Fig. 3i-k, *FDR* = 2.5 10^-2^ for “MYC Targets V1” for sensitization to JQAD1). Thus, knockout of some genes that confer chemoresistance, appear to confer sensitivity to JQAD1. This activity may result from JQAD1’s ability to downregulate MYC^15^—however as an investigational compound, orthogonal drug activity compared to the existing standard-of-care is an extremely desirable characteristic, as it is suggestive of potential to confer clinical benefit by independent action^49^.

### Broadly chemo-sensitizing gene knockouts can be identified by using a Bayesian model that shares information across related drugs

In addition to the *MET* and *BCL2L1* examples discussed above, several hits could be identified in the analyses where genes appeared among the top sensitizers to multiple drugs that were independently screened. This is consistent with the mechanistic convergence of many chemotherapeutics on processes like DNA-damage, cell cycle, and apoptotic pathways (Extended Data Tables 6). This is also consistent with the results of the functional enrichment analysis (Fig. 3i). To formally identify these broad sensitizers in a statistically coherent framework, we extended our Bayesian model to share information across related drugs (see Methods), specifically the 6 DNA damaging agents which tended to co-cluster (Fig. 2a; doxorubicin, etoposide, cisplatin, topotecan, vincristine and phosphoramide mustard). These analyses revealed several broad sensitizers (Extended Data Table 8) with the top-ranked gene being *MCL1*, followed by the aforementioned BCL2L1 (Fig. 3l). This is interesting because *MCL1* has already been suggested as a potent sensitizer to several drugs in multiple diseases. These include the cytotoxic chemotherapeutics paclitaxel and docetaxel in breast cancer^50, 51^, with broader applications reviewed in Bolomsky *et al.*^52^

These broad hits can also be used to highlight an interesting feature of our experimental design. MCL1, while synergizing with multiple chemotherapeutics in neuroblastoma cell lines, also promotes cytotoxicity in our outgroup; for example, *MCL1* knockout also has strong synergy with doxorubicin in our cardiomyocyte cell line (Fig. 3m). Cardiotoxicity is an often-fatal side effect of doxorubicin treatment^16, 53–56^ and such a result suggests that this risk may be potentiated by MCL1 inhibition, an idea for which there is already some support in the literature^57^. Thus, while a joint model applied to these data can identify broadly synergizing knockouts, such targets may be a high risk for toxicity if selectivity is not considered. Motivated by this concept, we next introduce the idea of “selective drug synergy”, where drug sensitization in our neuroblastoma cell lines is compared to our outgroup, thus assessing a gene knockout’s potency *and* selectivity, which could help identify drug synergies more likely to have a therapeutic window *in vivo*.

### Gene knockouts selectively sensitize neuroblastoma cell lines to standard-of-care drugs

To test whether we could identify neuroblastoma-selective hits, we further extended our statistical models to handle case/control designs (see Methods), as well as probing covariates such as genomic features (Fig. 4a-i; Extended Data Table 9).

**Fig. 4:**
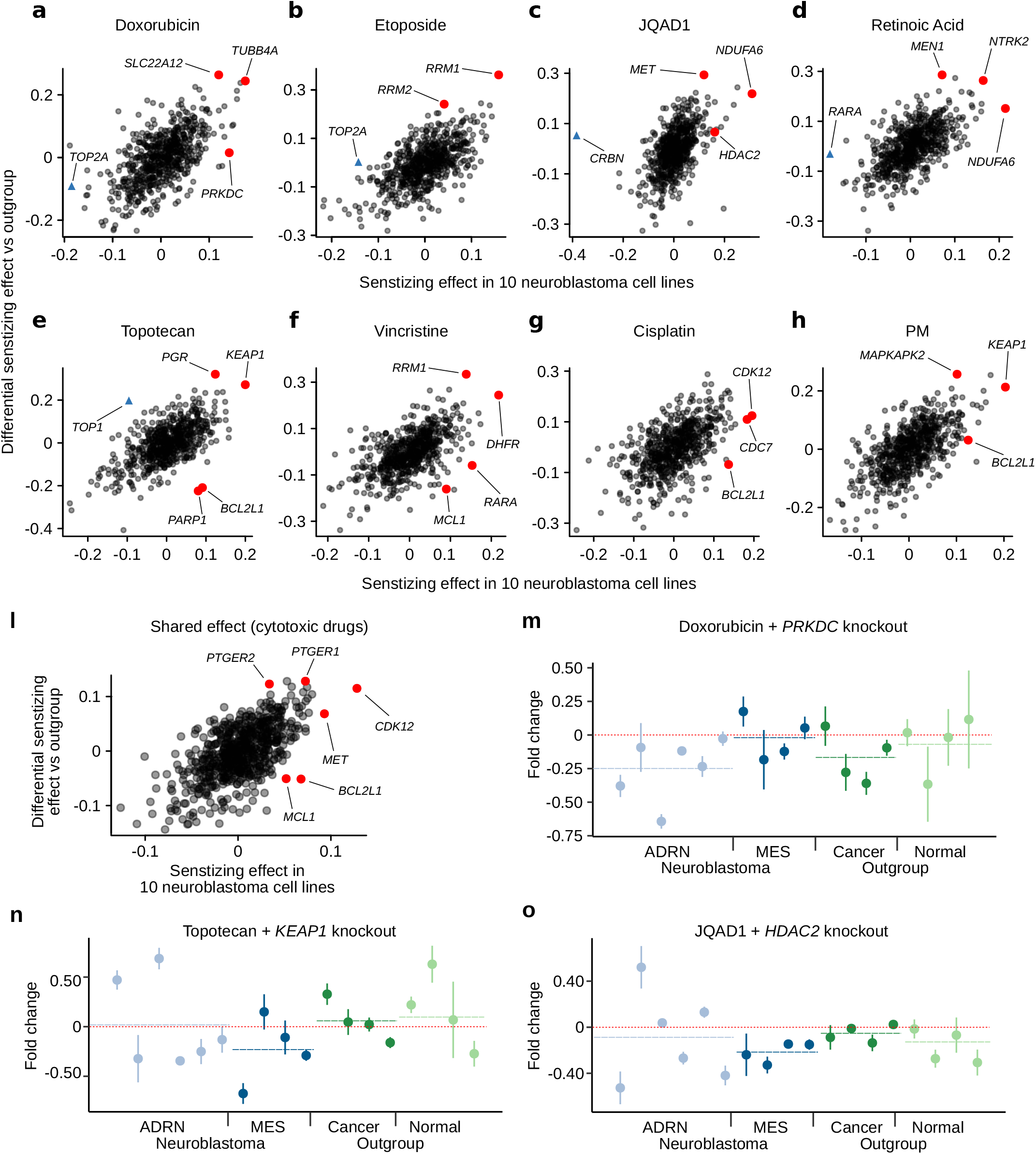
Summary of differential drug sensitizing hits identified in neuroblastoma cell lines vs the outgroup cell lines. (a-h) Scatterplots for each drug showing the estimated mean drug vs mock-treated control normalized gRNA log fold change across in the 10 neuroblastoma cell lines (x axis) and the differential sensitization effect between the 10 neuroblastoma and 8 outgroup cell lines (y axis). Higher values on the x axis imply greater sensitization in neuroblastoma and higher values on the y axis imply greater sensitization in neuroblastoma relative to the outgroup. (i) Like (a-h) but for the shared effect for the 6 DNA damaging agents, estimated using our Bayesian hierarchical model. (j) Doxorubicin vs mock-treated control normalized gRNA log fold changes following *PRKDC* knockout (y axis) for all 18 cell lines screened. Lower values imply sensitization. Neuroblastoma cell lines are colored blue and outgroup cell lines are green. The order of the cell lines (x axis) is the same as Fig. 1b. Whiskers represent the standard error of the mean, estimated from the 6 gRNAs targeting each gene. (k) Like (j) but for topotecan and *KEAP1* knockout. (l) Like (j) but for JQAD1 and *HDAC2* knockout.

Prior knowledge of the selective drug synergy landscape of neuroblastoma is currently almost non-existent, thus, unlike above, it is difficult to identify expected selective synergies in this context. However, *RRM1* and *RRM2* may represent partial exceptions to this, which we identified among the top hits sensitizing to etoposide and vincristine. Recently, combined RRM2 and CHK1 inhibition was shown to be synergistic and non-toxic in neuroblastoma xenografts owing to replicative stress due to stalled replication forks^58^, a process in which TOP2, the target of etoposide, also plays a key role^59^. However, most of the neuroblastoma selective synergies are novel. Interestingly, for some of the most potent drug synergizing knockouts identified across the full dataset, these results suggest their activity is stronger in the outgroup than the neuroblastoma group. *MCL1* and *BCL2L1* represent two examples of this, evident in the topotecan, vincristine, and cisplatin screens (Fig. 4e-g). This suggests that combinations of inhibitors of these targets with DNA-damaging agents should be approached with caution in neuroblastoma because while there may be potent synergistic activity in neuroblastoma cells, there is also strong potential for activity in normal cell types. In the context of other diseases, there is already some evidence that pharmacological inhibition of these targets in combination with DNA-damaging agents causes toxicities^60–62^.

In general, while our number of screened cell lines and combinatorial perturbations is far larger than previous screens in neuroblastoma, it is still not large enough to confidently resolve the context specificity of many hits. However, there is some tentative evidence of orthogonal activity of some nominated combinations, with differential activity emerging on the background of, for example, the expression of mesenchymal-like genes—a key drug resistance state in neuroblastoma ^19^. For example, the knockout of *PRKDC* appears to have a greater sensitizing affect in adrenergic neuroblastoma cell lines (Fig. 4j), whereas *KEAP1*’s effect on topotecan (Fig. 4k), and *HDAC2’s* effect on JQAD1 (Fig. 4l), are both sensitizing in mesenchymal-like cell lines. These observations provide tentative evidence that such context-specific synergies may exist for standard-of-care drugs in neuroblastoma, but broadly resolving these and deconvolving the various confounding factors will require detailed prospective experimental work. Thus, while we explore a selection of hits in detail below, we have also made the data and processed results available in a graphical web-based interface that can be used to motivate new studies dissecting the promising selective hits (available at https://stjude.shinyapps.io/CASAVA/).

### Synergies identified in high-throughput pooled CRISPR-drug screens translate to other genetic and pharmacological assays

We were next interested in assessing the robustness with which CRISPR-drug nominated synergies could be recapitulated with other *in vitro* and *in vivo* assays. We first assessed this using an orthogonal genetic perturbational assay, specifically shRNA knockdown of the CRISPR-nominated targets, for putative synergistic and non-synergistic interactions. First, we tested the knockdown efficiency of individual shRNAs against *PRKDC, HDAC2*, *KEAP1*, and *MET* to select the most efficient one (Extended Data Figure 3a; see Methods). After selection, we knocked down four individual genes in a subset of cell lines (10 for *PRKDC*, 11 for *HDAC2*, *KEAP1*, and *MET*, a total of 43 knockdown experiments, Extended Data Figure 3b) and treated with the corresponding drugs (IC_50_ of doxorubicin in *PRKDC* knockdown, IC_50_ or max 10 µM of JQAD1 in *HDAC2* knockdown, IC_50_ of topotecan in *KEAP1* knockdown, and IC_50_ or max 10 µM of cisplatin in *MET* knockdown). Encouragingly, we observed a strong positive correlation between the shRNA and CRISPR-based perturbations (Fig. 5a – d, Extended Data Figure 3). Thus, the results from the pooled CRISPR screen could be clearly replicated in a different low-throughput genetic assay.

**Fig. 5:**
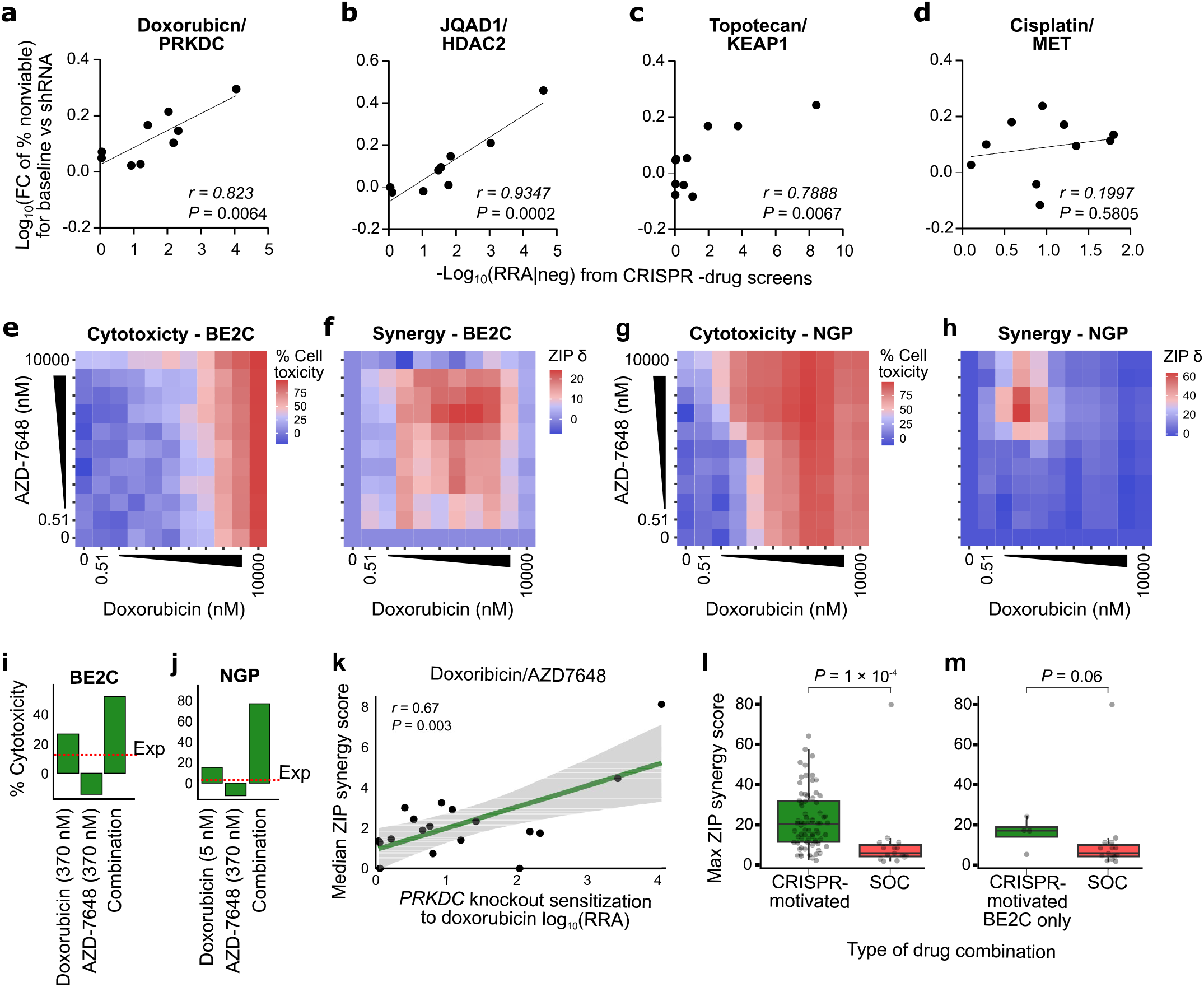
Synergies identified in high-throughput pooled CRISPR-drug screens translate to other genetic and pharmacological assays. (a) Scatterplot of the -log_10_ sensitizing RRA scores from the CRISPR -drug screens (x-axis) against the -log_10_ fold change of cell toxicity percentage upon knockdown of *PRKDC* using single shRNAs (y-axis) in 10 cell lines. For the shRNA knockdown, potency changes were estimated from the IC_50_ of doxorubicin. See Extended Data Table 10 for source data. (b) Like (a) but *HDAC2* shRNA knockdown in 11 cell lines treated with JQAD1. (c) Like (a) but *KEAP1* shRNA knockdown in 11 cell lines treated with topotecan. (d) Like (a) but *MET* shRNA knockdown in 11 cell lines treated with cisplatin. (e) Heatmap matrices of percent cytotoxicity (1 - cell viability) in BE2C cells conferred by treatment with doxorubicin (x axis) and AZD-7648 (y axis). Each matrix represents the average of three independent experiments. (f) Heatmap matrices of synergy scores derived from cytotoxicity values in (e). All synergy scores δ were calculated based on the zero-interaction potency (ZIP) model. Combinations conferring synergy have ZIP scores of >0. (g) Like (e) but for the NGP cells. (h) Like (f) but for the NGP cells. (i) Bar plot showing the cytotoxicity values for the region of max synergy in the BE2C cells (panel (f)). The red dashed line shows expected cytotoxicity under additivity. (j) Like (i) but for the NGP cells. (k) Scatterplot of sensitization RRA scores in each cell line for doxorubicin sensitization by *PRKDC* knockout (x axis) versus overall synergy scores (y axis) from the AZD7648/doxorubicin drug-drug screen See Extended Data Table 11 for source data. (l) Boxplot of maximum synergy scores achieved in our complete set of CRISPR-motivated drug combination screens (green box) and standard-of-care (SOC) motivated drug combinations in the BE2C cell line (red box). *P*-values were calculated from a 2-sided Wilcoxon rank sum test. (m) Like (l) but showing only CRISPR-motivated drug combinations in the BE2C cell line.

Next, to test the translation of gene targets to pharmacological inhibition of protein products, we used dense drug-drug rescreening for these same hits (PRKDC/doxorubicin, KEAP1/topotecan, HDAC2/JQAD1, and MET/cisplatin). The drugs used to target the CRISPR-nominated genes were AZD7648 for PRKDC, dimethyl fumarate (DMF) for KEAP1, panobinostat for HDAC2, and cabozantinib (CAB) for MET. For each pair of drugs, we used high-throughput robotic handling to perform these rescreens in dense 10 × 10 matrices of drug concentrations, with the standard half-log scale concentration (10 doses from 10 µM to 1 nM for AZD7648, DMF, and CAB, and 10 doses from 0.01 µM to 0.5 pM for panobinostat), representing a comprehensive assessment of potential synergies (Fig. 5e-j). In all cases, very strong synergy was observed in the drug-drug combination rescreens (Fig. 5k; Extended Data Figures 4 and 5). Overall, the results suggest that CRISPR-drug screening results can translate to similar drug-drug combination screening results, especially when the compound has a high selectivity for its target. We also performed similar dense 10 × 10 concentration drug-drug rescreens for each pair of standard-of-care neuroblastoma drugs (Cisplatin, PM, ATRA, Topotecan, Doxorubicin, and Vincristine; 15 total combinations) using the BE2C cell line and compared the synergy scores of the CRISPR-motivated compound pairs to synergy scores when standard-of-care drugs are paired. Remarkably, the CRISPR-motivated compound pairs were far more synergistic (*P* = 1×10^-4^, Fig. 5l-m). Thus, it is likely that the drug combinatorial search space contains drug pairings that can improve upon the existing standard-of-care combinations and high throughput drug-CRISPR screens represent a reasonable means to identify these.

### PRKDC inhibition represents a mechanistically plausible combination with doxorubicin in high-risk neuroblastoma, with evidence of synergistic activity *in vivo*

The evidence above suggests that PRKDC inhibition represents a viable clinical option in combination with doxorubicin in neuroblastoma. Thus, we performed several additional assays to assess this possibility. The synergistic relationship between doxorubicin and PRKDC inhibition is mechanistically plausible, as doxorubicin’s primary mechanism of cytotoxicity is DNA double-strand breaks caused by trapping of topoisomerase 2 to DNA. These breaks are repaired in part by the non-homologous end joining (NHEJ) pathway, where PRKDC plays a critical role.

Upon DNA damage, PRKDC undergoes phosphorylation of residue ser-2056, causing a conformational change required for efficient end processing and DNA repair^63^. Thus, we performed western blots to examine the induction of phosphorylation at ser-2056 (pPRKDC) following doxorubicin treatment in the neuroblastoma BE2C and GIMEN cell lines (0.4 μM in BE2C, 0.03 μM in GIMEN – the approximate IC_50_ of these cell lines used in all experiments in this section; PRKDC knockout strongly synergized with doxorubicin in BE2C in the CRISPR screen, but did not sensitize GIMEN cells). The blots showed that single agent doxorubicin-induced pPRKDC in BE2C, but not GIMEN, suggesting NHEJ was only strongly activated in BE2C, the comparatively doxorubicin-resistant cell line (Fig. 6a). Additionally, we confirmed the on-target activity of AZD7648, with single-agent treatment at 3 μM repressing pPRKDC in both cell lines (neither cell line was sensitive to single-agent AZD6748). Surprisingly however, in BE2C, the combination of doxorubicin and AZD7648 markedly increased pPRKDC over doxorubicin alone at 72 h (Fig. 6a), suggesting NHEJ activity had increased, a seeming contradiction worthy of further investigation. Thus, we next tested the level of induction of apoptosis in each cell line using a luminescent caspase 3/7 assay and found that despite the high pPRKDC levels in the doxorubicin/AZD7648 treated BE2C, these cells also had the highest levels of apoptotic response (Fig. 6b, *P* < 1 × 10^-4^ compared to doxorubicin alone). Unsurprisingly, the combination had little effect in potentiating apoptosis in GIMEN. We hypothesized these trends were likely due to much higher levels of DNA damage in the combination-treated BE2C cells. Indeed, at 72 h there was a 4-fold increase in γH2AX foci and a clear increase in overall DNA damage as estimated by a comet tail assay (Fig. 6c-d, *P* < 1 × 10^-4^ for doxorubicin treated vs combination treated in both assays). Interestingly, the number of γH2AX foci in BE2C was approximately 10 times higher than GIMEN when both cell lines were treated with an approximate IC_50_ of doxorubicin (Fig. 6c; Extended Data Figure 6). This is consistent with BE2C being a *TP53* mutant generally chemo-resistant cell line, requiring much higher levels of DNA damage to induce cell death programs. Since the relative use of DNA-repair pathways is also cell cycle-dependent, we further tested the combination effects when arresting cells in G0/G1 or G2 phases of the cell cycle, but neither could explain the increase in pPRKDC (Fig. 6e, Extended Data Figure 7; see Methods). Finally, we performed a cell-based NHEJ activity assay, collecting data at 6 time points from 0 to 72 h. In BE2C, at later time points, NHEJ activity was higher in the combination-treated cells than in cells treated with doxorubicin alone and there was little evidence of induction of NHEJ in GIMEN (Fig. 6f, Extended Data Figure 8). Thus, it seems likely that early DNA damage in the combination-treated BE2C cells leads (counterintuitively) to higher NHEJ activity in surviving cells at later time points, despite AZD7648 actively inhibiting NHEJ over the time course. Overall, these results suggest that differences in the reliance on the NHEJ pathway could predict the effectiveness of PRKDC inhibitors in combination with doxorubicin, and the main mechanism driving synergy is massive potentiation of DNA damage in cells dependent on NHEJ, which can be sufficient to induce cell death even in generally chemoresistant cells. Thus, potentiating doxorubicin activity in this context has the potential to address a clear clinical need^18^. As a final proof of principal, we tested this combination of doxorubicin and AZD7648 *in vivo* using mouse models of neuroblastoma. We used a pharmacologically relevant dosage of each drug based on existing pharmacokinetics data of doxorubicin^64, 65^ and AZD7648^36^ (see Methods). In the first set of experiments, we implanted the BE2C cell line. Encouragingly, the doxorubicin/AZD7648 combination also had clear synergistic activity against this cell line *in vivo*. We observed no significant effect of either AZD7648 or doxorubicin alone, but a clear reduction in tumor volumes when the combination was administered (Fig. 6g).

**Fig. 6:**
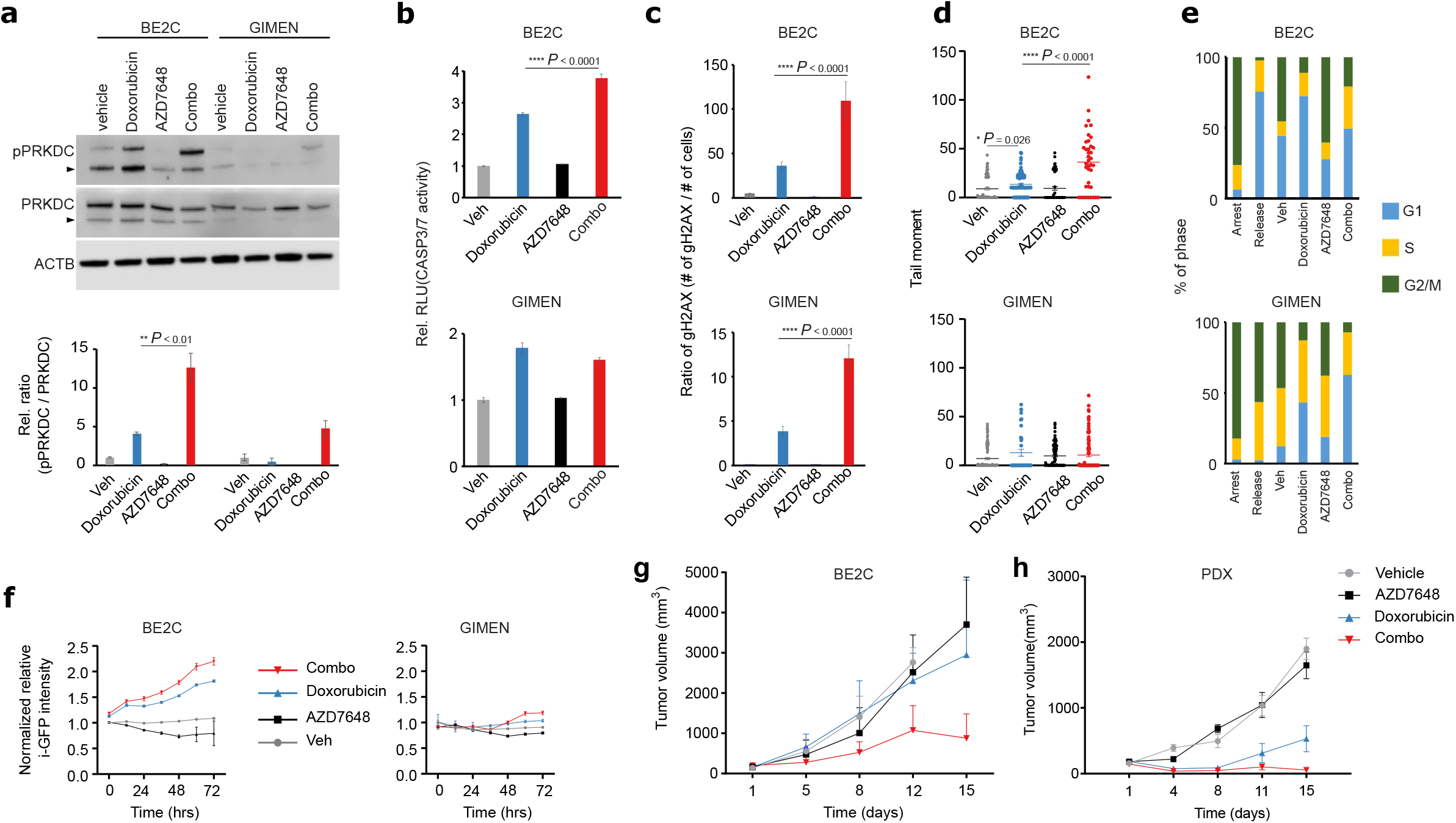
PRKDC inhibition represents a mechanistically plausible combination with doxorubicin in neuroblastoma, with evidence of synergistic activity *in vivo*. (a) Western blot for phosphorylated PRKDC (active PRKDC) in BE2C and GIMEN cell lines. Results are quantified in the lower panel, where doxorubicin (0.4 µM in BE2C, 0.03 µM in GIMEN) activated PRKDC compared to control (veh) whereas AZD7648 (3 µM) inhibited PRKDC. The N-terminus fragment of PRKDC is indicated by the triangle. (b) Luminescent caspase 3/7 assay quantifying the level of apoptosis (y axis) in vehicle, doxorubicin, AZD-7648, or combination (x axis) treated cells. The results for BE2C are shown in the upper panel and GIMEN in the lower panel. (c) Like (b) but for γH2AX foci from immunofluorescence assay. See Extended Data Table 14 for source data. (d) Like (b) but for comet tail assay. (e) Bar plot showing the percentage of cells (y axis) in different phases of the cell cycle (colors), following either cell cycle arrest, 24h following release, or treatment with vehicle, doxorubicin, AZD7648, or the combination for 72h (x axis). (f) Line plot showing the results of a cell-based assay for NHEJ using i-GFP quantifying the activity of NHEJ (y axis) in BE2C and GIMEN cells over 72h (x axis). (g) Changes in tumor growth for BE2C xenografts (vehicle = 3, AZD7648 = 4, Doxorubicin = 5, Combo = 5). Data presented as mean ± SEM. Unpaired t-test for comparison of doxorubicin with combination therapy: *P*-values were 0.035(day 8), 0.012 (day 12), 0.00016 (day 15). (h) Changes in tumor growth for SJNB14 PDX xenografts (vehicle = 4, AZD7648 = 4, Doxorubicin = 4, Combo = 5). Data presented as mean ± SEM. Unpaired t-test for comparison of doxorubicin with combination therapy: *P*-values were 0.054 (day 11), 0.00006 (day 15).

We conducted a second *in vivo* study, this time xenografting a human PDX model (NB14; previously established at St. Jude^66^) of high-risk MYCN amplified neuroblastoma. AZD7648 had no detectable activity as a single agent, but, as in the previous experiment, strikingly potentiated the activity of doxorubicin, exhibiting strong control over tumor growth in all mice (Fig. 6h). Remarkably, this PDX is biologically independent of our cell line discovery cohort, suggesting broad relevance of this synergy in neuroblastoma and that this combination should be further evaluated in this disease. Overall, these mechanistic and *in vivo* experiments show that a large-scale CRISPR-drug perturbational map can be used to prioritize potential synergies with common chemotherapeutics and that these hits can be used to nominate new drug combinations with clinical potential.

## DISCUSSION

Drug combinations are required for essentially all curative cancer treatment strategies. However, the number of possible drug combinations is much larger than can be reasonably screened with existing approaches. The two largest existing drug combinatorial screens are the NCI’s ALMANAC study^67^, which screened the NCI60 cell line panel, and a very recent screen in 125 colorectal, breast, and pancreatic cancer cell lines (published in *Nature* in 2022)^32^. Both studies screened a total of approximately 100,000 unique combination-cell line pairs (compared to 94,320 here). These previous studies employed reasonably dense drug-drug combinatorial screening designs, where drugs were screened in 384 well plates in 3 × 3 or 2 × 7 grids of drug concentrations. Even with modern robotic handling, this approach is extremely resource intensive, thus limiting the number of cell lines captured. Hence it is not surprising that most cancer types are absent from these studies, with for example almost no representation of pediatric cancer cell lines. To broadly capture the astronomical drug-drug combinatorial search space, resource-efficient approaches will need to be developed and deployed. Here, to find new and effective combinations of treatment modalities, we performed large-scale targeted CRISPR knockout screens creating a map of potential new drug synergies with existing commonly used chemotherapeutics. To address the clear clinical need, we emphasized high-risk neuroblastoma cell lines, although several additional cancer and normal cell types were represented. We have made this dataset available to the research community as a resource to prioritize potential standard-of-care drug-drug synergies and to study drug mechanisms and resistance. We have used this resource to discover new drug combinations with clinical potential, which we have demonstrated to be effective *in vivo* for doxorubicin and PRKDC inhibition using patient-derived xenograft models. The observation that CRISPR knockout of a gene typically mimics pharmacological inhibition of the gene’s protein product has now been widely exploited as a massively high-throughput proxy for single-agent drug screening. This has led to very large dependency maps^12^, now profiling over 1,000 cancer cell lines, which have been an immensely successful resource for drug repurposing^68^ and drug discovery^69^, especially in pediatric cancer^70^. However, CRISPR knockout has not yet been extensively deployed to increase the throughput of drug combination discovery, although a few smaller-scale combinatorial knockout studies have been performed^12, 32, 71, 72^. Some of the trends described in these previous studies were also evident in our data, for example, that cell line-specific synergies appear to be common, although we extend this idea to commonly used chemotherapeutics. Additionally, broad synergies, evident across a range of cell types are also common, which we were able to determine by screening an outgroup, a step typically overlooked in drug combination screening. However, even given these “too narrow” or “too broad” synergies, we were still able to identify many examples of synergies that appear to have context specificity in neuroblastoma, which was in part possible due to the development of novel statistical modeling. To promote the widespread use of these models, we have made these analytical tools available for future studies. Arguably, the major limitation of our study may be that, while it is on par with the largest drug combination screens ever performed, context specificity of synergies could likely be resolved in greater detail if screens were carried out using even larger numbers of cell lines in the future, or if data from many screens were aggregated. Interestingly, the proposed approach is highly scalable and effective and can be scaled up for many (if not most) cancers where the drug-drug combinatorial landscape remains almost entirely unexplored. Overall, CRISPR-drug combinatorial screens are effective for the discovery of potentially clinically relevant combinations with existing chemotherapeutics, which has the potential to impact patient care across a wide range of cancer types.

## METHODS

### Animals

All murine experiments were done in accordance with a protocol approved by the Institutional Animal Care and Use Committee of St. Jude Children’s Research Hospital. Around 5 weeks old female NSG mice (NOD.Cg-Prkdc scid Il2rg tm1Wjl /SzJ) were purchased from St Jude Children’s Research Hospital Animal Research Resource and housed in pathogen-free conditions with food and water provided ad libitum. To establish SJNB14-PDX model, PDX tumor was finely minced with sterile scissors and blade in a sterile petri dish. ∼50 μl of minced tumor tissue was subcutaneously engrafted on the right flank of NSG mice. For generating BE2C xenograft, BE2C cells (5 × 10^6^/mouse) in 100 μl in Matrigel (Corning, 354230) were injected subcutaneously on the right flank of NSG mice.

### Cell culture and generation of Cas9 stably expressing cell lines

18 cell lines and their associated Cas9-expressing cell lines (total 36) were cultured in the indicated culture condition (Extended Data Key resources) and maintained in a mycoplasma-free condition. For CRISPR screens, Cas9 stably expressing cell lines were generated or obtained. Cas9 expressing SKNFI, TGW, SKNSH, SKES1, SKMEL2, RH30, HEK293T, BJ-TERT, AC16, and GM12878 were generated by transducing lentiviral Cas9-2A-Blast, followed by blasticidin selection. Cas9 expressing MHHNB11, BE2C, NGP, KELLY, CHP212, GIMEN, and SKNAS were provided by Dr. Adam Durbin. Cas9 expressing HCT116 was purchased from Horizon Discovery (Cat # Cas9-002). Cas9 activity in 18 Cas9-expressing cell lines were verified and it was over 85% on average using the Cas9 activity assay described at Method details.

### Generation of CRISPR KO lentiviral library

sgRNAs for the human CRISPR KO library were first designed using CRISPick^73, 74^. The top 30 sgRNAs for each gene then underwent an additional round of filtering using in-house off-target analysis to identify highly unique sgRNAs. Up to 6 sgRNAs per gene were selected for the library along with non-targeting controls making up ∼10% of the final library. The sgRNA sequences are described in Extended Data Table 1. Library oligos were designed according to Sanson et al.^73^. The oligo pool was synthesized by TWIST Bioscience. Library amplification and Golden Gate cloning into the pLentiGuide-Puro backbone (Addgene #52963) were performed according to Sanson et al.^73^. The plasmid library was amplified and validated in the Center for Advanced Genome Engineering at St. Jude as described in the Broad GPP protocol. The only exception being the use of Endura DUOs electrocompetent cells. The St. Jude Hartwell Center Genome Sequencing Facility provided all NGS sequencing. Single end 100 cycle sequencing was performed on a NovaSeq 6000 (Illumina). Validation to check gRNA presence and representation was performed using calc_auc_v1.1.py (https://github.com/mhegde/) and count_spacers.py^75^. Viral particles were produced by the St. Jude Vector Development and Production laboratory. CRISPR KO screens were analyzed using Mageck-Vispr/0.5.7^76^

### Cas9 activity assay

Using a Cas9 activity assay kit (Cellecta), Cas9 activity was measured by following the manufacture’s protocol. Briefly, Cas9-expressing cells were infected by CT-active [CT-A] or CT-background [CT-B] premade lentiviruses and maintained the infected cells for 10 days to avoid 100% confluency. After 10 days, the cells were analyzed by flow cytometry to measure the changes in GFP levels and Cas9 activity was determined (Extended Data Figure 9 and Extended Data Table 13).

### Compounds and pharmacological profiling

Six standard drugs, cisplatin (CDDP), doxorubicin, etoposide, topotecan, vincristine (VCR), and all-trans retinoic acid (ATRA) were obtained from Medchemexpress (USA), and phosphoramide mustard (PM) was obtained from Niomech IIT GmbH (Germany). JQAD1 was provided by Dr. Jun Qi (Dana-Farber Cancer Institute). As a broad-spectrum cell death compound (positive control for cell death), staurosporine (Medchemexpress) was used for evaluating cell viability. All compounds were reconstituted in DMSO, except CDDP. CDDP was reconstituted in normal saline or a mixture of DMSO and HCl (30v:1v). For pharmacological profiling, individual standard drugs were dispensed by Echo 650 (Labcyte) into white 384-well plates in a dose-dependent manner, followed by plating Cas9 expressing cells with desired numbers (500 or 1000 cells per well). After 3 days (CDDP, PM, doxorubicin, etoposide, topotecan, VCR) or 6 days (ATRA, JQAD1) incubation, CellTiter-Glo (Promega) assay was performed to determine viability and IC_20_, IC_30_, and IC_50_ was calculated by fitting Hill Slope equation (Extended Data Figure 10). An IC_20_ to IC_30_ was used for negative selection in our CRISPR screens. For the combinatorial drug screen, selected partner compounds, AZD7648 (PRKDC inhibitor), panobinostat (HDAC2 inhibitor), dimethyl fumarate (KEAP1 inhibitor), and cabozantinib (MET inhibitor) were purchased from Medchemexpress.

### CRISPR screening

Our Cas9-expressing cell lines were infected with our SKOOL library at MOI (∼0.3), followed by puromycin selection. In between days 10 – 12, once survived knockout cells reached at desired numbers to maintain representation, they were treated with individual standard drugs (IC_20_-_30_) for 3 days (CDDP, PM, doxorubicin, etoposide, topotecan, VCR) or 6 days (ATRA and JQAD1). Genomic DNA was extracted using the PureLink genomic DNA kit (Invitrogen) and the sgRNA sequences were recovered by genomic PCR analysis, followed by deep sequencing using NovaSeq for paired-end minimum length 75 bp read (Illumina). Sequencing data were analyzed using MAGeCK-VISPR ^76^.

### Basic data analyses

UMAP plots were created using the M3C^77^ package in R, which automatically selects the typically-user-defined UMAP plotting parameters using a Monte Carlo approach. Basic analysis of the CRISPR results was performed using MAGeCK-VISPR ^76^. The visualization of cell line genomic features was created using ProteinPaint (https://proteinpaint.stjude.org/)78. All other basic analysis and statistical tests were performed using R version 4.0.2 ^79^. False discovery rates were estimated using the Benjamini and Hochberg method.

### Hierarchical Bayesian model to identify neuroblastoma selective hits and to borrow information across mechanistically related drugs

First, gRNAs acting as outliers were filtered using a Dixon outlier test. For each gene, this test was applied to log drug vs vehicle fold-changes of normalized read counts, and misbehaving gRNAs were removed at a nominal *P*-value threshold of 0.05. We filtered single gRNAs in the cases where the directionality of that single gRNA was different from the other 5 gRNAs targeting the same gene. We also filtered gRNAs with very low (<5) read counts in both the drug and treatment groups, as these produce very unstable fold change estimates. Drug vs vehicle fold changes and their associated standard error were calculated from the remaining gRNAs.

The differential synergizing effect of a gene knockout and the potency of that effect in neuroblastoma was estimated using the following hierarchical Bayesian model:

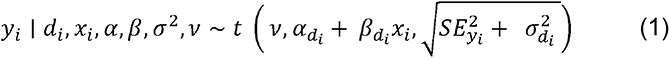

Where *Y_i_* is the normalized gRNA fold change, *d_i_* indicates the drug, *x_i_* indicates case/control status encoded as 0 or 1, α are the estimates of synergizing potency in the neuroblastoma group and β are the estimates of the differential drug synergizing effect of a knockout between the neuroblastoma cell lines and the outgroup, *v*, and σ^2^ represent the degrees-of-freedom and the variance of the t-distribution respectively. *SE*_yi_^2^ is the standard error squared associated with each fold change estimate *Y_i_*, and the model formulation above allows these error estimates to be accounted for in the estimation of this model’s parameters. Sharing of information across DNA-damaging agents was achieved using the following hierarchical structure, where the *α* and *β* parameters for each drug are assumed to be drawn from shared parameterized Normal distributions:

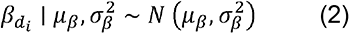

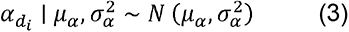

The *µ_α_* and *µ_β_* parameters can then be interpreted as the shared sensitizing effect of a gene knockout on this entire group of drugs, described for example in Fig. 4i for DNA-damaging agents. Each of these hierarchical parameters are assigned a weakly informative prior:

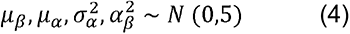

We used a gamma prior distribution for the degrees of freedom parameter of the t-distribution, with the following previously proposed shape and rate parameter values, which are suitable to implement a model reasonably robust to outliers^80^:

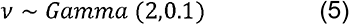

For the joint model sharing information across the 6 DNA-damaging agents and 6 × 18 = 108 cell lines, the following indices were used:

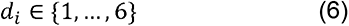

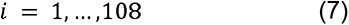

The description and equations above specifically describe the most complex model we used, jointly modeling the DNA damaging agents. In practice, we also employed several simpler iterations of this model. For example, independent models can be fit for each drug by dropping this model’s hierarchical structure (i.e. eqns. 2-4) and fitting the resulting model for each drug independently and the general drug sensitizing effect of a knockout, irrespective of neuroblastoma/outgroup status, can be estimated by dropping the model’s β parameter. The parameters were estimated using Hamiltonian Monte Carlo, implemented in the R package *rstan* (http://mc-stan.org/).

Finally, the gene set level analyses were performed by comparing the drug vs vehicle fold changes of genes in the relevant set (a pathway, a process, etc.) against a set of negative control genes. These negative control genes were created by randomly grouping the 400 non-targeting control gRNAs included in all screens into 66 negative control genes (6 gRNAs per gene). A non-parametric Wilcoxon rank sum test was then used to test the difference of the fold changes observed in each gene set against these negative control genes, which provides an estimate of the expectation under the null.

### Cisplatin-DNA adduct assay

Cells were treated with vehicle (normal saline or DMSO-HCl), cisplatin in normal saline, and cisplatin in DMSO-HCl for 24 h. Genomic DNA was extracted from treated cells and blotted in nitrocellulose membrane. The membrane was baked at 80°C to immobilize blotted gDNA for 2 hrs. The baked membrane was blocked by TBST at room temperature (RT) for 30 min, followed by incubation of primary anti-cisplatin DNA adducts antibody (1:1000, Millipore) at 4°C overnight. The next day, the membrane was rinsed three times with TBST and incubated with secondary HRP-conjugated anti-rat at RT for 30 min. After rinsing three times, the membrane was imaged by Licor Odyssey XF to measure DNA adduct (Extended Data Figure 11).

### Genetic validation of sensitized candidates

Three shRNAs per gene (PRKDC, HDAC2, KEAP1, and MET), were purchased (Sigma, Key resources). Lentiviruses of each shRNA were produced at our institutional core facility, including non-targeting control. 10 or 11 cell lines were transduced by individual shRNA lentiviruses for 3 days and followed by puromycin selection to isolate the cells with the desired knockdown. Knockdown efficiency was verified by western blot analysis. We tested the knockdown efficiency of individual shRNAs against *PRKDC, HDAC2*, *KEAP1*, and *MET* to select the most efficient one (TRCN0000194985 for *PRKDC,* TRCN0000004819 for *HDAC2*, TRCN0000154657 for *KEAP1*, and TRCN0000121087 for *MET*). Knockdown cells were treated with the corresponding drugs (IC_50_ or 10 µM) for 3 days (doxorubicin/PRKDC knockdown, JQAD1/HDAC2 knockdown, topotecan/KEAP1 knockdown, CDDP/MET knockdown) to test whether the potency of four drugs was increased.

### Dense drug-drug synergy screening assays using high throughput robotic handling

Doxorubicin, AZD7648, cisplatin, cabozantinib, topotecan, dimethyl fumarate, JQAD1, and panobinostat were added into 384-well Perkin Elmer Culture plates (Perkin Elmer #6007688) using Labcyte Echo 555 and 655T acoustic liquid dispensers moving a total volume of 80 nL into each well to generate desired combinations with an equivalent final DMSO concentration of 0.2%. Each assay plate contained ten-point dose-response curve with 1:3 dilution intervals for each compound, three replicates of the pairwise drug combination matrices, three replicates of each compound alone, and twelve replicates of each control (DMSO and 20 µM staurosporine to represent 0% and 100% cell death, respectively). Seeding densities were determined *a priori* by growth assays for each of the 18 cell lines. Cell lines were plated into assay plates in volumes of 40 μL using a Multidrop Combi reagent dispenser to reach desired the final desired concentrations, and then settled for 20 seconds at 200 × g in a Sorvall Legend XTR centrifuge. Plates were then incubated at 37 °C in 5% CO2 for 72 hours in a High Resolution Biosolutions Steristore incubator. After incubation the plates were moved to a Liconic STX220 incubator, at 37 °C in 5% CO2, integrated onta an Agilent BioCell, to determine viability. The plates were each placed at room temperature for 20 minutes before viability assessment. 25 μL of CellTiter-Glo reagent (Promega #G9241) was added to each well using a Multidrop Combi to measure viability, and the plates were incubated for an additional 20 minutes at room temperature. Luminescence was then measured with an EnVision 2102 Multilabel Plate Reader.

### Analysis of dense drug-drug synergy screening data

The raw luminescent data was imported into R. Background-subtracted values in RLU were assigned to the appropriate drugs and concentrations. All replicates were normalized to the mean of their respective inter-plate controls (vehicle for 0% cell death and staurosporine for 100% cell death). Normalized drug-only data were fit with log-logistic regression to produce dose-response curves using the DR4PL package^81, 82^. Matrices of the percent cell death values were constructed using means of normalized data from each of the replicates per treatment combination as input. From these normalized values, synergy scores were calculated for all tested concentration combinations, using the Zero Interaction Potency (ZIP) model implemented using the *SynergyFinder* package in R. The resulting synergy matrices were used to extract the highest- and lowest-scoring concentration pairs to represent the most significant synergy and antagonism.

### AZD7648 (PRKDCi) and Doxorubicin treatment in SJNB14-PDX and BE2C Xenograft models

Doxorubicin was purchased from Selleckchem (Selleckchem, S1208). AZD7648 were purchased from Chemietek (Chemietek, CT-A7648) and formulated in 0.5% hydroxypropyl methylcellulose (Sigma, H3785-100)/ 0.1% Tween80 (Sigma, P4780-500mL) (HPMC/T) for oral gavage. Tumor size was measured with electronic calipers. The tumor volume was calculated using the formula π/6 × d^3^, where d is the mean of two diameters taken at right angles. The tumor volume and mice weight were measured twice a week. When tumor size reached up to ∼100-200 mm^3^, the animals were randomized into four groups (n=4-5 mice per group). Mice were treated with vehicle (HPMC/T), doxorubicin (0.75 mg/kg, intraperitoneal, twice weekly), and AZD7648 (50 mg/kg, twice/day, oral gavage every day and the time between the morning and evening doses was 8 h) and combination of doxorubicin and AZD7648 for two weeks. On the day of doxorubicin treatment, doxorubicin was dosed 1 h after the morning dosing of AZD7648. Mice were euthanized when tumor size reached over 20% of body weight or became morbid.

### Western blot analysis

Cells were treated with vehicle (DMSO) or drug for 72hrs. Total proteins were extracted by using a modified RIAP buffer (HEPES, NaCl, EDTA, PI cocktail tablet, PPi cocktail tablet, PMSF, DTT). Total 30 μg of proteins were resolved at gradient gels (Biorad). Resolved proteins were transferred to nitrocellulose membrane by using iBlot (Invitrogen). The membrane was blocked by TBST at room temperature (RT) for 30 min, followed by primary antibodies (1:1000) at 4°C for overnight. The next day, the membrane was rinsed three times with TBST and incubated with secondary antibodies at RT for 1 hr. After rinsing three times, the membrane was imaged by Licor Odyssey XF to measure the level of target protein levels.

### Immunofluorescent staining

72 hrs after treatment with vehicle (DMSO) or drug, the cells were fixed with 4% paraformaldehyde at RT for 10 min and rinsed with 1x PBS three times. The rinsed cells were permeabilized with 0.1% Triton-X 100 in 1x PBS at RT for 10min, blocked with 0.5% fetal bovine serum, 0.01% Triton-X 100 in 1x PBS at RT for 30 min. After blocking, the cells were incubated with mouse anti-γH2AX or rabbit anti-phospho PRKDC (pS2056) at 4°C overnight. The next day, cells were rinsed with 1x PBS three times and incubated with anti-mouse Alexa488 or anti-rabbit Alexa594 at RT for 60 min. Fluorescent images were taken by Nikon Ti Eclipse and analyzed by CellProfiler (https://cellprofiler.org).

### Caspase 3/7 assay

Apoptosis was measured by luminescent caspase 3/7 assay (Promega). Briefly, 72 hrs after treatment with vehicle (DMSO) or drug in BE2C and GIMEN in 96 well plates, 100 µL of caspase 3/7 reagent was directly added to the cell plates. The cells were incubated in the dark at room temperature for 60 min, followed by reading luminescence using a CLARIOstar plate reader (BMG Labtech).

### Cell cycle arrest and drug treatment

BE2C and GIMEN were incubated with serum free medium for 48 hrs to arrest G0/G1. 48 hr after starvation, the cells were released by adding fresh complete growth medium. BE2C and GIMEN were treated with 100ng/mL of nocodazole (NOC) for 18 hrs to arrest G2/M phase. 18 hrs after treatment, the cells were released by washing out NOC three times and replaced with fresh complete growth medium. 24 hrs after release of each synchronization, the cells were treated with vehicle (DMSO) or drugs for 72 hrs. The treated cells were harvested at 1000RPM for 10 min, fixed with 70% ethanol at - 20C for 2 hrs, then washed with 1x PBS, and staining with propidium iodide (50ug/mL) for FACS.

### NHEJ and PRKDC-related assays

Non-homologous end joining (NHEJ) in BE2C and GIMEN neuroblastoma cell lines was visualized by using a custom cell-based kit for NHEJ (Topogen, Cat # DR5000A). Following the manufacturer’s guide, the cells were transfected with a modified GFP reporter, along with a plasmid encoding I-*Sce*I, or empty vector by using TransIT (Mirus, Cat # MIR5400) in 6-well plates. 24 hrs after incubation, the cells were trypsinized, and replated into 12-well plates for single treatment of doxorubicin, AZD7648, vehicle (DMSO), and combination treatment of doxorubicin and AZD7648. The changes in GFP of cells in 12-well plates were monitored by using IncuCyte SX5 (Sartorius) for 72 hrs.

### Comet assay

Comet assay was performed according to the manufacturer’s protocol (Abcam). Vehicle or drug treated BE2C and GIMEN cells were harvested and resuspended in cold 1x PBS at 100 cells/uL. The cells were mixed with comet agarose (1:10 volume ratio) and transferred to slides. The slides were lysed with alkaline buffer (0.3M NaOH, 1mM EDTA) at 4°C for 60 min. Slides were then subjected to electrophoresis at 35V for 30 min in alkaline buffer, then fixed with 70% cold ethanol for 5 min. After fixation, the slides were stained with Vista DNA dye (Abcam) to visualize DNA/nuclei. The images of the slides were taken by Nikon Ti Eclipse and analyzed by OpenComet tool (https://cometbio.org/)

### Data and code availability

The raw sequencing data have been deposited in GEO (GSE223991). Summarized gRNA and gene level data are included as Extended Data Tables. The code to reproduce the analyses have been deposited on Open Science Framework (https://osf.io/d9xgn/). Detail information on materials is in the “Key resources” table.

## Supporting information

Combined supplementary figures

Combined supplementary tables

## ACKNOWLEDGEMENTS

P.G. is supported by an NIGMS R35 award [R35GM138293] an R01 grant from NCI [R01CA260060]; K99/R00 [R00HG009679] from NHGRI; P.G, A.D.D, and J.Y also receive support from ALSAC. Funding for open access charge: NIH. A.D.D and the St. Jude Center for Advanced Genome Engineering is funded by the NCI P30 CA021765. JY was partly supported by American Cancer Society-Research Scholar (130421-RSG-17-071-01-TBG, J.Y.) and National Cancer Institute (1R01CA229739-01, 1R01CA266600-01A1). A.D.D is supported by funding from the NCI (K08CA245251-01A1), the Curesearch For Children’s Cancer Foundation, the Alex’s Lemonade Stand Foundation, Hyundai Hope on Wheels Foundation, V Foundation for Cancer Research and the Rally Foundation for Childhood Cancer Research. The content is solely the responsibility of the authors and does not necessarily represent the official views of the National Institutes of Health. The authors have declared that no conflict of interest exists.

## Notes

https://stjude.shinyapps.io/CASAVA

### Competing Interest Statement

The authors have declared no competing interest.

