## Supplementary material for "A CRISPR-drug perturbational map for identifying new compounds to combine with commonly used chemotherapeutics": Combined supplementary figures

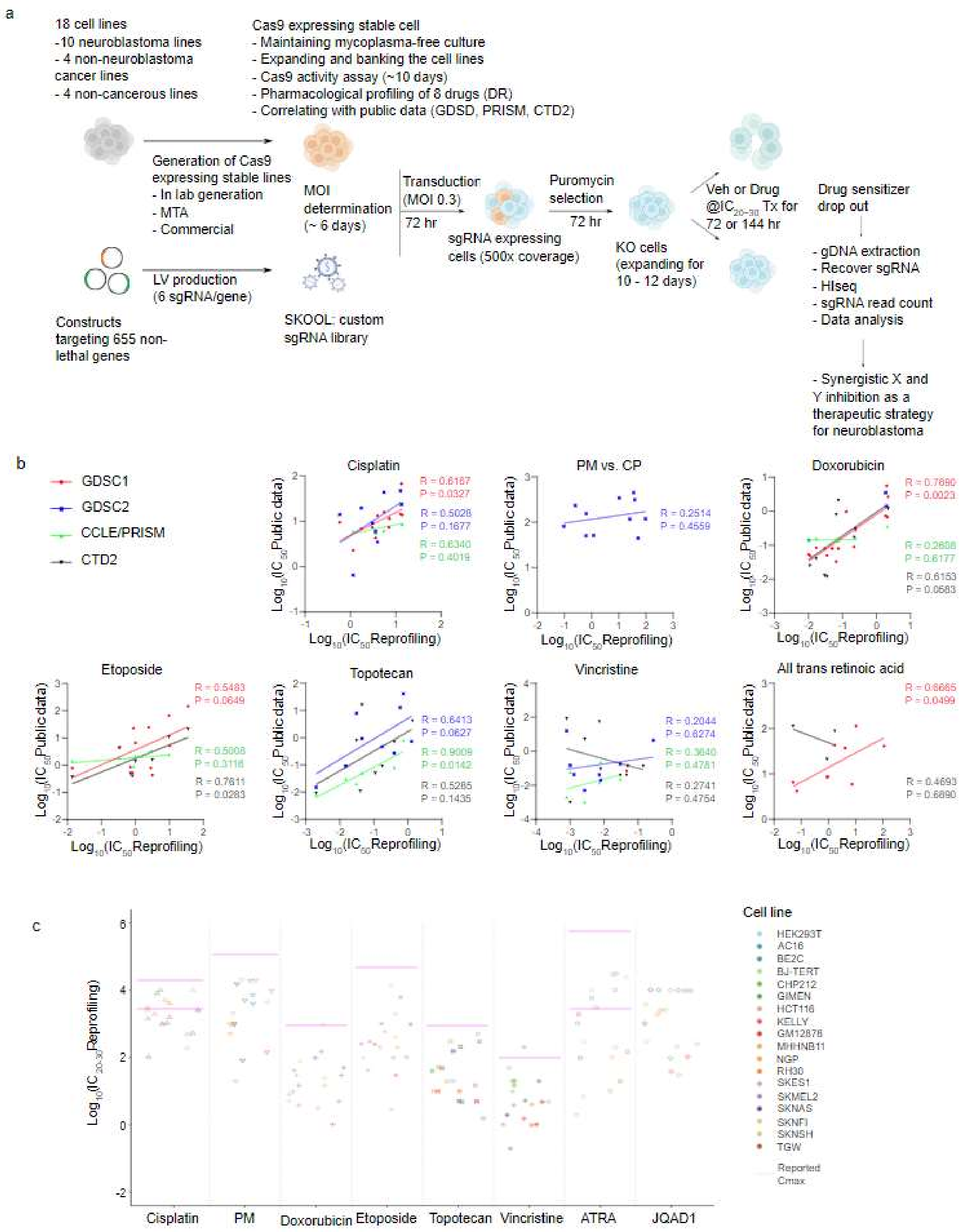


**Extended Data Figure 1. Drug screening details.**

(a) Detailed schematic of CRISPR-drug screens.

(b) Scatterplots showing the correlation of IC50 values obtained for each of our 8 drugs plotted against IC50 values reported in the public GDSC/CCLE/CTD[2] screening datasets from the Sanger/Broad Institutes.

(c) A table with all IC20-30 used across all screens (the pink lines show known Cmax values, note in resistant cells we generally capped our screening concentrations approximately at these Cmax values)

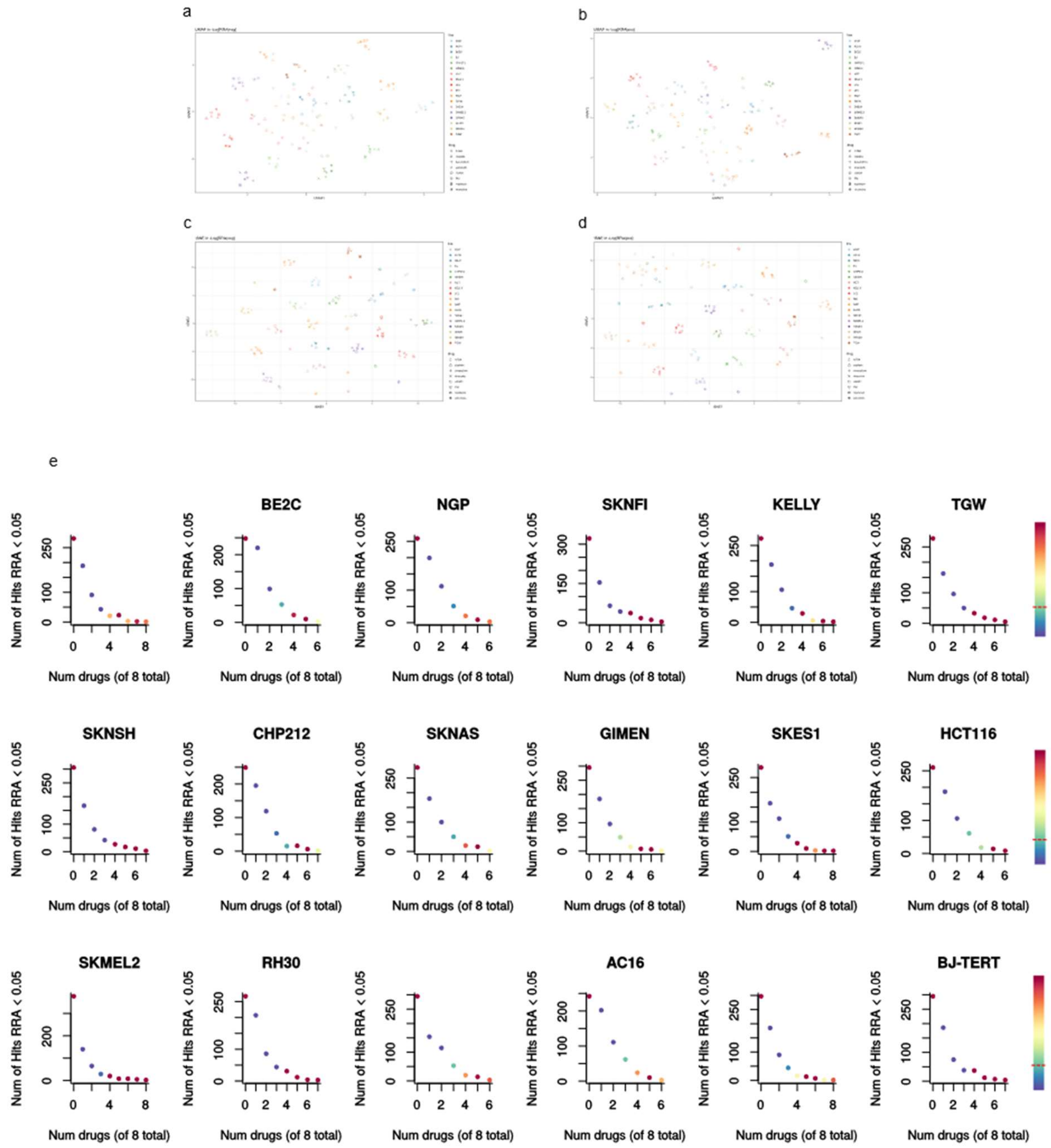


**Extended Data Figure 2: Details of systematic behaviors in screen.**

(a, b) Extra UMAP plots of all RRA scores for sensitization (a) and resistance (b) in screen.

(c, d) tSNE plots of all RRA scores for sensitization (c) and resistance (d) in screen.

(e) Scatterplot showing the number of drugs (x axis) that are sensitized to some gene knockout (RRA < 0.05; y axis) for all 18 cell lines screened. P-values (colors) were calculated by permutation.

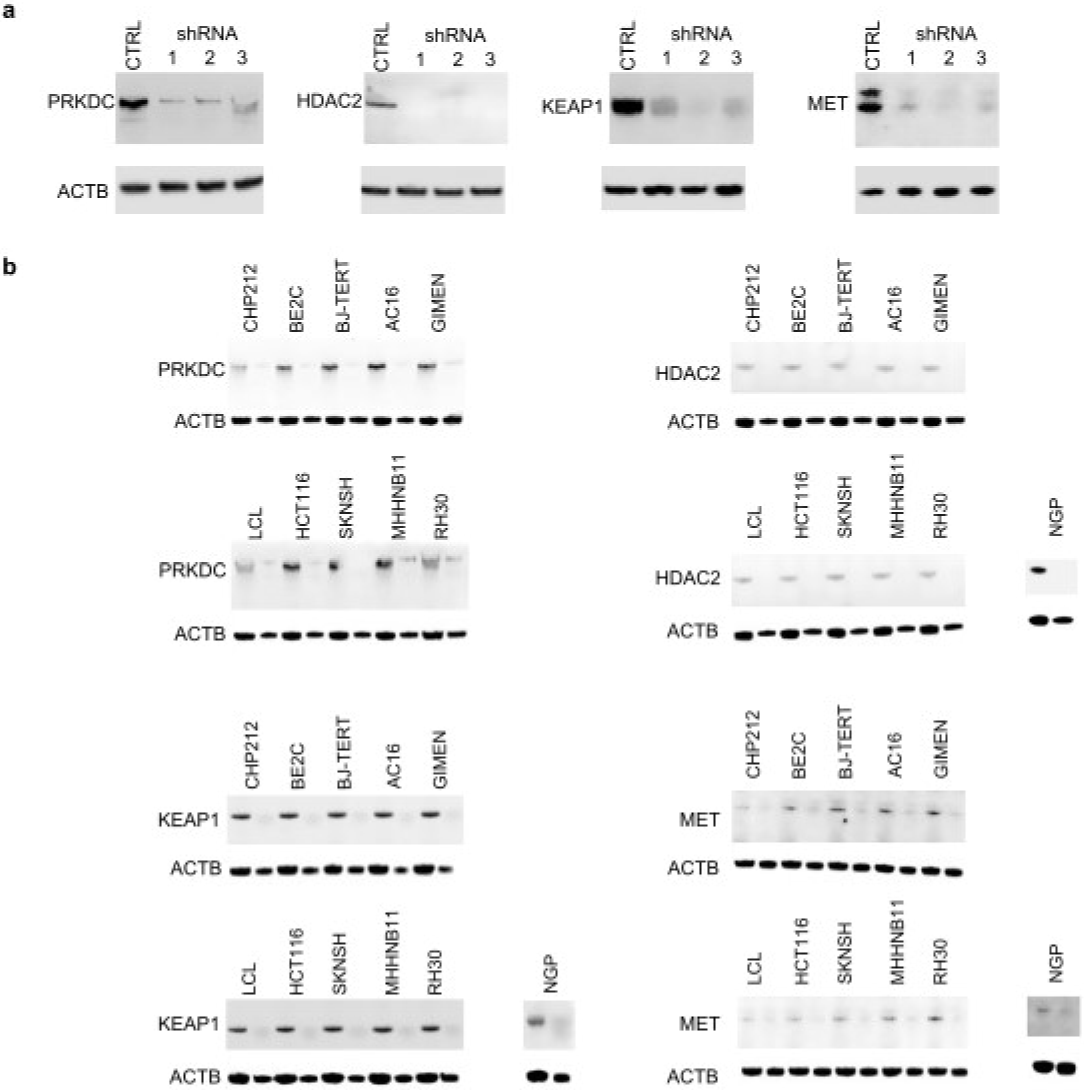

a
CTRL
shRNA
1 2 3
PRKDC
ACTB
HDAC2
KEAP1
MET
b
CHP212
BE2C
BJ-TERT
AC16
GIMEN
LCL
HCT116
SKNSH
MHHNB11
RH30
NGP

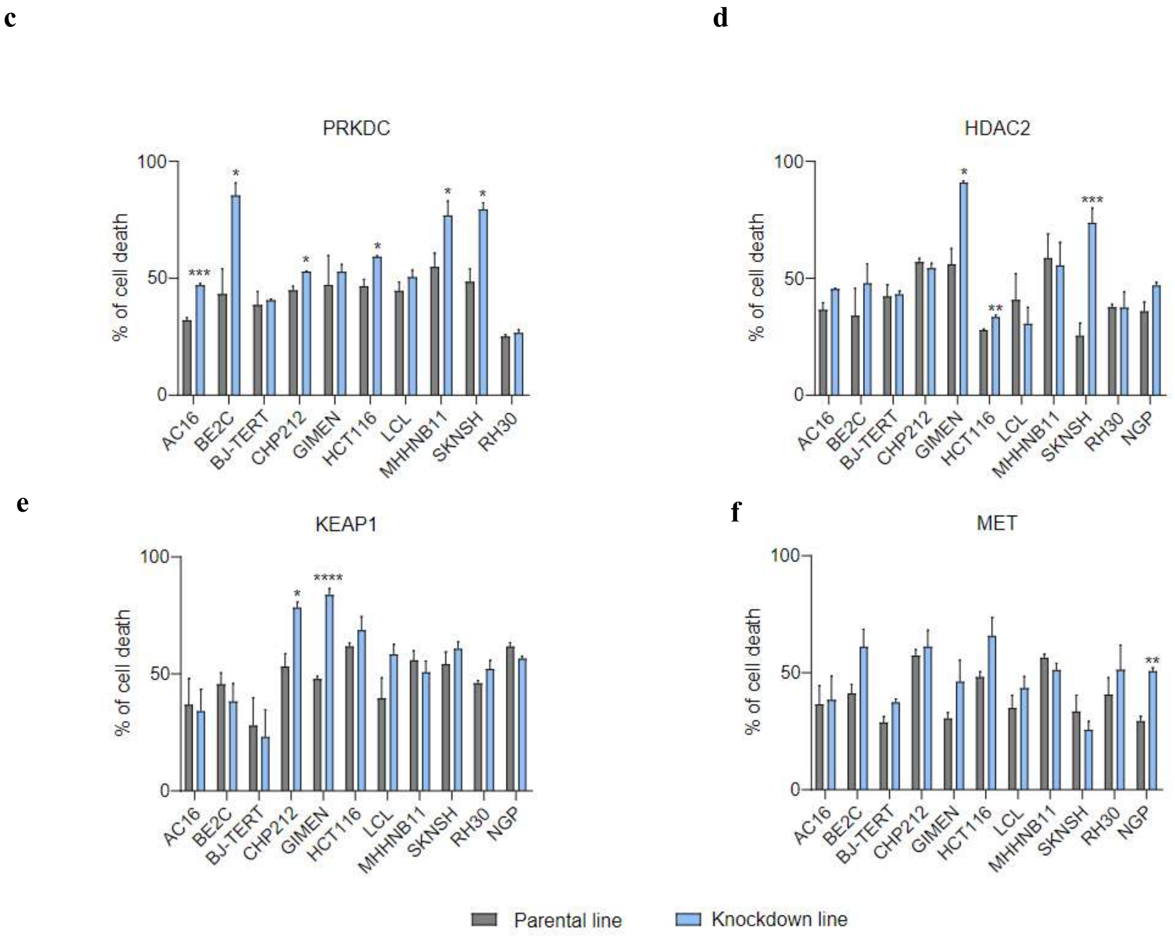


**Extended Data Figure 3. Drug responses in selected parental and associated shRNA knockdown cell lines.**

(a) Knockdown efficiency of individual shRNAs per gene. shPRKDC #1, shHDAC2 #1, shKEAP1 #2, and shMET #2 were used for subsequent drug response profiling (c), (d), (e), and (f), respectively.

(b) Knockdown of individual genes in individual cell lines. PRKDC (top left, corresponding (c)), HDAC2 (top right, corresponding to (d)), KEPA1 (bottom left, corresponding to (e)), MET (bottom right, corresponding to (f))

(c) The response of 10 parental lines (grey) and associated PRKDC knockdown cell lines (blue) to doxorubicin (~$IC_{50}$). Six knockdown cell lines were sensitized to doxorubicin, suggesting synergy (*** P < 0.001, * P < 0.05).

(d) The response of 11 parental lines (grey) and associated HDAC2 knockdown cell lines (blue) to JQAD1 (~$IC_{50}$ or capped 10μM). Three knockdown cell lines were sensitized JQAD1, suggesting synergy (*** P < 0.001, ** P < 0.01, * P < 0.05).

(e) The response of 11 parental lines (grey) and associated KEAP1 knockdown cell lines (blue) to topotecan (~$IC_{50}$). Two knockdown cell lines were sensitized to topotecan, suggesting synergy (**** P < 0.0001, * P < 0.05).

(f) The response of 11 parental lines (grey) and associated MET knockdown cell lines (blue) to cisplatin (~$IC_{50}$ or capped 10μM). One knockdown cell line were sensitized to cisplatin, suggesting synergy (** P < 0.01).

In (c-f) x-axis indicates cell lines and y-axis indicates % of cell death obtained by Cell-Titer Glo assay.

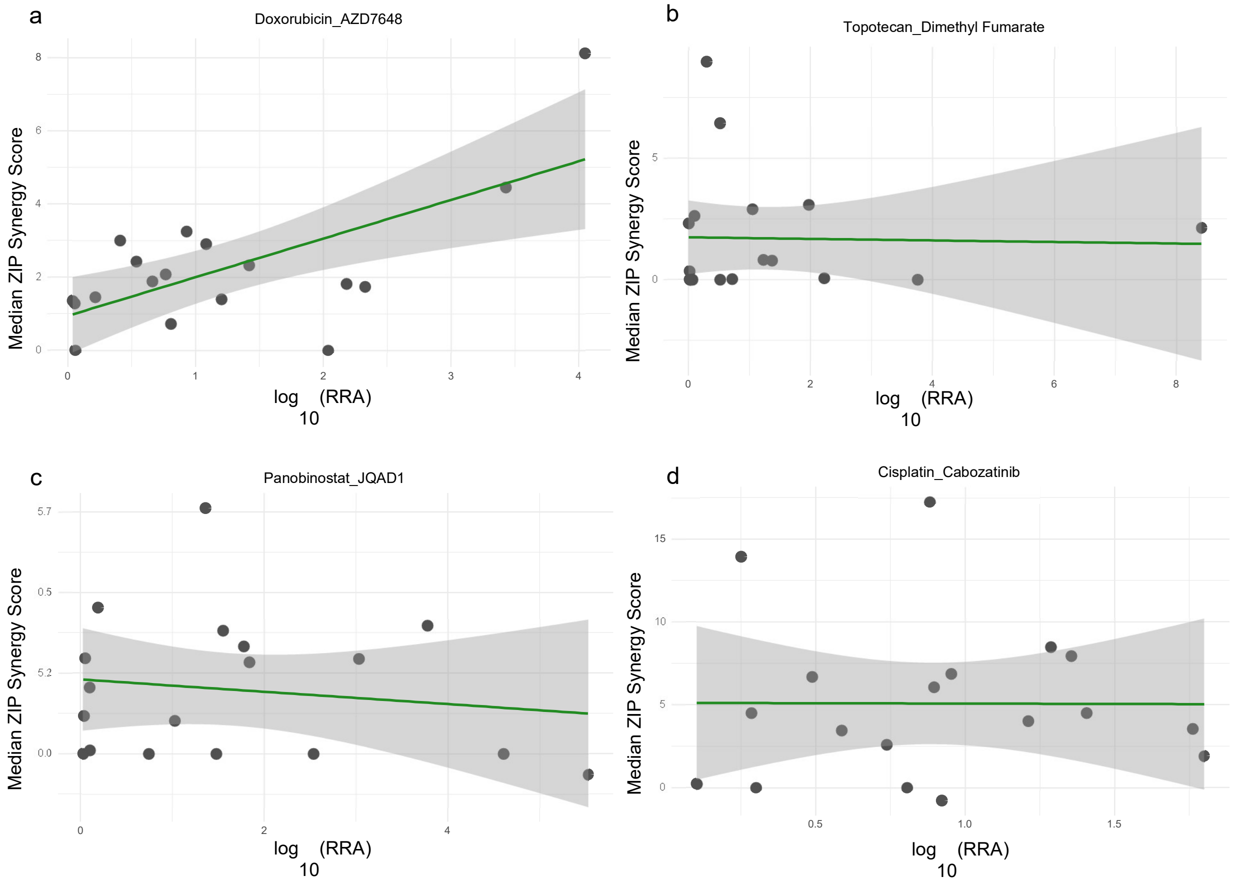


**Extended Data Figure 4. Comparison of the effect of knockout in pooled CRISPR screen to synergy scores in drug combinations**

Pearson correlation between -Log10(RRA|neg) and median ZIP in PRKDCi (a), KEAP1i (b), HDAC2i (c), and METi (d). Associated data sets are “Extended Data Table 11 for F5k and others”.

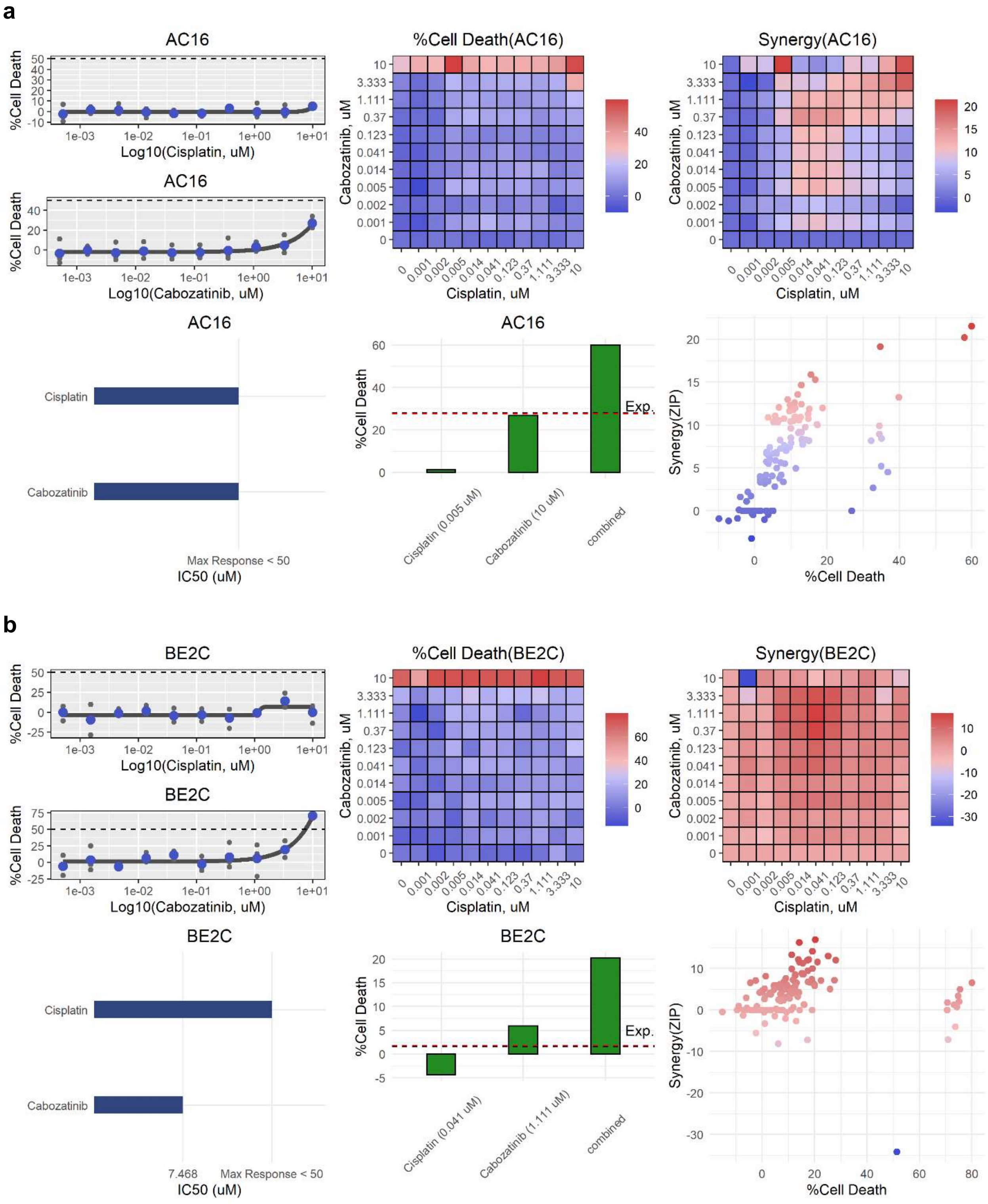
a
AC16
%Cell Death
Log10(Cisplatin, uM)
Log10(Cabozatinib, uM)
%Cell Death(AC16)
Synergy(AC16)
Cabozatinib, uM
Cisplatin, uM
Cisplatin
Cabozatinib
Max Response < 50
IC50 (uM)
Cisplatin (0.005 uM)
Cabozatinib (10 uM)
combined
Exp.
Synergy(ZIP)
b
BE2C
%Cell Death(BE2C)
Synergy(BE2C)
7.468
Cisplatin (0.041 uM)
Cabozatinib (1.111 uM)

c

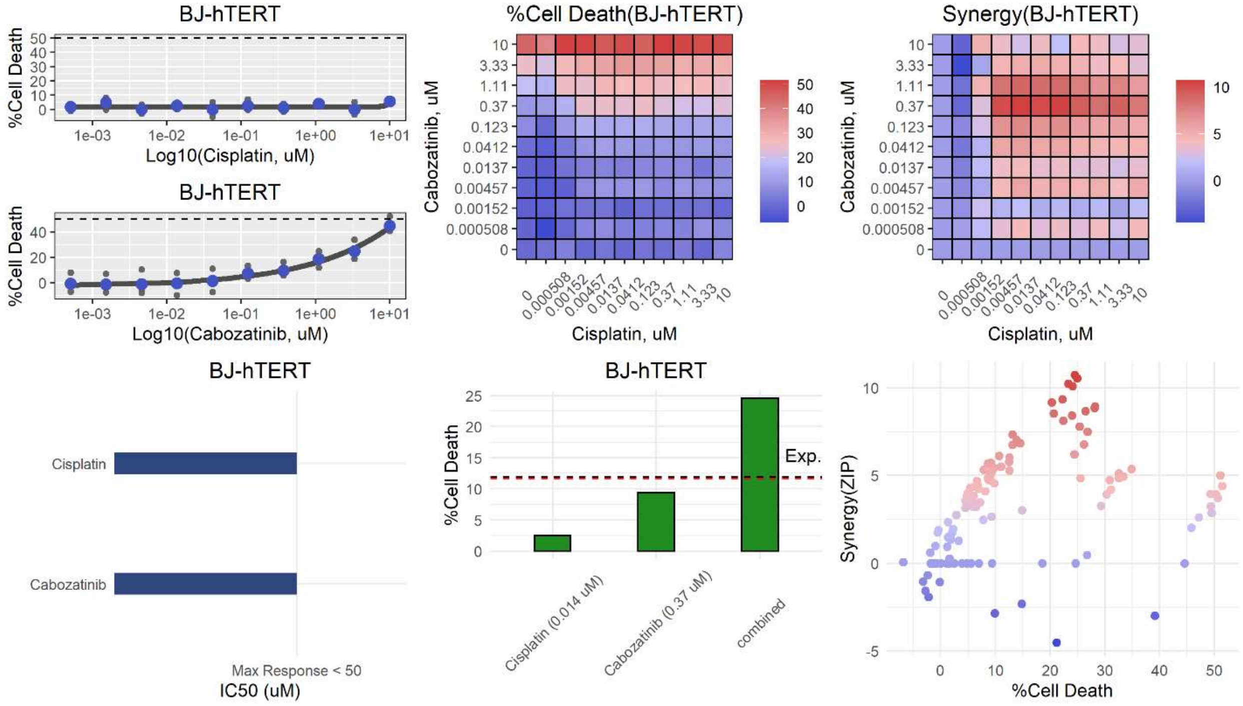
BJ-hTERT
%Cell Death
Log10(Cisplatin, uM)
1e-03
1e-02
1e-01
1e+00
1e+01
BJ-hTERT
Log10(Cabozatinib, uM)
%Cell Death(BJ-hTERT)
Cabozatinib, uM
Cisplatin, uM
10
3.33
1.11
0.37
0.123
0.0412
0.0137
0.00457
0.00152
0.000508
0
Synergy(BJ-hTERT)
BJ-hTERT
Cisplatin
Cabozatinib
Max Response < 50
IC50 (uM)
BJ-hTERT
Exp.
Cisplatin (0.014 uM)
Cabozatinib (0.37 uM)
combined
Synergy(ZIP)
%Cell Death

d

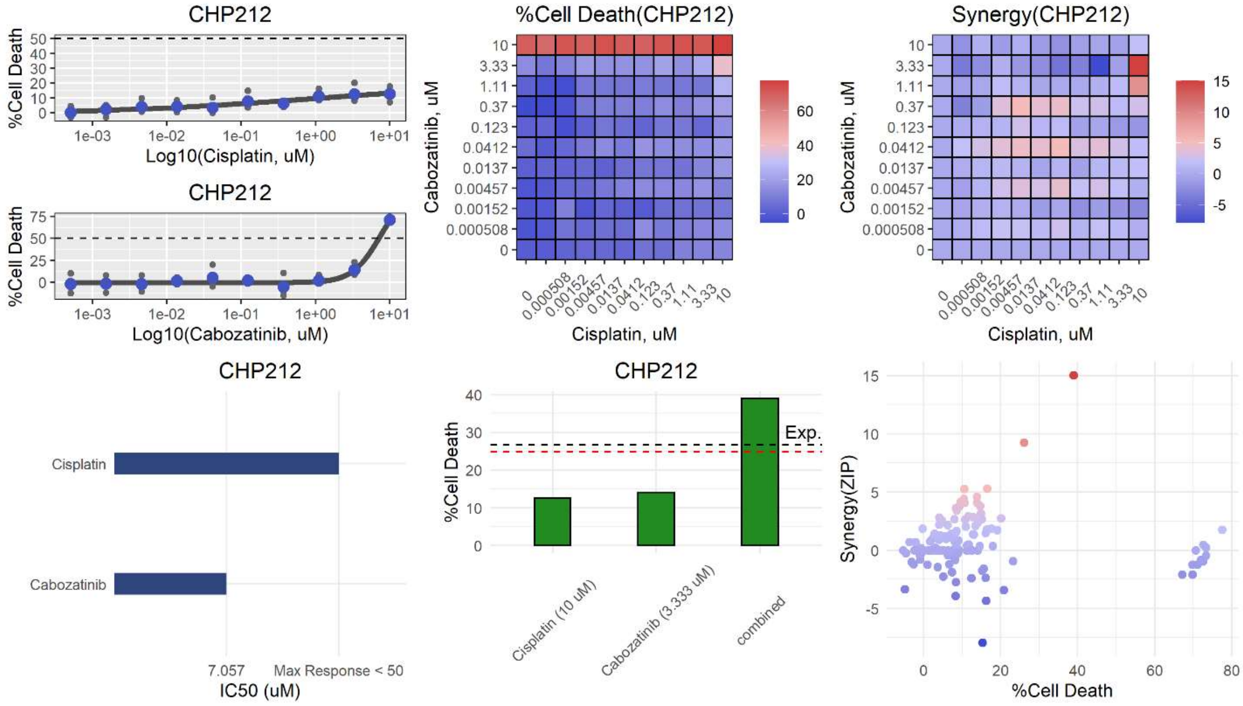
CHP212
%Cell Death
Log10(Cisplatin, uM)
1e-03
1e-02
1e-01
1e+00
1e+01
CHP212
Log10(Cabozatinib, uM)
%Cell Death(CHP212)
Cabozatinib, uM
Cisplatin, uM
10
3.33
1.11
0.37
0.123
0.0412
0.0137
0.00457
0.00152
0.000508
0
Synergy(CHP212)
CHP212
Cisplatin
Cabozatinib
7.057
Max Response < 50
IC50 (uM)
CHP212
Exp.
Cisplatin (10 uM)
Cabozatinib (3.333 uM)
combined
Synergy(ZIP)
%Cell Death

**e**

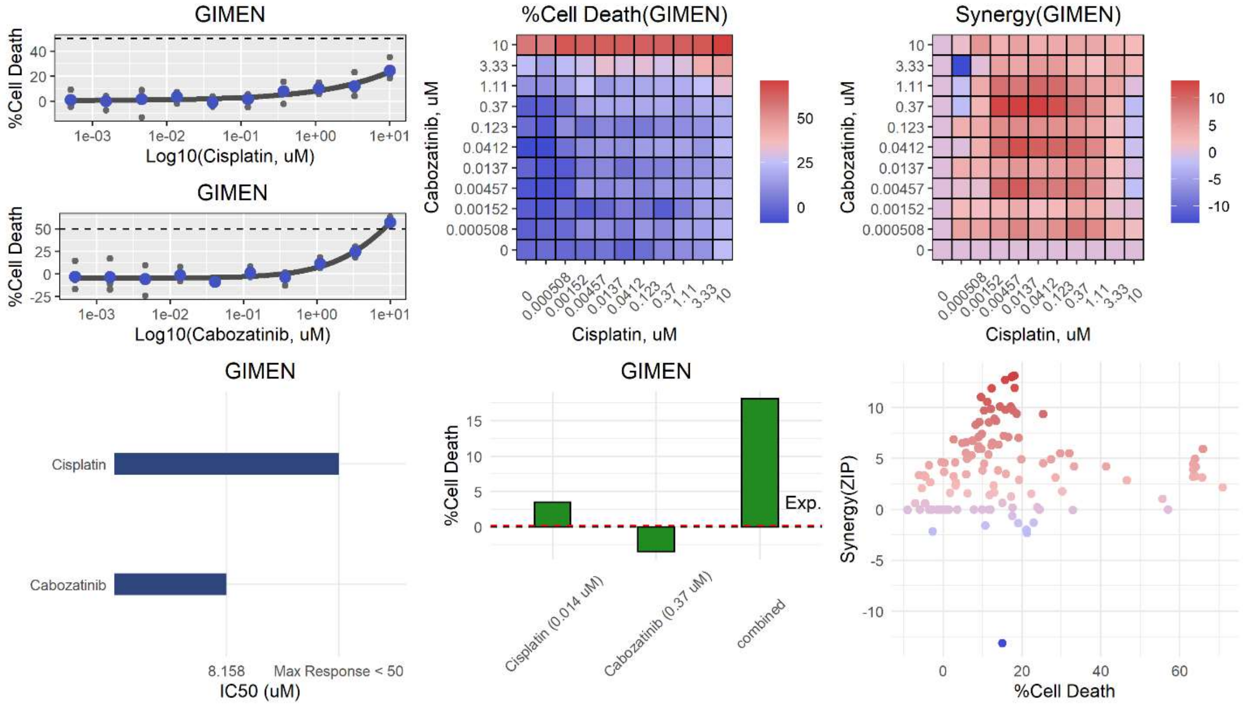
GIMEN
%Cell Death
Log10(Cisplatin, uM)
GIMEN
%Cell Death
Log10(Cabozatinib, uM)
%Cell Death(GIMEN)
Cabozatinib, uM
Cisplatin, uM
Synergy(GIMEN)
Cabozatinib, uM
Cisplatin, uM
GIMEN
Cisplatin
Cabozatinib
8.158
Max Response < 50
IC50 (uM)
GIMEN
%Cell Death
Exp.
Cisplatin (0.014 uM)
Cabozatinib (0.37 uM)
combined
Synergy(ZIP)
%Cell Death

**f**

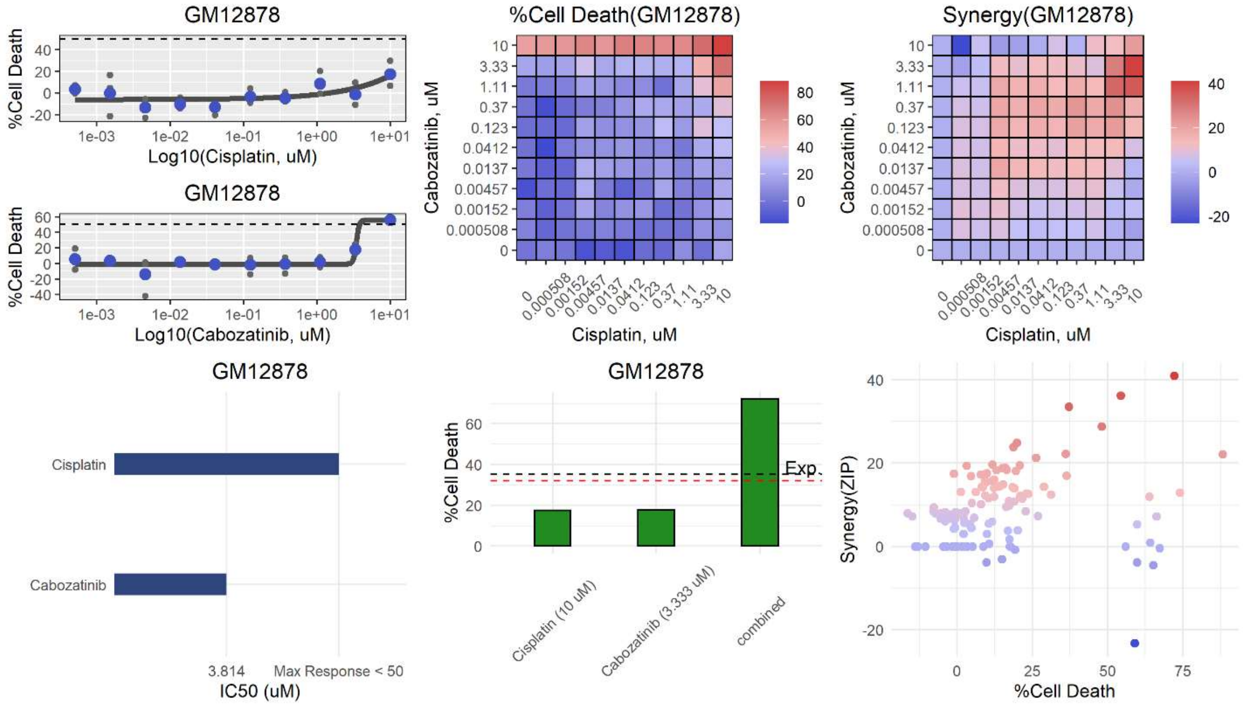
GM12878
%Cell Death
Log10(Cisplatin, uM)
GM12878
%Cell Death
Log10(Cabozatinib, uM)
%Cell Death(GM12878)
Cabozatinib, uM
Cisplatin, uM
Synergy(GM12878)
Cabozatinib, uM
Cisplatin, uM
GM12878
Cisplatin
Cabozatinib
3.814
Max Response < 50
IC50 (uM)
GM12878
%Cell Death
Exp.
Cisplatin (10 uM)
Cabozatinib (3.333 uM)
combined
Synergy(ZIP)
%Cell Death

**g**

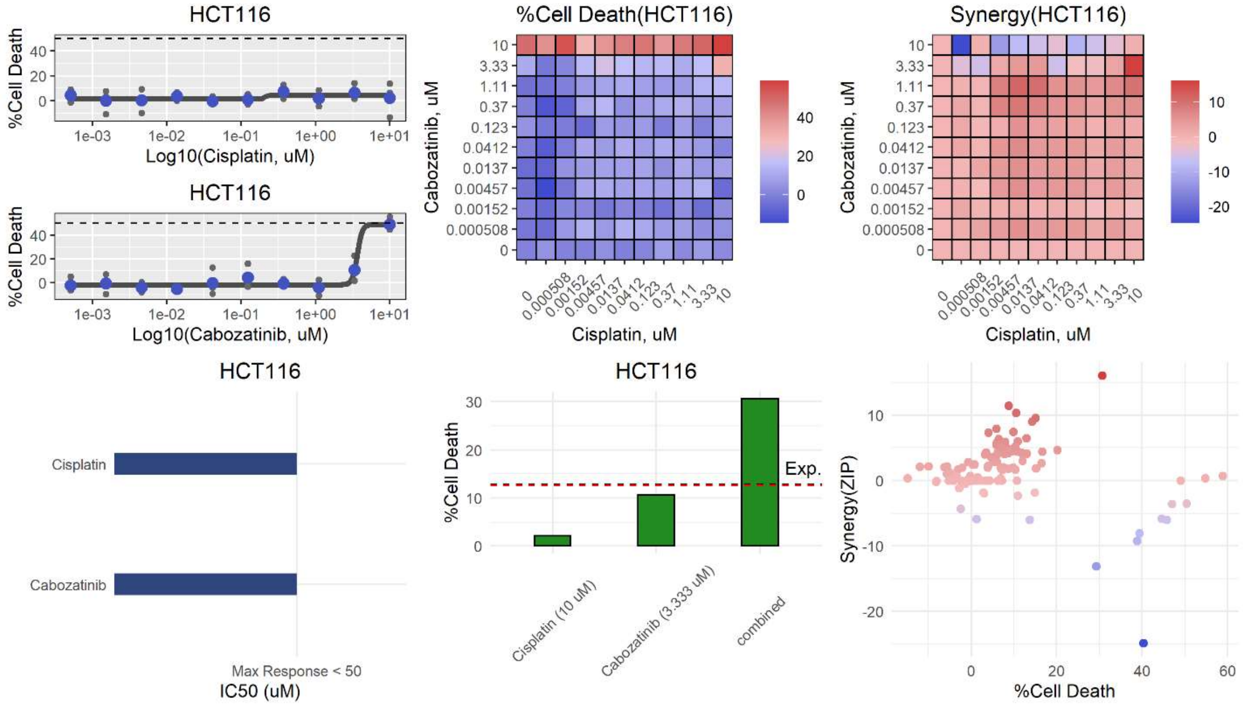
HCT116
%Cell Death
Log10(Cisplatin, uM)
HCT116
Log10(Cabozatinib, uM)
%Cell Death(HCT116)
Cabozatinib, uM
Cisplatin, uM
Synergy(HCT116)
HCT116
Cisplatin
Cabozatinib
Max Response < 50
IC50 (uM)
HCT116
Exp.
Cisplatin (10 uM)
Cabozatinib (3.333 uM)
combined
Synergy(ZIP)
%Cell Death

**h**

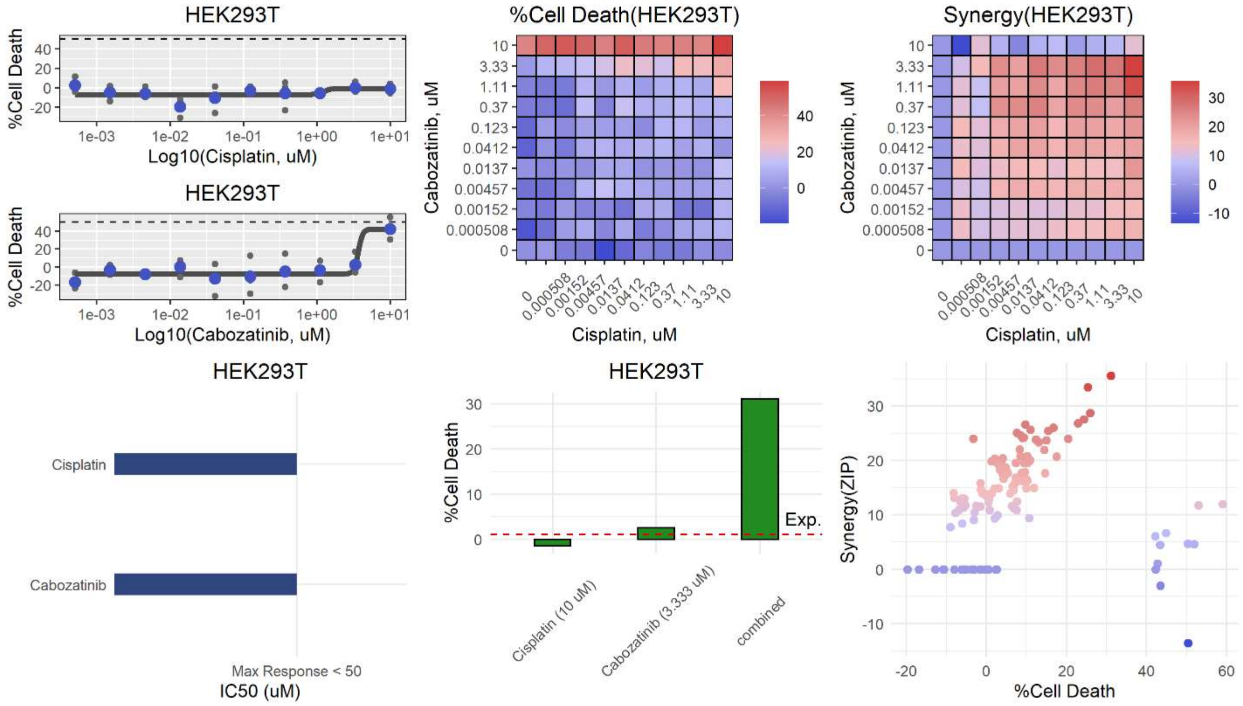
HEK293T
%Cell Death
Log10(Cisplatin, uM)
HEK293T
Log10(Cabozatinib, uM)
%Cell Death(HEK293T)
Cabozatinib, uM
Cisplatin, uM
Synergy(HEK293T)
HEK293T
Cisplatin
Cabozatinib
Max Response < 50
IC50 (uM)
HEK293T
Exp.
Cisplatin (10 uM)
Cabozatinib (3.333 uM)
combined
Synergy(ZIP)
%Cell Death

**i**

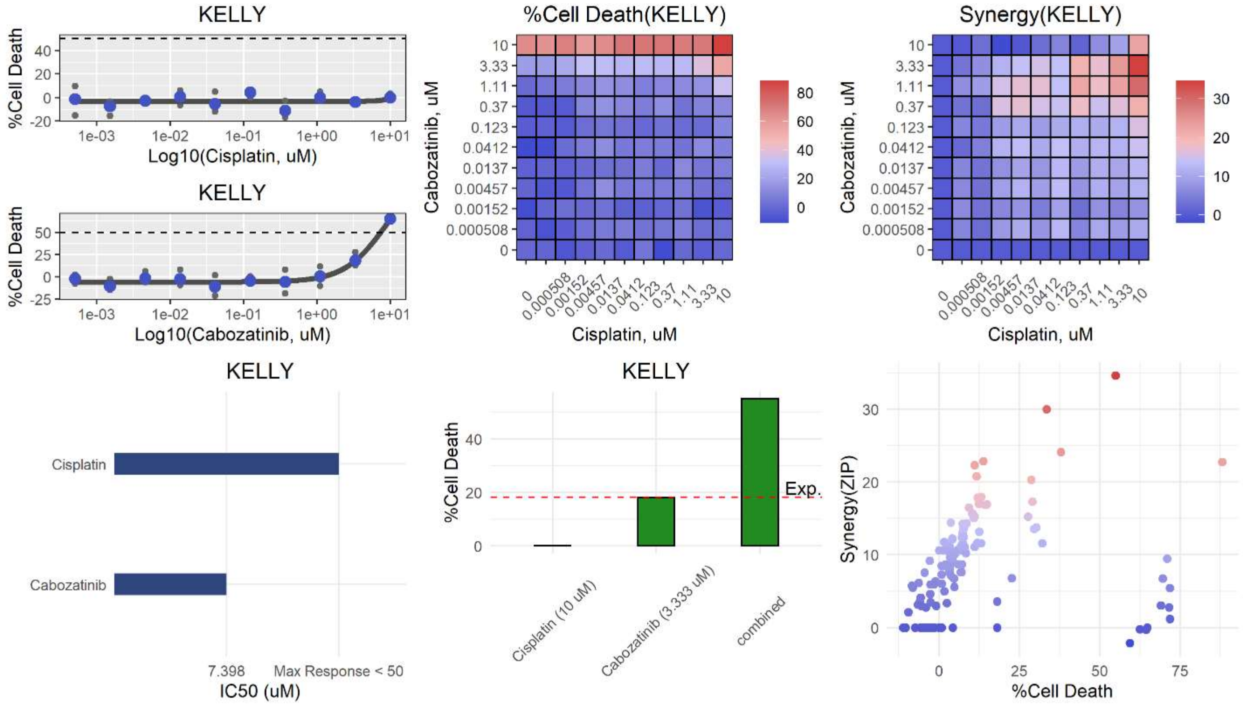
KELLY
%Cell Death
Log10(Cisplatin, uM)
KELLY
%Cell Death
Log10(Cabozatinib, uM)
%Cell Death(KELLY)
Cabozatinib, uM
Cisplatin, uM
Synergy(KELLY)
Cabozatinib, uM
Cisplatin, uM
KELLY
Cisplatin
Cabozatinib
7.398
Max Response < 50
IC50 (uM)
KELLY
%Cell Death
Exp.
Cisplatin (10 uM)
Cabozatinib (3.333 uM)
combined
Synergy(ZIP)
%Cell Death

**j**

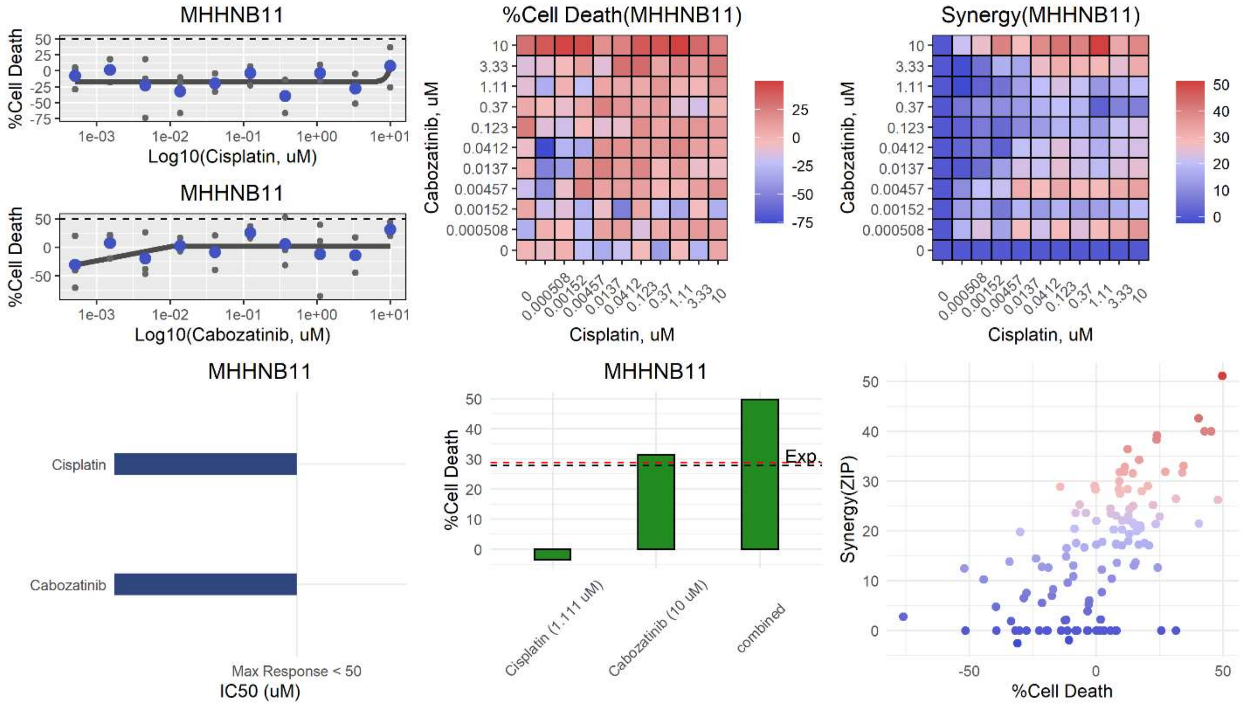
MHHNB11
%Cell Death
Log10(Cisplatin, uM)
MHHNB11
%Cell Death
Log10(Cabozatinib, uM)
%Cell Death(MHHNB11)
Cabozatinib, uM
Cisplatin, uM
Synergy(MHHNB11)
Cabozatinib, uM
Cisplatin, uM
MHHNB11
Cisplatin
Cabozatinib
Max Response < 50
IC50 (uM)
MHHNB11
%Cell Death
Exp.
Cisplatin (1.111 uM)
Cabozatinib (10 uM)
combined
Synergy(ZIP)
%Cell Death

**k**

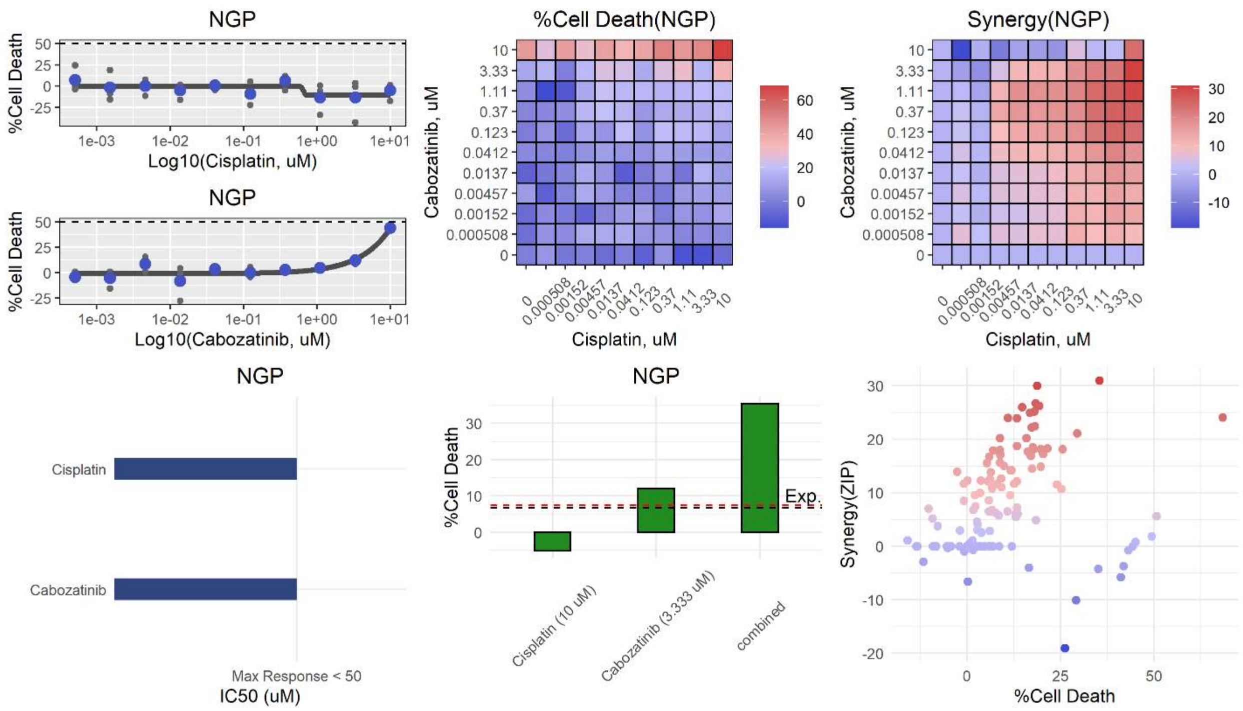
NGP
%Cell Death
Log10(Cisplatin, uM)
NGP
%Cell Death
Log10(Cabozatinib, uM)
%Cell Death(NGP)
Cabozatinib, uM
Cisplatin, uM
Synergy(NGP)
Cabozatinib, uM
Cisplatin, uM
NGP
Cisplatin
Cabozatinib
Max Response < 50
IC50 (uM)
NGP
%Cell Death
Exp.
Cisplatin (10 uM)
Cabozatinib (3.333 uM)
combined
Synergy(ZIP)
%Cell Death

**l**

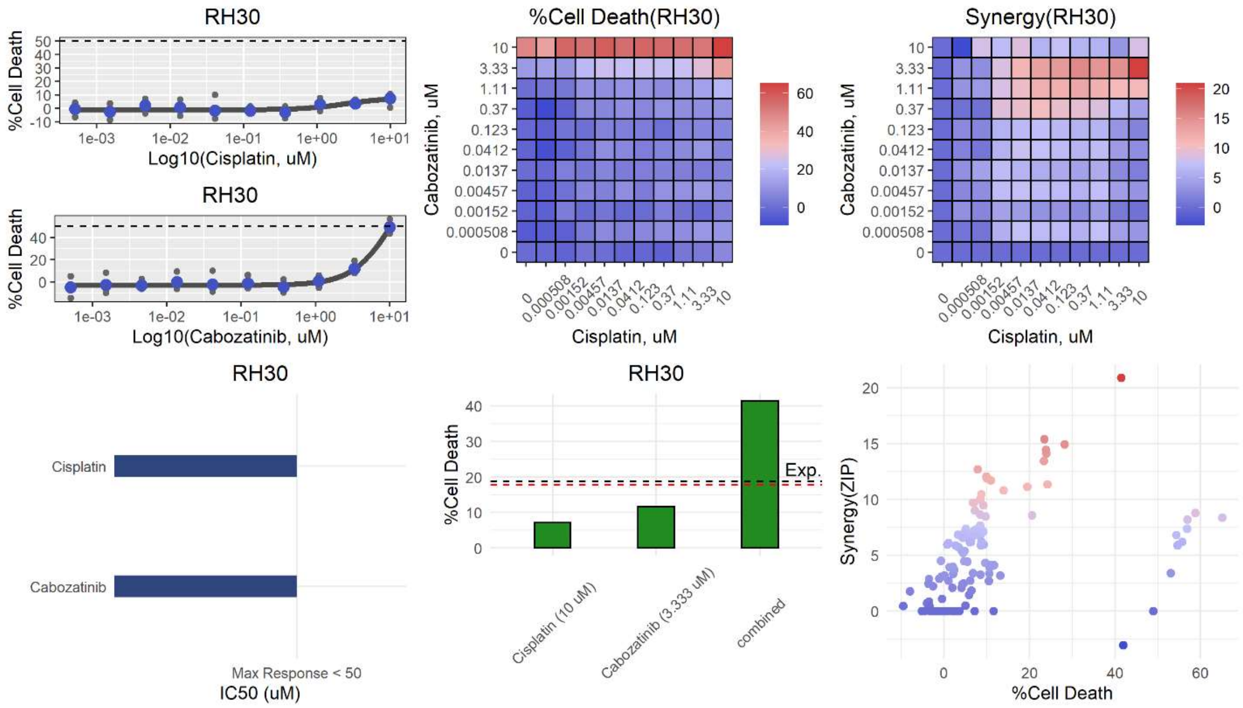
RH30
%Cell Death
Log10(Cisplatin, uM)
RH30
%Cell Death
Log10(Cabozatinib, uM)
%Cell Death(RH30)
Cabozatinib, uM
Cisplatin, uM
Synergy(RH30)
Cabozatinib, uM
Cisplatin, uM
RH30
Cisplatin
Cabozatinib
Max Response < 50
IC50 (uM)
RH30
%Cell Death
Exp.
Cisplatin (10 uM)
Cabozatinib (3.333 uM)
combined
Synergy(ZIP)
%Cell Death

**m**

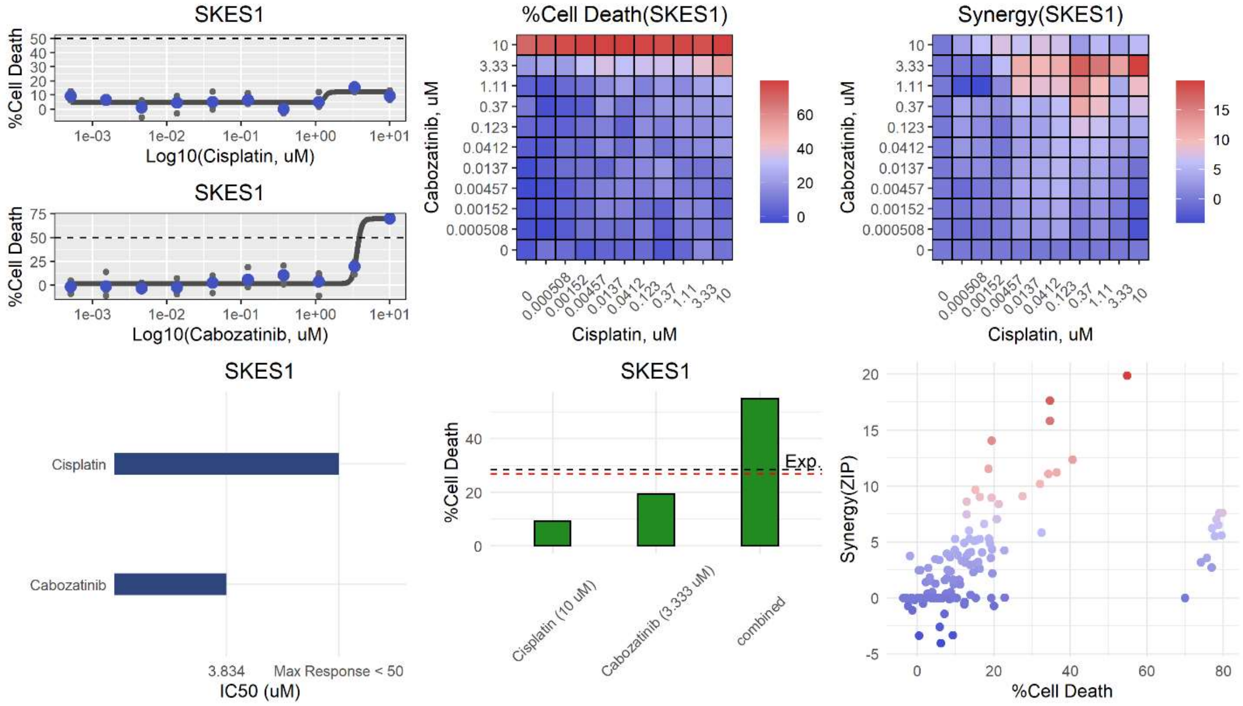

SKES1
%Cell Death
Log10(Cisplatin, uM)
SKES1
%Cell Death
Log10(Cabozatinib, uM)
%Cell Death(SKES1)
Cabozatinib, uM
Cisplatin, uM
Synergy(SKES1)
Cabozatinib, uM
Cisplatin, uM
SKES1
Cisplatin
Cabozatinib
3.834
Max Response < 50
IC50 (uM)
SKES1
%Cell Death
Exp.
Cisplatin (10 uM)
Cabozatinib (3.333 uM)
combined
Synergy(ZIP)
%Cell Death


**n**

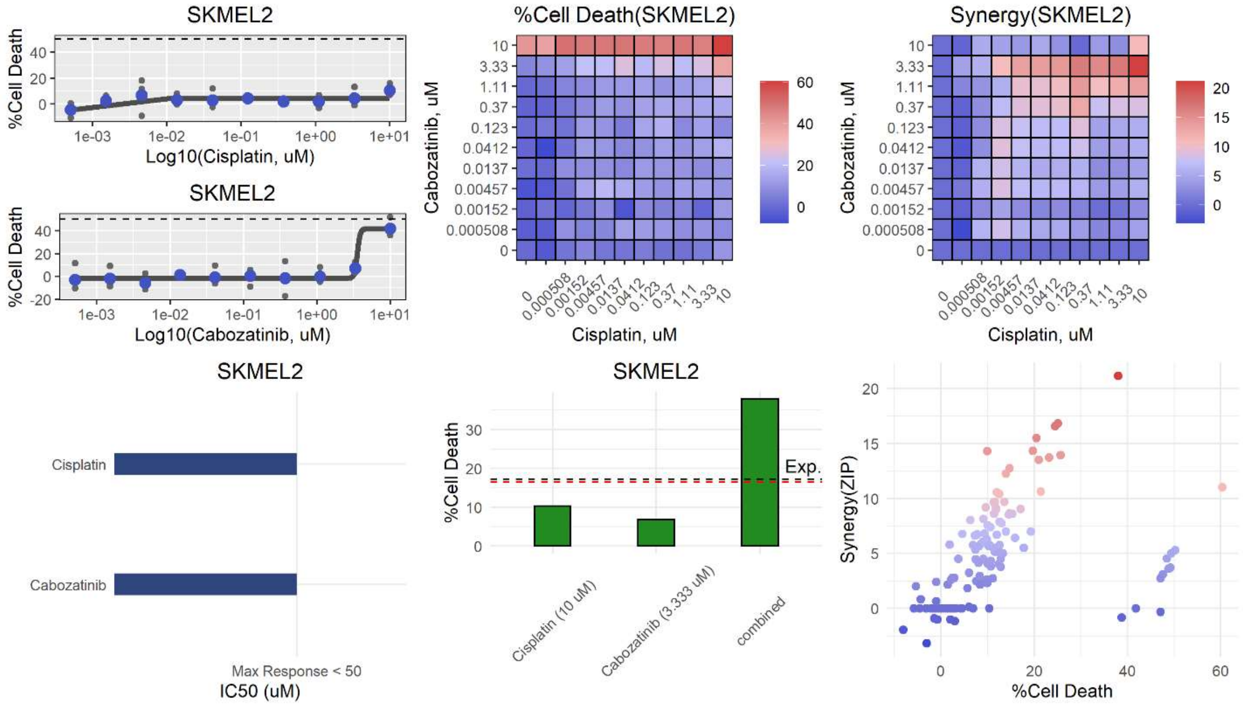

SKMEL2
%Cell Death
Log10(Cisplatin, uM)
SKMEL2
%Cell Death
Log10(Cabozatinib, uM)
%Cell Death(SKMEL2)
Cabozatinib, uM
Cisplatin, uM
Synergy(SKMEL2)
Cabozatinib, uM
Cisplatin, uM
SKMEL2
Cisplatin
Cabozatinib
Max Response < 50
IC50 (uM)
SKMEL2
%Cell Death
Exp.
Cisplatin (10 uM)
Cabozatinib (3.333 uM)
combined
Synergy(ZIP)
%Cell Death

**o**

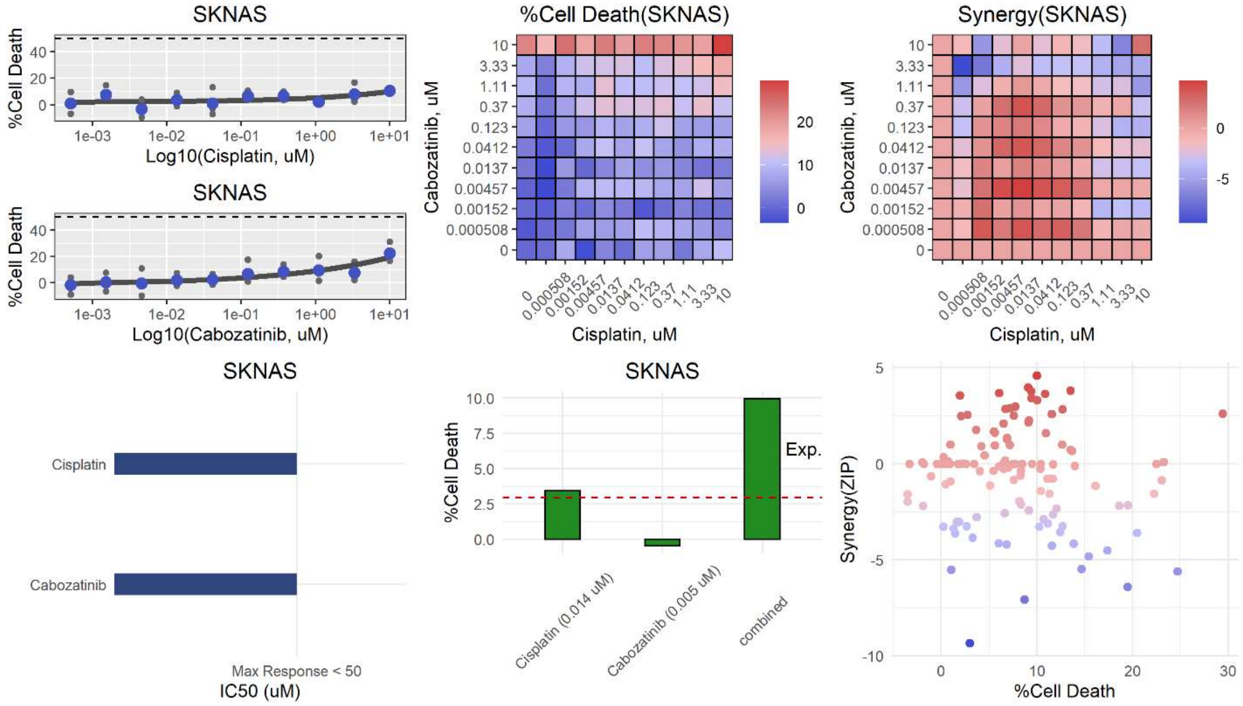
SKNAS
%Cell Death
Log10(Cisplatin, uM)
Log10(Cabozatinib, uM)
%Cell Death(SKNAS)
Cabozatinib, uM
Cisplatin, uM
Synergy(SKNAS)
Cisplatin
Cabozatinib
Max Response < 50
IC50 (uM)
Cisplatin (0.014 uM)
Cabozatinib (0.005 uM)
combined
Exp.
Synergy(ZIP)
%Cell Death

**p**

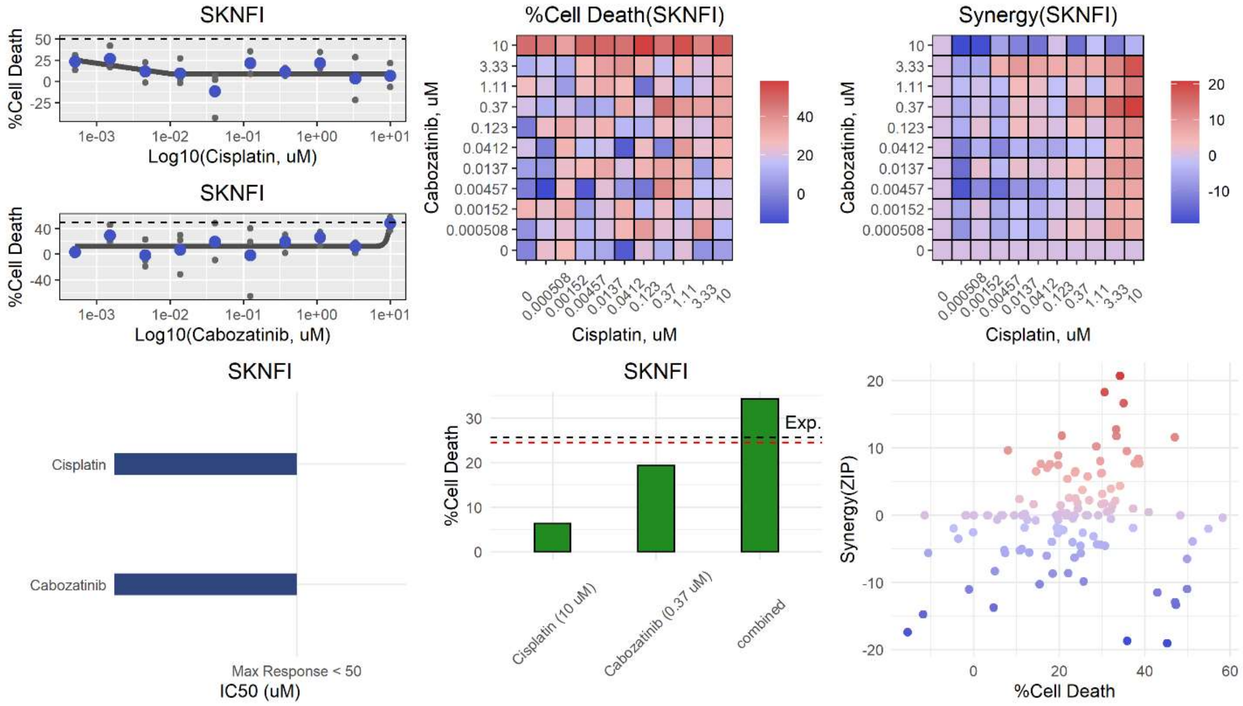
SKNFI
%Cell Death
Log10(Cisplatin, uM)
Log10(Cabozatinib, uM)
%Cell Death(SKNFI)
Cabozatinib, uM
Cisplatin, uM
Synergy(SKNFI)
Cisplatin
Cabozatinib
Max Response < 50
IC50 (uM)
Cisplatin (10 uM)
Cabozatinib (0.37 uM)
combined
Exp.
Synergy(ZIP)
%Cell Death

q

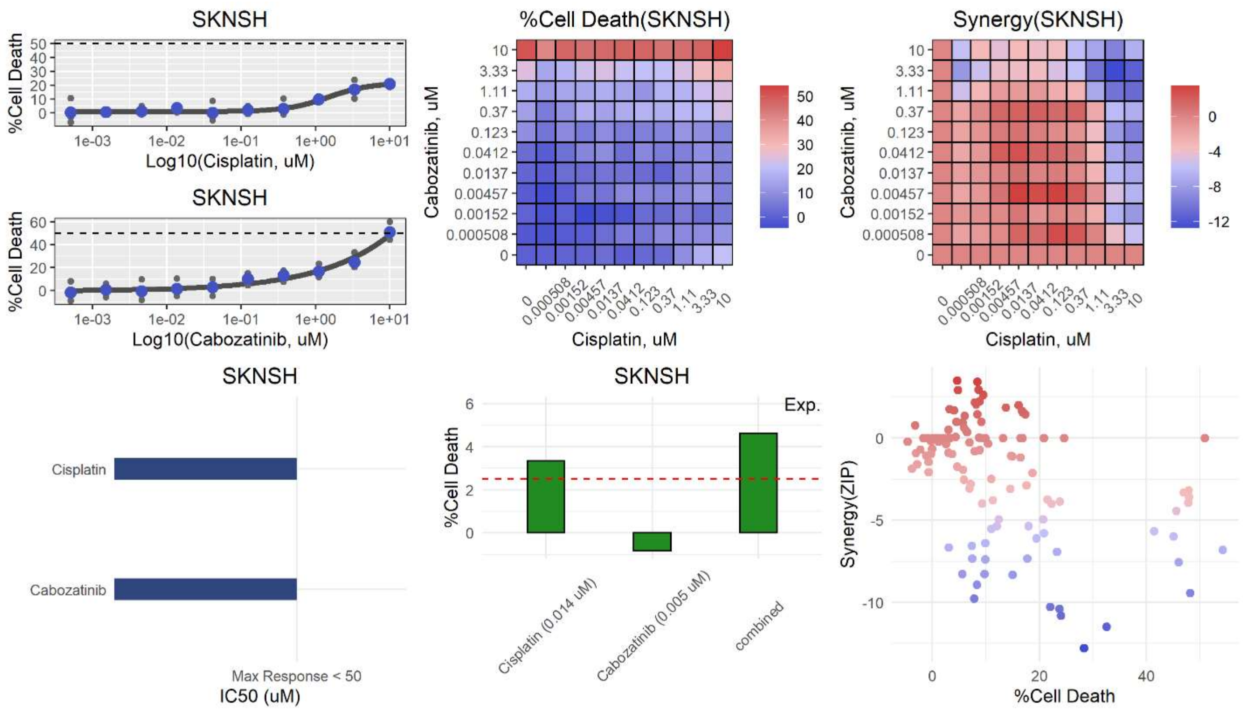
SKNSH
%Cell Death
Log10(Cisplatin, uM)
SKNSH
%Cell Death
Log10(Cabozatinib, uM)
%Cell Death(SKNSH)
Cabozatinib, uM
Cisplatin, uM
Synergy(SKNSH)
Cabozatinib, uM
Cisplatin, uM
SKNSH
Cisplatin
Cabozatinib
Max Response < 50
IC50 (uM)
SKNSH
%Cell Death
Exp.
Cisplatin (0.014 uM)
Cabozatinib (0.005 uM)
combined
Synergy(ZIP)
%Cell Death

r

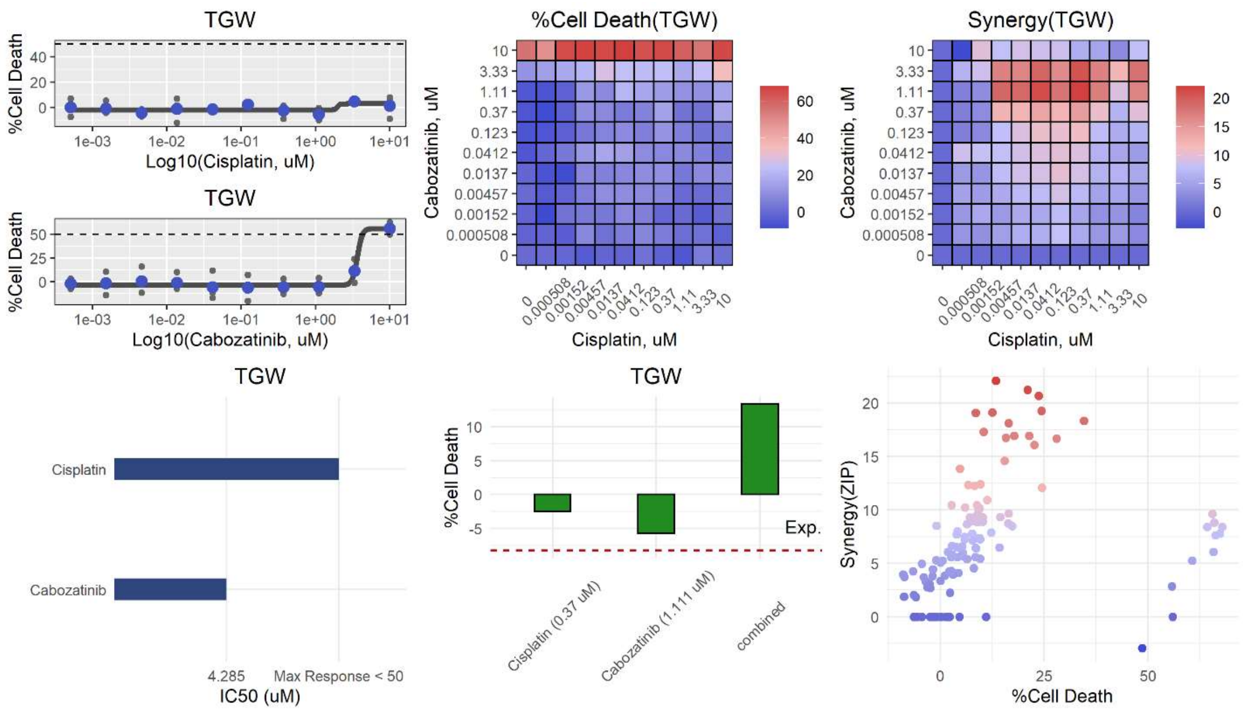
TGW
%Cell Death
Log10(Cisplatin, uM)
TGW
%Cell Death
Log10(Cabozatinib, uM)
%Cell Death(TGW)
Cabozatinib, uM
Cisplatin, uM
Synergy(TGW)
Cabozatinib, uM
Cisplatin, uM
TGW
Cisplatin
Cabozatinib
4.285
Max Response < 50
IC50 (uM)
TGW
%Cell Death
Exp.
Cisplatin (0.37 uM)
Cabozatinib (1.111 uM)
combined
Synergy(ZIP)
%Cell Death

**s**

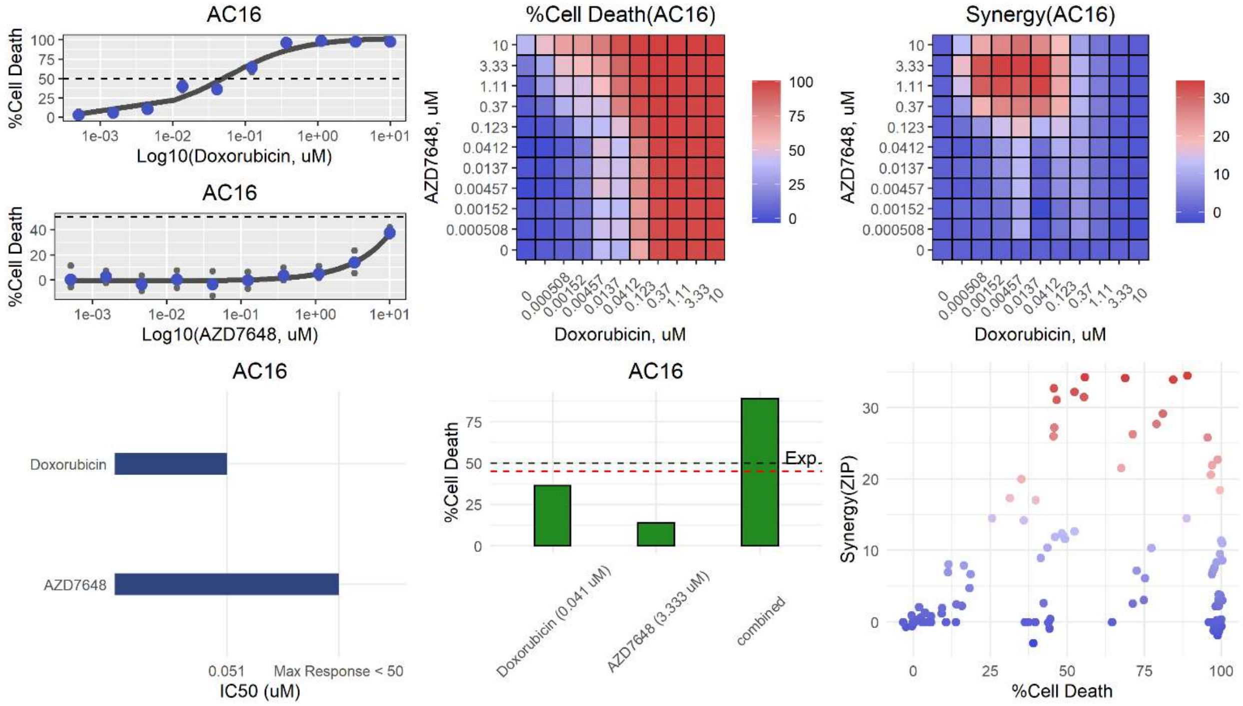
AC16
%Cell Death
Log10(Doxorubicin, uM)
AC16
%Cell Death
Log10(AZD7648, uM)
%Cell Death(AC16)
AZD7648, uM
Doxorubicin, uM
Synergy(AC16)
AZD7648, uM
Doxorubicin, uM
AC16
Doxorubicin
AZD7648
0.051
Max Response < 50
IC50 (uM)
AC16
%Cell Death
Exp.
Doxorubicin (0.041 uM)
AZD7648 (3.333 uM)
combined
Synergy(ZIP)
%Cell Death

**t**

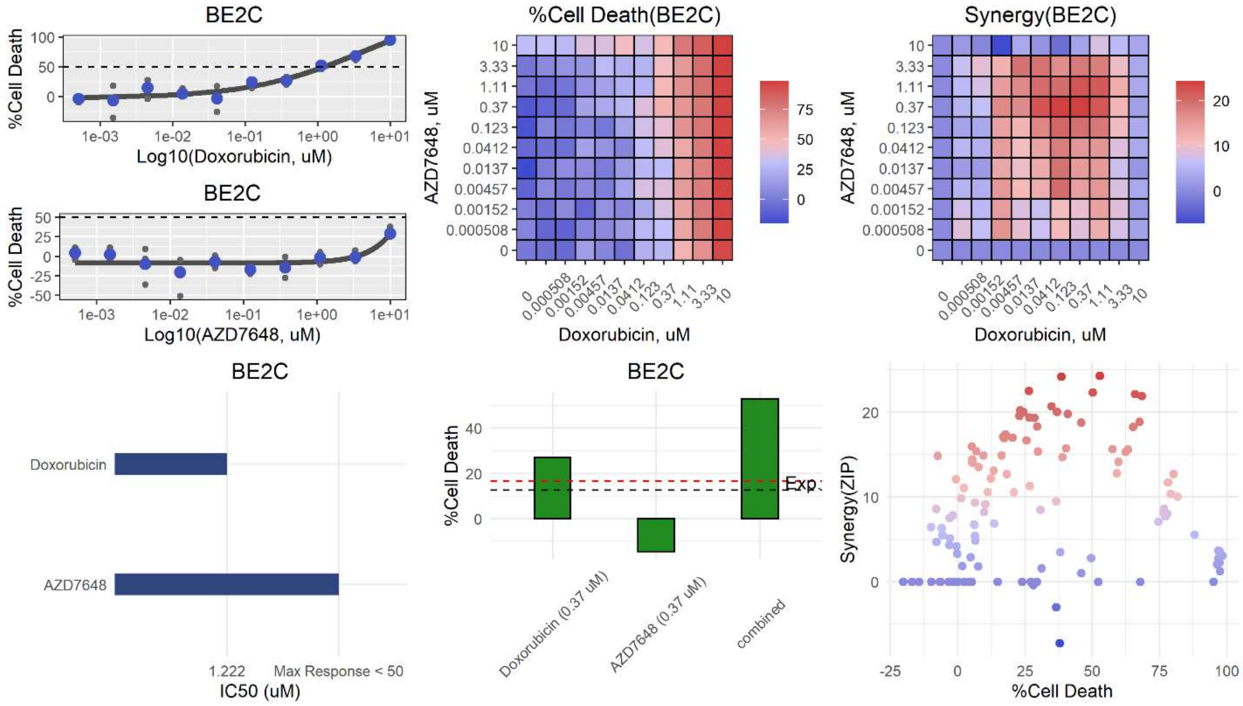
BE2C
%Cell Death
Log10(Doxorubicin, uM)
BE2C
%Cell Death
Log10(AZD7648, uM)
%Cell Death(BE2C)
AZD7648, uM
Doxorubicin, uM
Synergy(BE2C)
AZD7648, uM
Doxorubicin, uM
BE2C
Doxorubicin
AZD7648
1.222
Max Response < 50
IC50 (uM)
BE2C
%Cell Death
Exp.
Doxorubicin (0.37 uM)
AZD7648 (0.37 uM)
combined
Synergy(ZIP)
%Cell Death

**u**

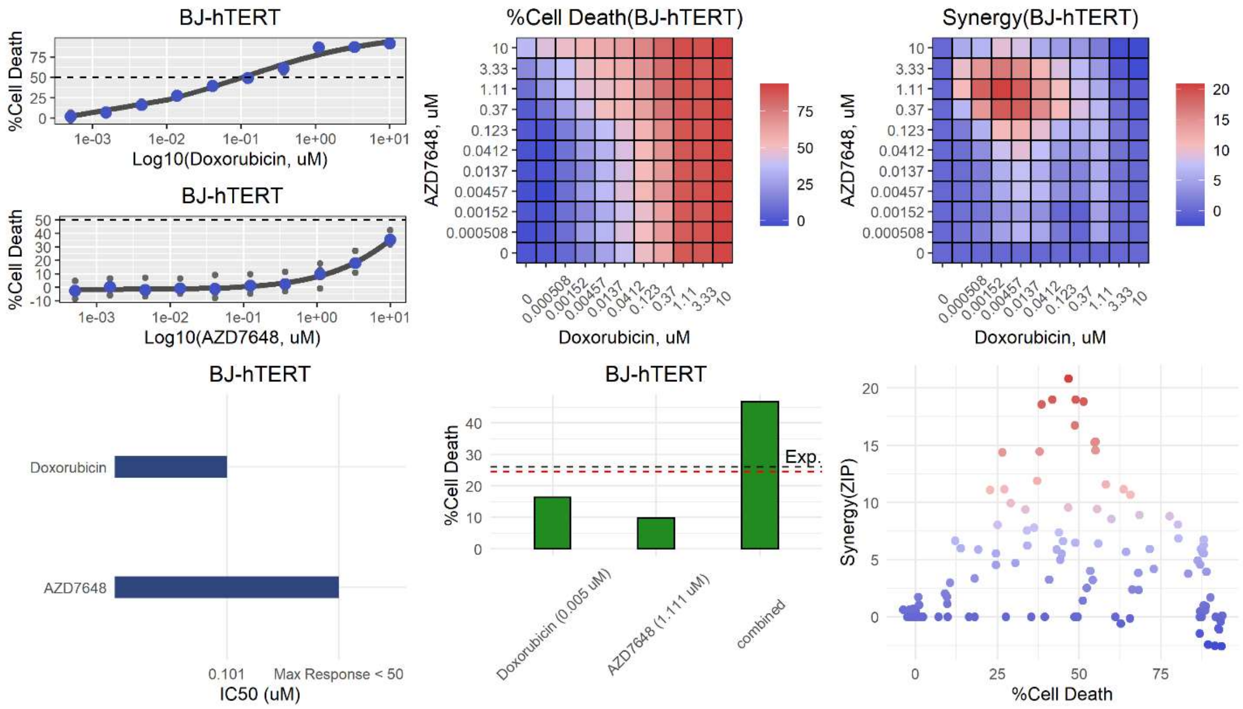
BJ-hTERT
%Cell Death
Log10(Doxorubicin, uM)
BJ-hTERT
%Cell Death
Log10(AZD7648, uM)
%Cell Death(BJ-hTERT)
AZD7648, uM
Doxorubicin, uM
Synergy(BJ-hTERT)
AZD7648, uM
Doxorubicin, uM
BJ-hTERT
Doxorubicin
AZD7648
0.101
Max Response < 50
IC50 (uM)
BJ-hTERT
%Cell Death
Exp.
Doxorubicin (0.005 uM)
AZD7648 (1.111 uM)
combined
Synergy(ZIP)
%Cell Death

**v**

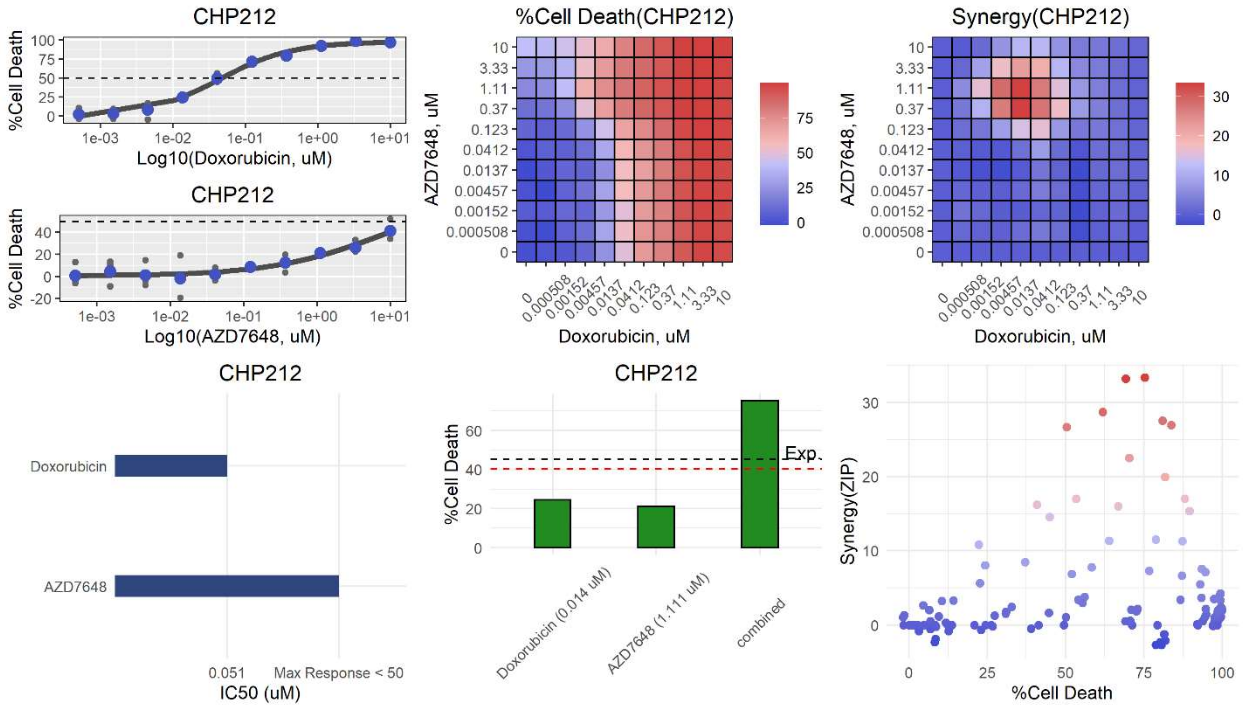
CHP212
%Cell Death
Log10(Doxorubicin, uM)
CHP212
%Cell Death
Log10(AZD7648, uM)
%Cell Death(CHP212)
AZD7648, uM
Doxorubicin, uM
Synergy(CHP212)
AZD7648, uM
Doxorubicin, uM
CHP212
Doxorubicin
AZD7648
0.051
Max Response < 50
IC50 (uM)
CHP212
%Cell Death
Exp.
Doxorubicin (0.014 uM)
AZD7648 (1.111 uM)
combined
Synergy(ZIP)
%Cell Death

**w**

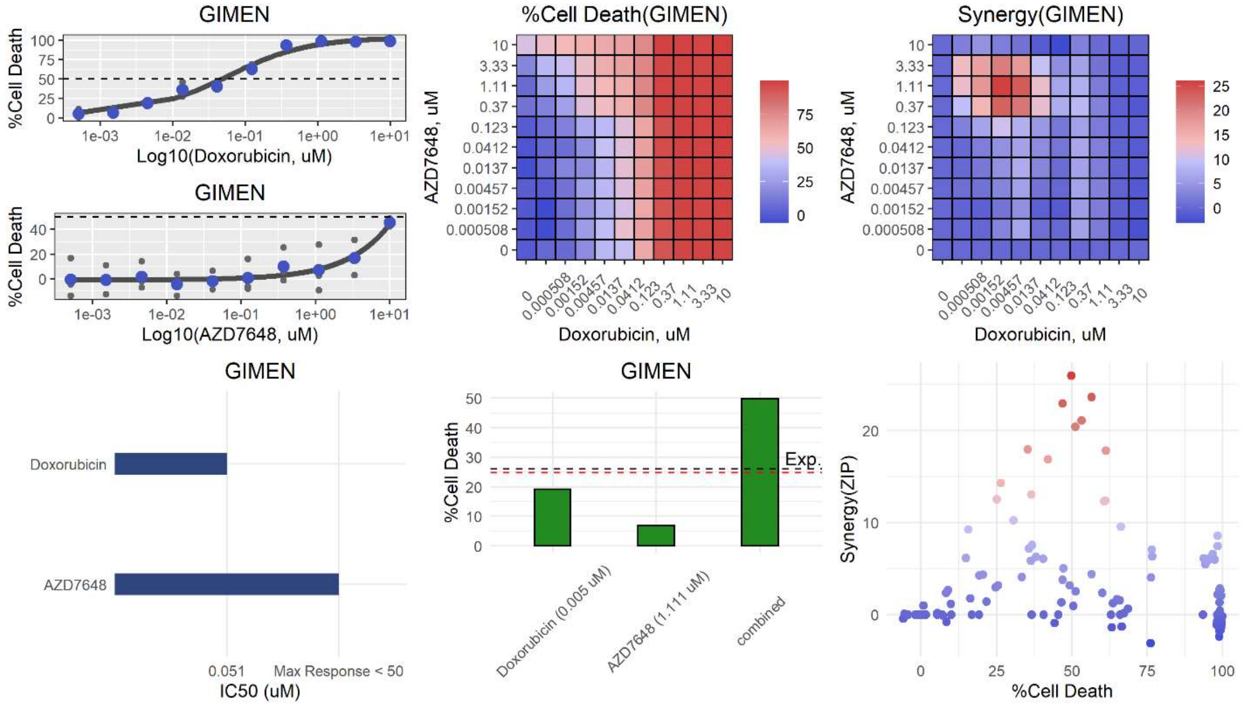
GIMEN
%Cell Death
Log10(Doxorubicin, uM)
GIMEN
Log10(AZD7648, uM)
%Cell Death(GIMEN)
AZD7648, uM
Doxorubicin, uM
Synergy(GIMEN)
GIMEN
Doxorubicin
AZD7648
0.051
Max Response < 50
IC50 (uM)
GIMEN
Exp.
Doxorubicin (0.005 uM)
AZD7648 (1.111 uM)
combined
Synergy(ZIP)
%Cell Death

**x**

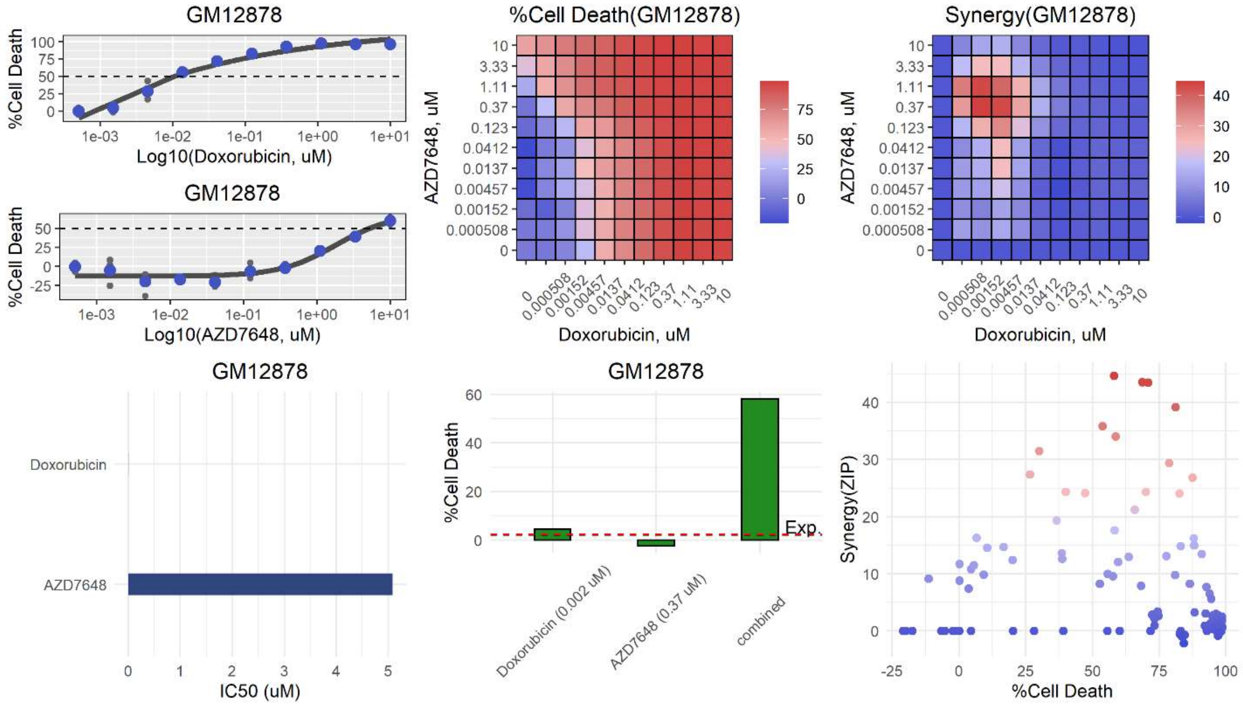
GM12878
%Cell Death
Log10(Doxorubicin, uM)
GM12878
Log10(AZD7648, uM)
%Cell Death(GM12878)
AZD7648, uM
Doxorubicin, uM
Synergy(GM12878)
GM12878
Doxorubicin
AZD7648
IC50 (uM)
GM12878
Exp.
Doxorubicin (0.002 uM)
AZD7648 (0.37 uM)
combined
Synergy(ZIP)
%Cell Death

**y**

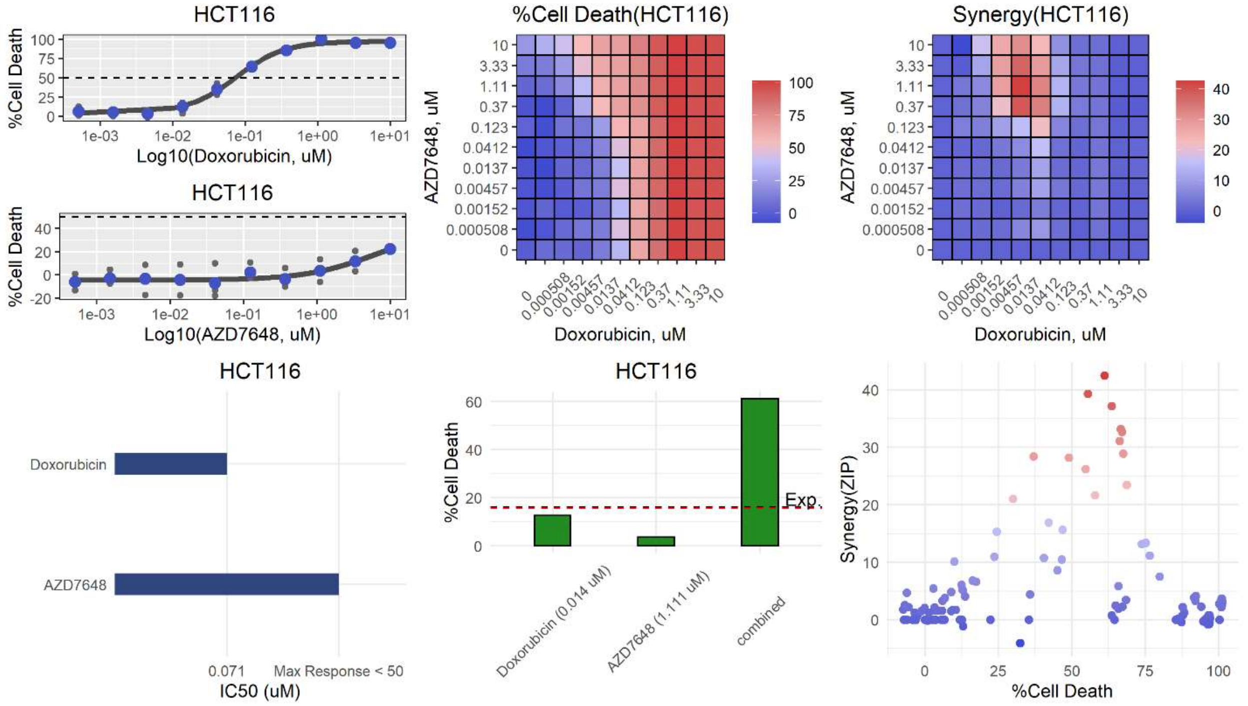

HCT116
%Cell Death
Log10(Doxorubicin, uM)
HCT116
%Cell Death
Log10(AZD7648, uM)
%Cell Death(HCT116)
AZD7648, uM
Doxorubicin, uM
Synergy(HCT116)
AZD7648, uM
Doxorubicin, uM
HCT116
Doxorubicin
AZD7648
0.071
Max Response < 50
IC50 (uM)
HCT116
%Cell Death
Exp.
Doxorubicin (0.014 uM)
AZD7648 (1.111 uM)
combined
Synergy(ZIP)
%Cell Death


**z**

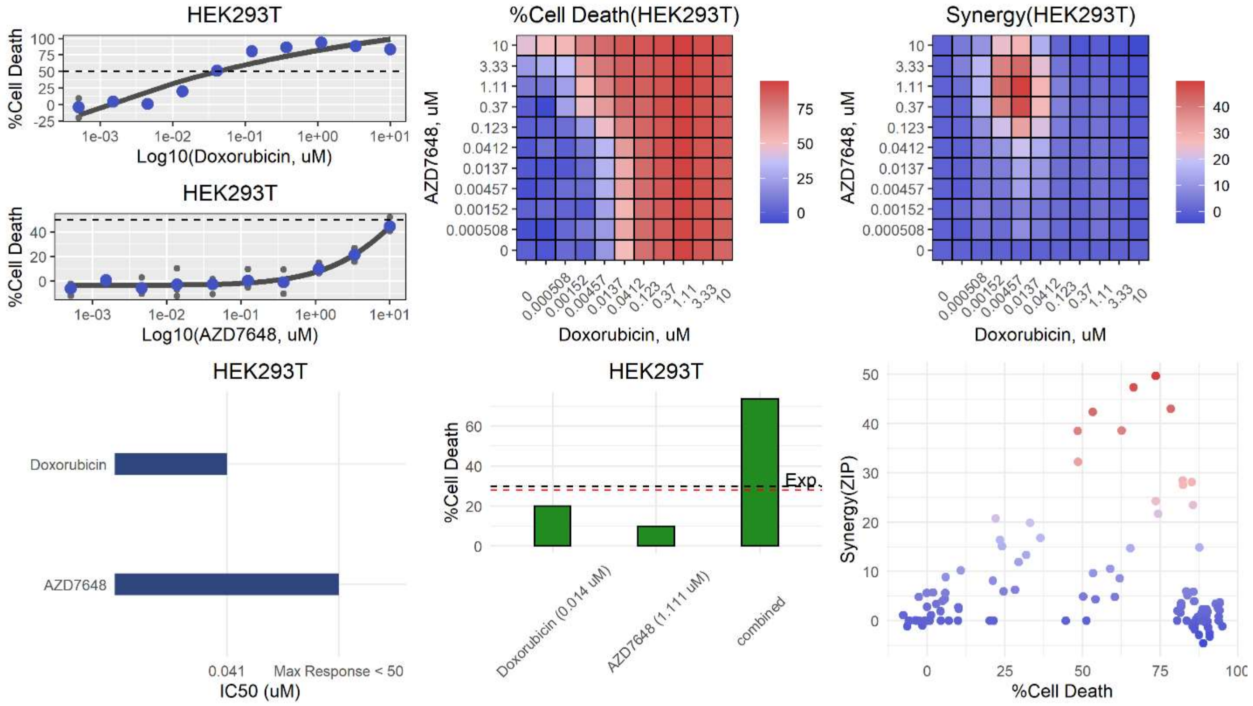

HEK293T
%Cell Death
Log10(Doxorubicin, uM)
HEK293T
%Cell Death
Log10(AZD7648, uM)
%Cell Death(HEK293T)
AZD7648, uM
Doxorubicin, uM
Synergy(HEK293T)
AZD7648, uM
Doxorubicin, uM
HEK293T
Doxorubicin
AZD7648
0.041
Max Response < 50
IC50 (uM)
HEK293T
%Cell Death
Exp.
Doxorubicin (0.014 uM)
AZD7648 (1.111 uM)
combined
Synergy(ZIP)
%Cell Death

**aa**

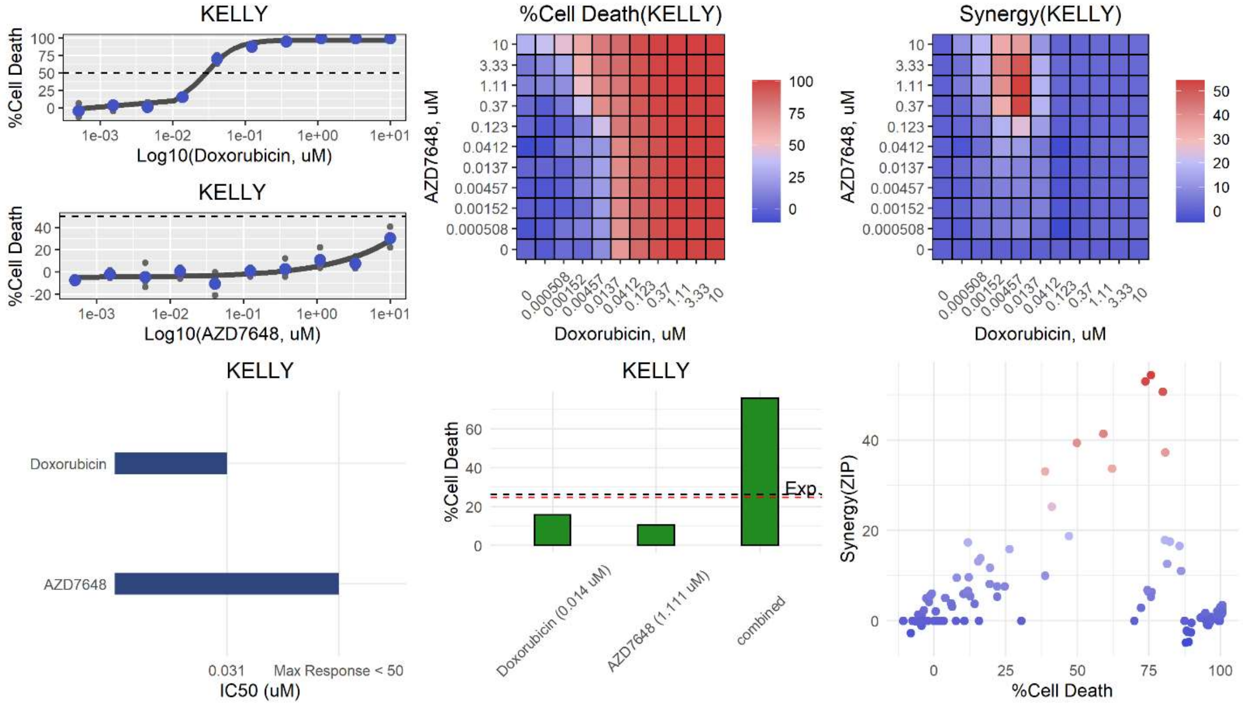
KELLY
%Cell Death
Log10(Doxorubicin, uM)
Log10(AZD7648, uM)
%Cell Death(KELLY)
AZD7648, uM
Doxorubicin, uM
Synergy(KELLY)
Doxorubicin
AZD7648
0.031
Max Response < 50
IC50 (uM)
Exp.
Doxorubicin (0.014 uM)
AZD7648 (1.111 uM)
combined
Synergy(ZIP)

**ab**

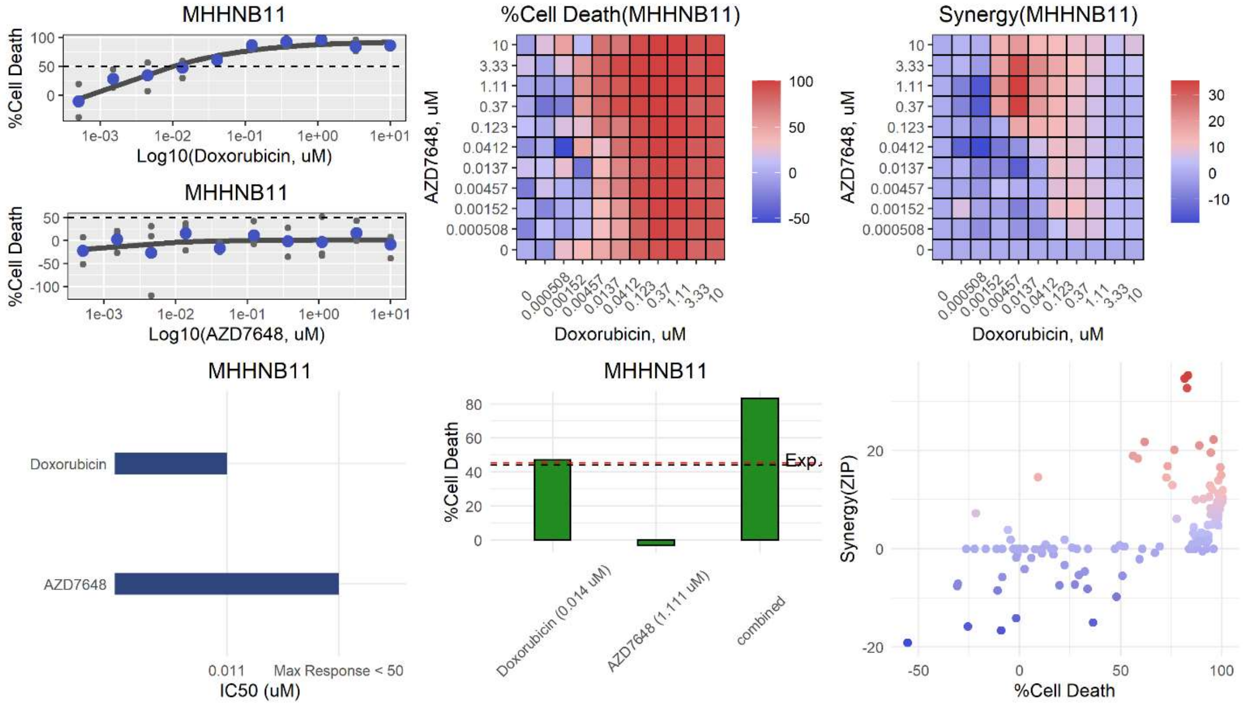
MHHNB11
%Cell Death
Log10(Doxorubicin, uM)
Log10(AZD7648, uM)
%Cell Death(MHHNB11)
AZD7648, uM
Doxorubicin, uM
Synergy(MHHNB11)
Doxorubicin
AZD7648
0.011
Max Response < 50
IC50 (uM)
Exp.
Doxorubicin (0.014 uM)
AZD7648 (1.111 uM)
combined
Synergy(ZIP)

**ac**

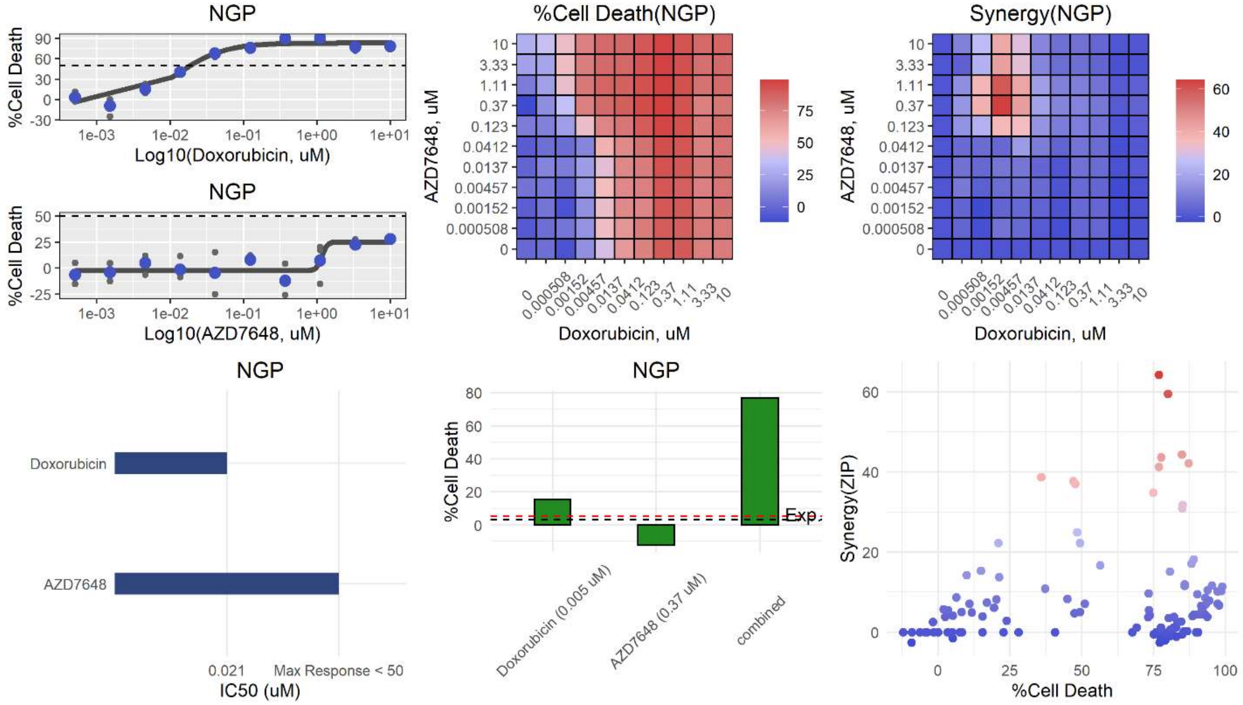
NGP
%Cell Death
Log10(Doxorubicin, uM)
NGP
%Cell Death
Log10(AZD7648, uM)
%Cell Death(NGP)
AZD7648, uM
Doxorubicin, uM
Synergy(NGP)
AZD7648, uM
Doxorubicin, uM
NGP
Doxorubicin
AZD7648
0.021
Max Response < 50
IC50 (uM)
NGP
%Cell Death
Exp.
Doxorubicin (0.005 uM)
AZD7648 (0.37 uM)
combined
Synergy(ZIP)
%Cell Death

**ad**

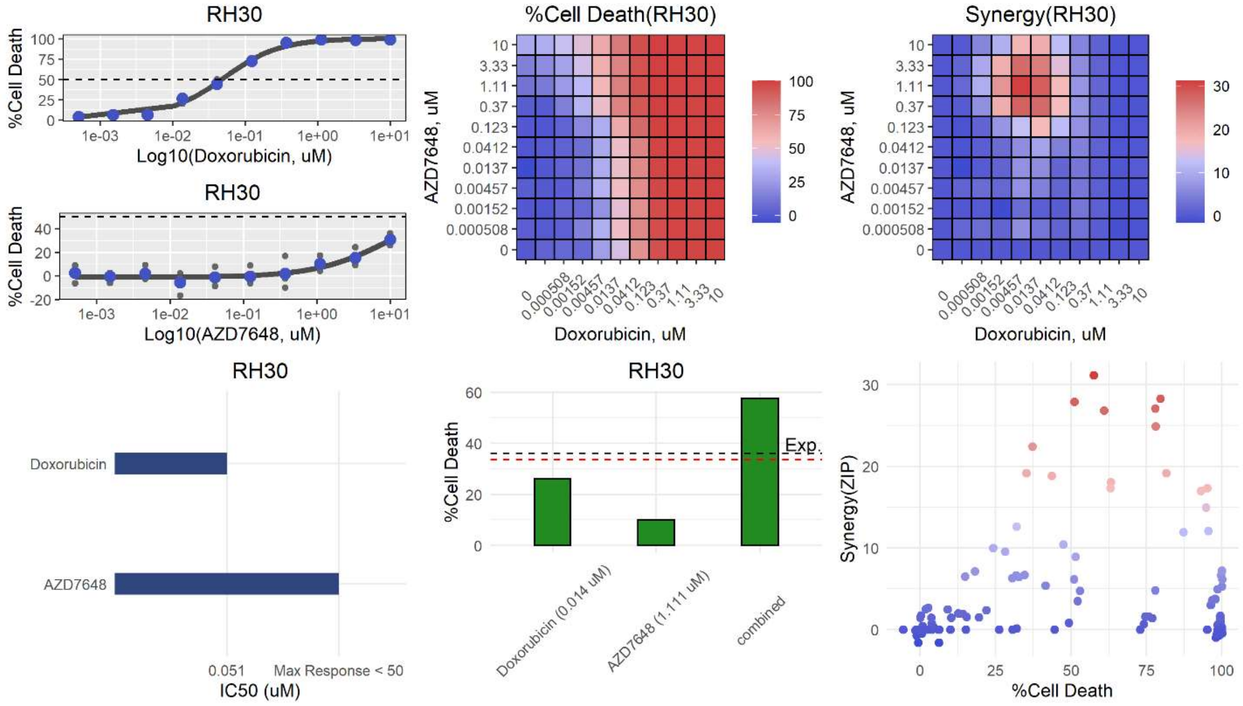
RH30
%Cell Death
Log10(Doxorubicin, uM)
RH30
%Cell Death
Log10(AZD7648, uM)
%Cell Death(RH30)
AZD7648, uM
Doxorubicin, uM
Synergy(RH30)
AZD7648, uM
Doxorubicin, uM
RH30
Doxorubicin
AZD7648
0.051
Max Response < 50
IC50 (uM)
RH30
%Cell Death
Exp.
Doxorubicin (0.014 uM)
AZD7648 (1.111 uM)
combined
Synergy(ZIP)
%Cell Death

**ae**

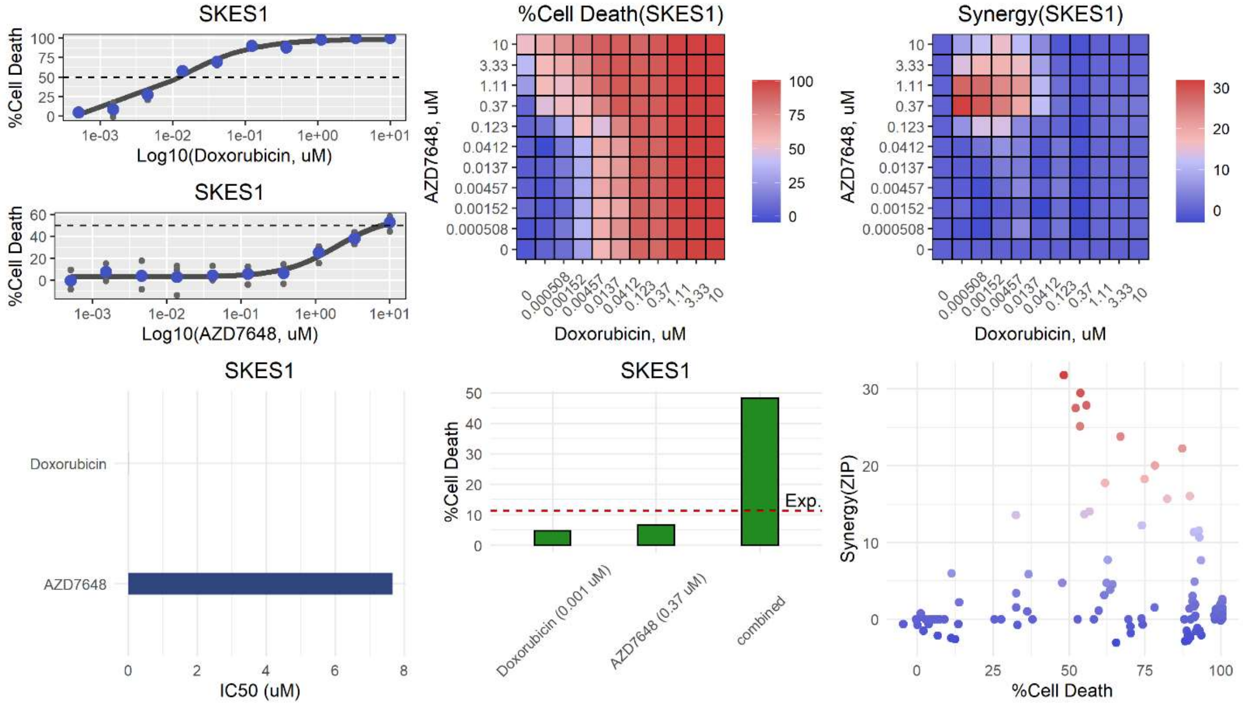
SKES1
%Cell Death
Log10(Doxorubicin, uM)
SKES1
Log10(AZD7648, uM)
%Cell Death(SKES1)
AZD7648, uM
Doxorubicin, uM
Synergy(SKES1)
SKES1
Doxorubicin
AZD7648
IC50 (uM)
SKES1
Exp.
Doxorubicin (0.001 uM)
AZD7648 (0.37 uM)
combined
Synergy(ZIP)
%Cell Death

**af**

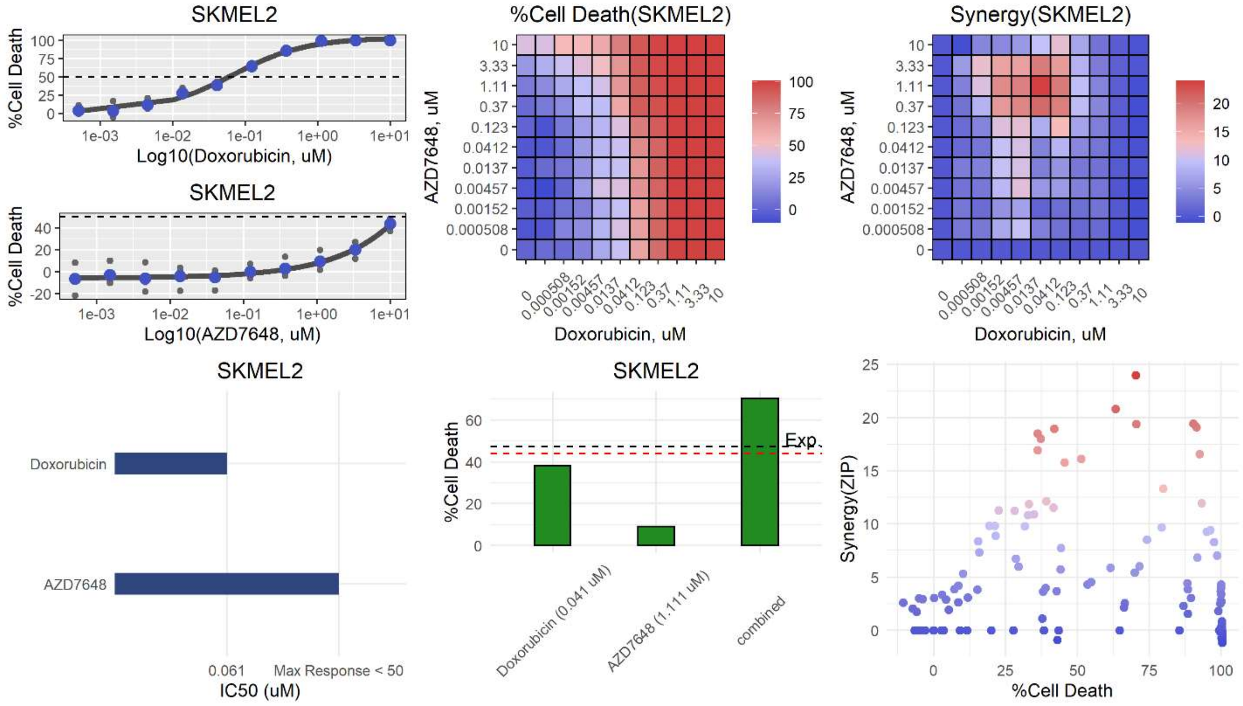
SKMEL2
%Cell Death
Log10(Doxorubicin, uM)
SKMEL2
Log10(AZD7648, uM)
%Cell Death(SKMEL2)
AZD7648, uM
Doxorubicin, uM
Synergy(SKMEL2)
SKMEL2
Doxorubicin
AZD7648
0.061
Max Response < 50
IC50 (uM)
SKMEL2
Exp.
Doxorubicin (0.041 uM)
AZD7648 (1.111 uM)
combined
Synergy(ZIP)
%Cell Death

**ag**

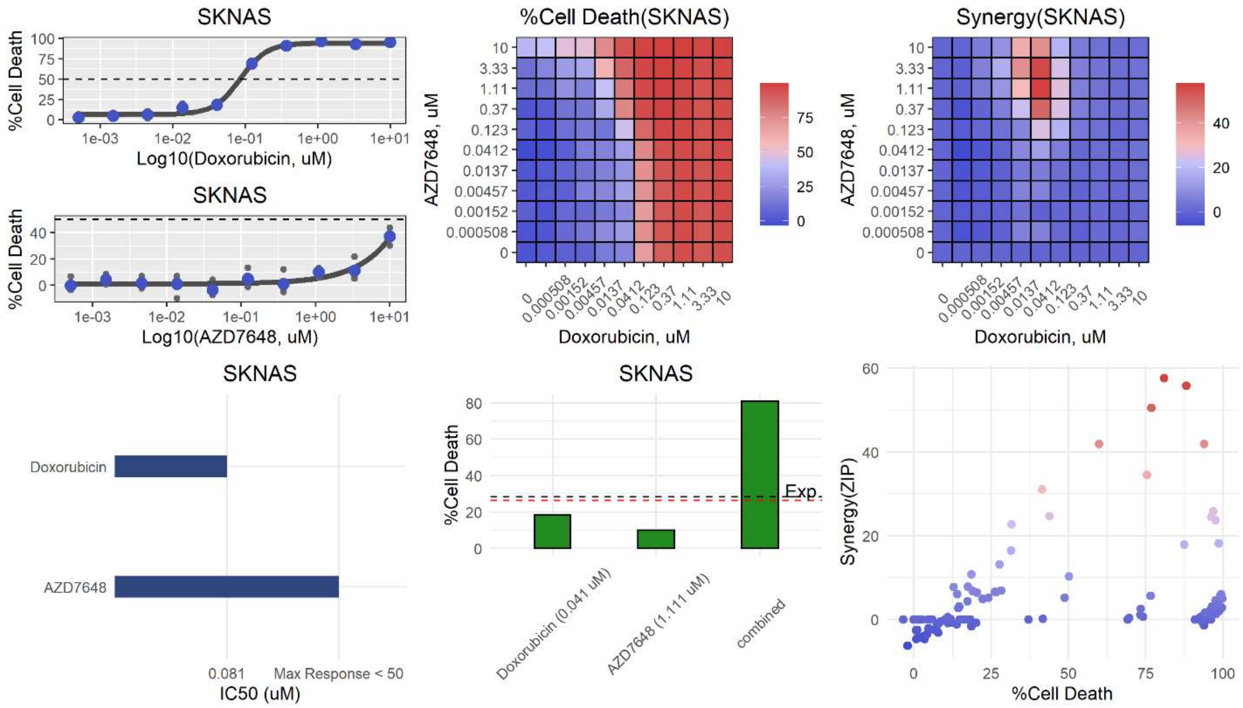
SKNAS
%Cell Death
Log10(Doxorubicin, uM)
Log10(AZD7648, uM)
%Cell Death(SKNAS)
Synergy(SKNAS)
AZD7648, uM
Doxorubicin, uM
Doxorubicin
AZD7648
0.081
Max Response < 50
IC50 (uM)
Exp.
Doxorubicin (0.041 uM)
AZD7648 (1.111 uM)
combined
Synergy(ZIP)

**ah**

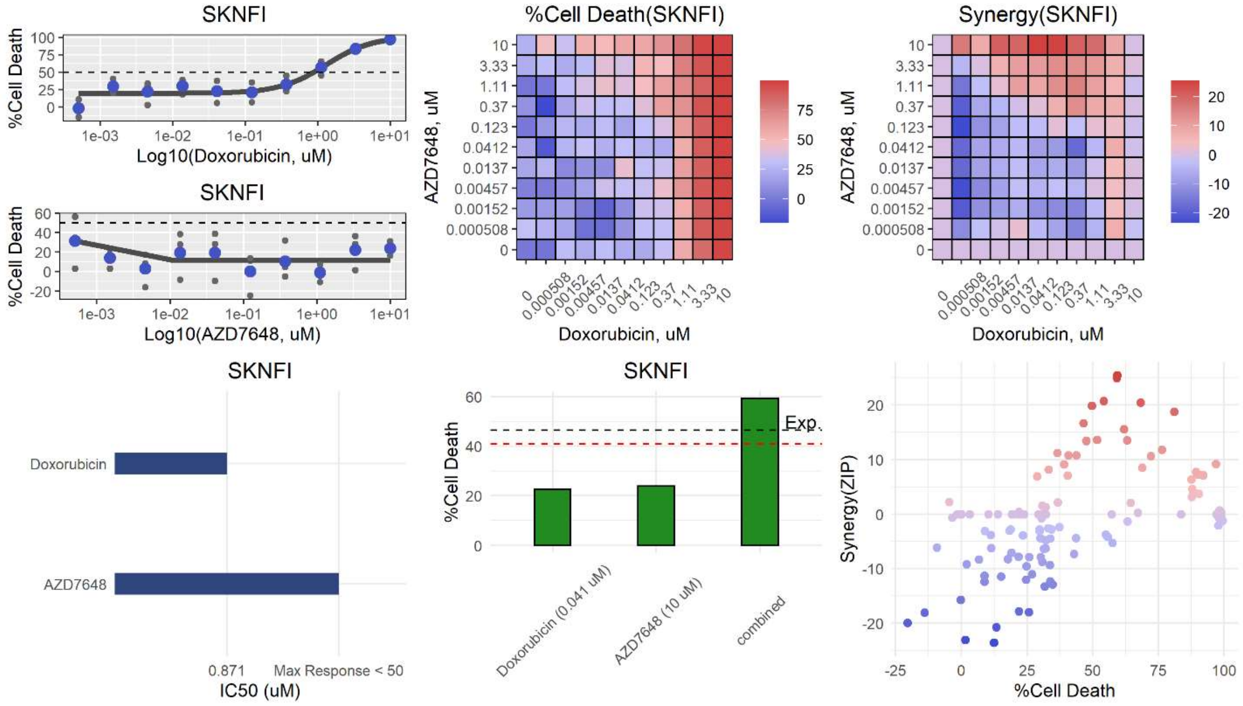
SKNFI
%Cell Death
Log10(Doxorubicin, uM)
Log10(AZD7648, uM)
%Cell Death(SKNFI)
Synergy(SKNFI)
AZD7648, uM
Doxorubicin, uM
Doxorubicin
AZD7648
0.871
Max Response < 50
IC50 (uM)
Exp.
Doxorubicin (0.041 uM)
AZD7648 (10 uM)
combined
Synergy(ZIP)

**ai**

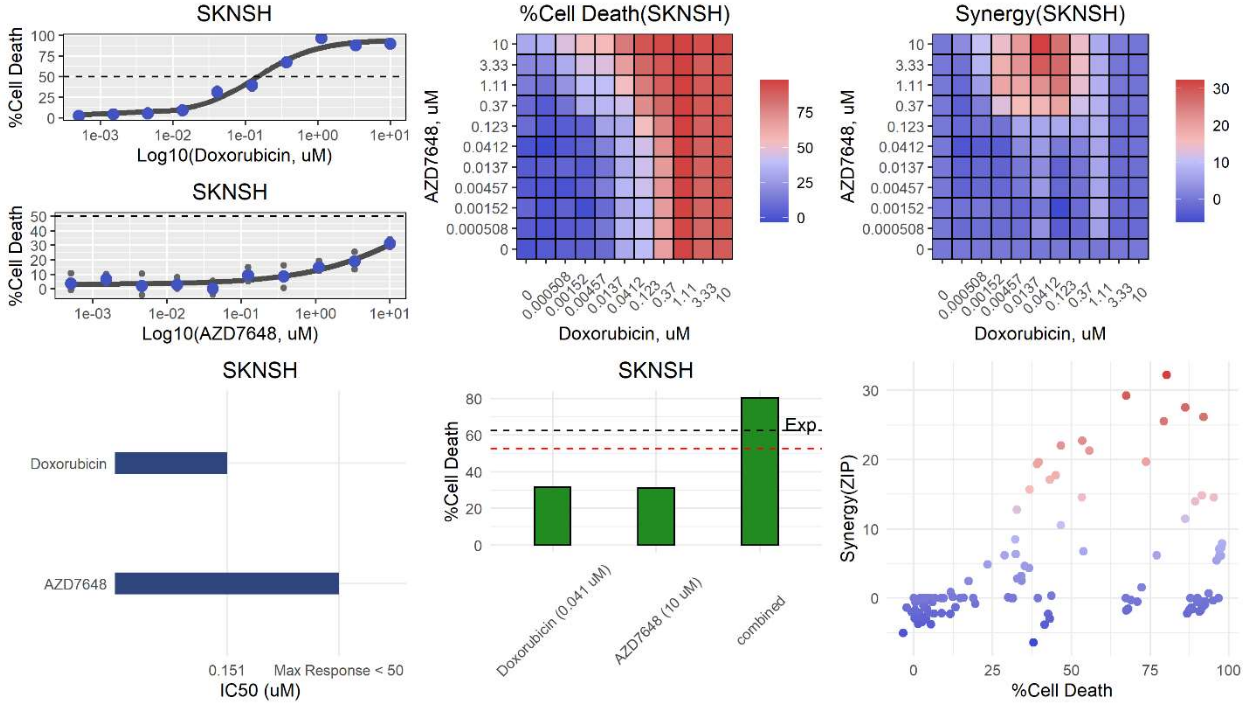
SKNSH
%Cell Death
Log10(Doxorubicin, uM)
SKNSH
%Cell Death
Log10(AZD7648, uM)
%Cell Death(SKNSH)
AZD7648, uM
Doxorubicin, uM
Synergy(SKNSH)
AZD7648, uM
Doxorubicin, uM
SKNSH
Doxorubicin
AZD7648
0.151
Max Response < 50
IC50 (uM)
SKNSH
%Cell Death
Exp.
Doxorubicin (0.041 uM)
AZD7648 (10 uM)
combined
Synergy(ZIP)
%Cell Death

**aj**

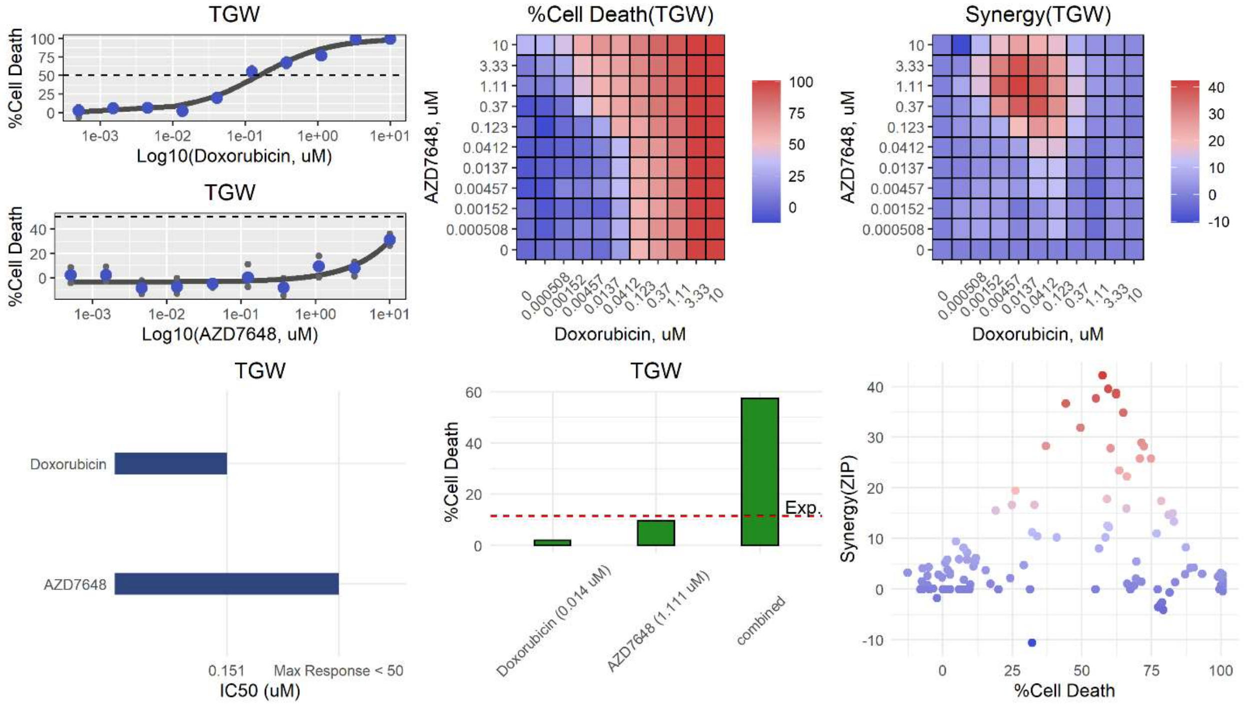
TGW
%Cell Death
Log10(Doxorubicin, uM)
TGW
%Cell Death
Log10(AZD7648, uM)
%Cell Death(TGW)
AZD7648, uM
Doxorubicin, uM
Synergy(TGW)
AZD7648, uM
Doxorubicin, uM
TGW
Doxorubicin
AZD7648
0.151
Max Response < 50
IC50 (uM)
TGW
%Cell Death
Exp.
Doxorubicin (0.014 uM)
AZD7648 (1.111 uM)
combined
Synergy(ZIP)
%Cell Death

**ak**

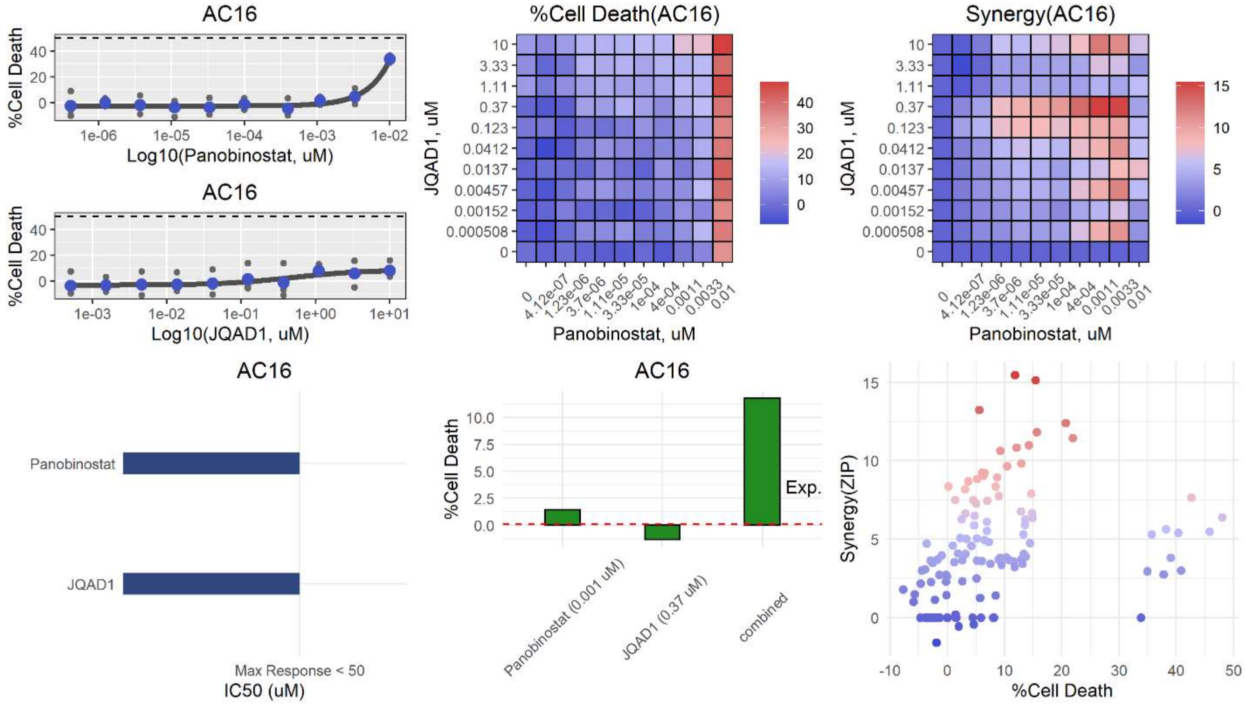

AC16
%Cell Death
Log10(Panobinostat, uM)
AC16
Log10(JQAD1, uM)
%Cell Death(AC16)
JQAD1, uM
Panobinostat, uM
Synergy(AC16)
AC16
Panobinostat
JQAD1
Max Response < 50
IC50 (uM)
AC16
Exp.
Panobinostat (0.001 uM)
JQAD1 (0.37 uM)
combined
Synergy(ZIP)
%Cell Death


**al**

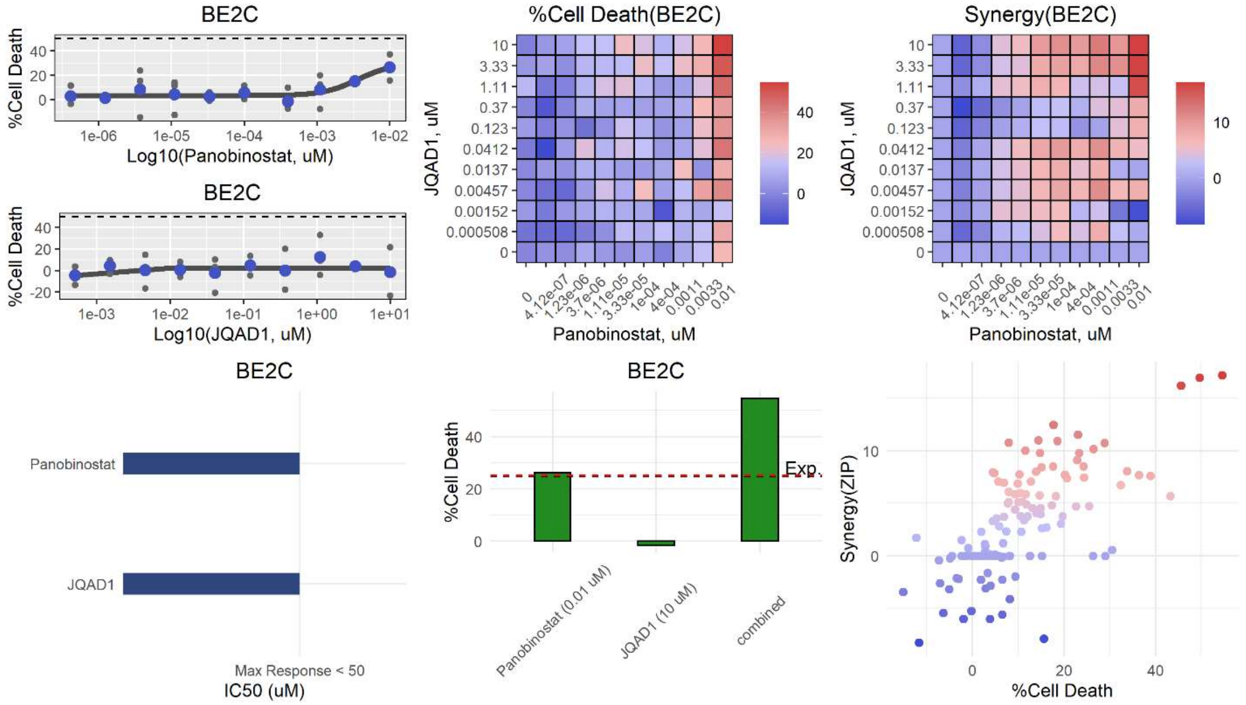

BE2C
%Cell Death
Log10(Panobinostat, uM)
BE2C
Log10(JQAD1, uM)
%Cell Death(BE2C)
JQAD1, uM
Panobinostat, uM
Synergy(BE2C)
BE2C
Panobinostat
JQAD1
Max Response < 50
IC50 (uM)
BE2C
Exp.
Panobinostat (0.01 uM)
JQAD1 (10 uM)
combined
Synergy(ZIP)
%Cell Death

**am**

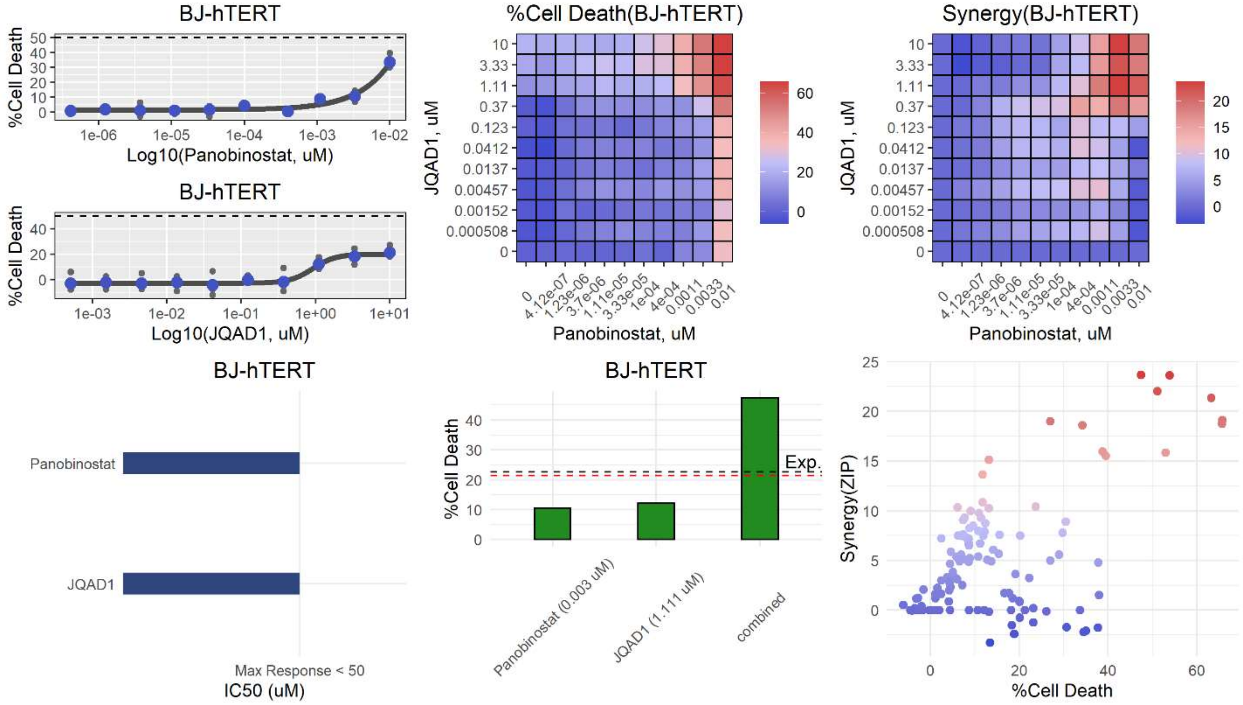

BJ-hTERT
%Cell Death
Log10(Panobinostat, uM)
BJ-hTERT
%Cell Death
Log10(JQAD1, uM)
%Cell Death(BJ-hTERT)
JQAD1, uM
Panobinostat, uM
Synergy(BJ-hTERT)
JQAD1, uM
Panobinostat, uM
BJ-hTERT
Panobinostat
JQAD1
Max Response < 50
IC50 (uM)
BJ-hTERT
%Cell Death
Exp.
Panobinostat (0.003 uM)
JQAD1 (1.111 uM)
combined
Synergy(ZIP)
%Cell Death


**an**

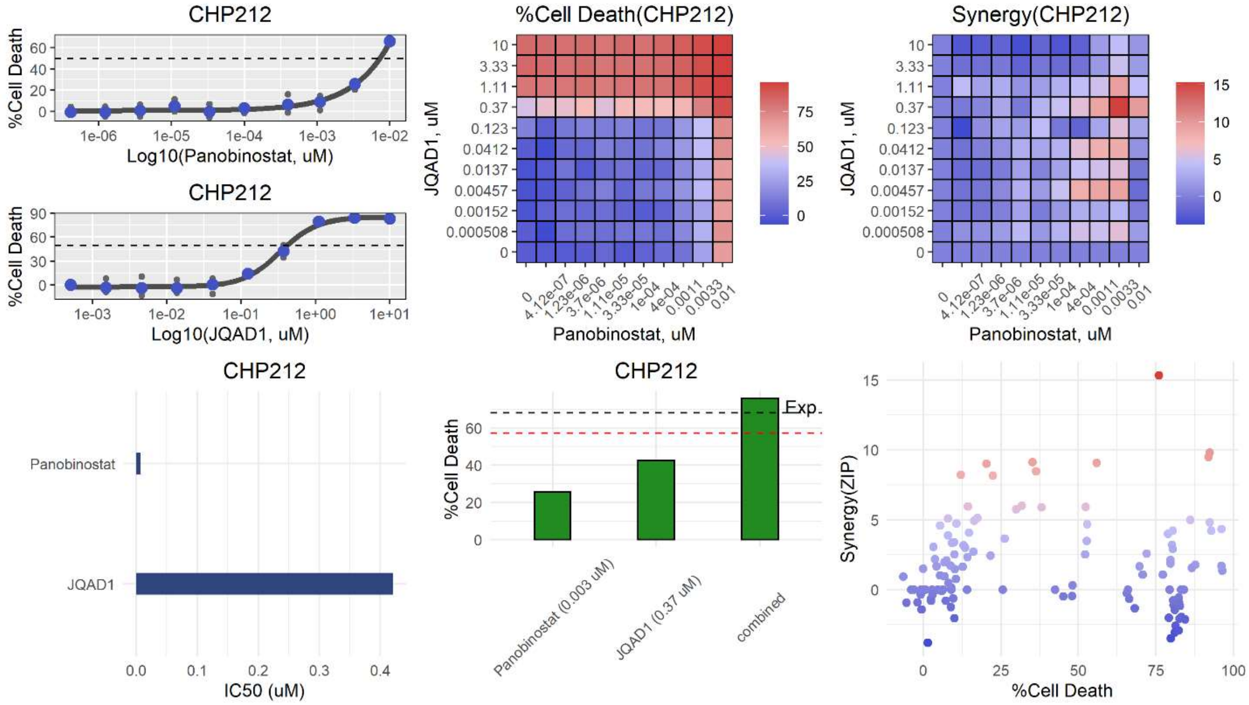

CHP212
%Cell Death
Log10(Panobinostat, uM)
CHP212
%Cell Death
Log10(JQAD1, uM)
%Cell Death(CHP212)
JQAD1, uM
Panobinostat, uM
Synergy(CHP212)
JQAD1, uM
Panobinostat, uM
CHP212
Panobinostat
JQAD1
IC50 (uM)
CHP212
%Cell Death
Exp.
Panobinostat (0.003 uM)
JQAD1 (0.37 uM)
combined
Synergy(ZIP)
%Cell Death

**ao**

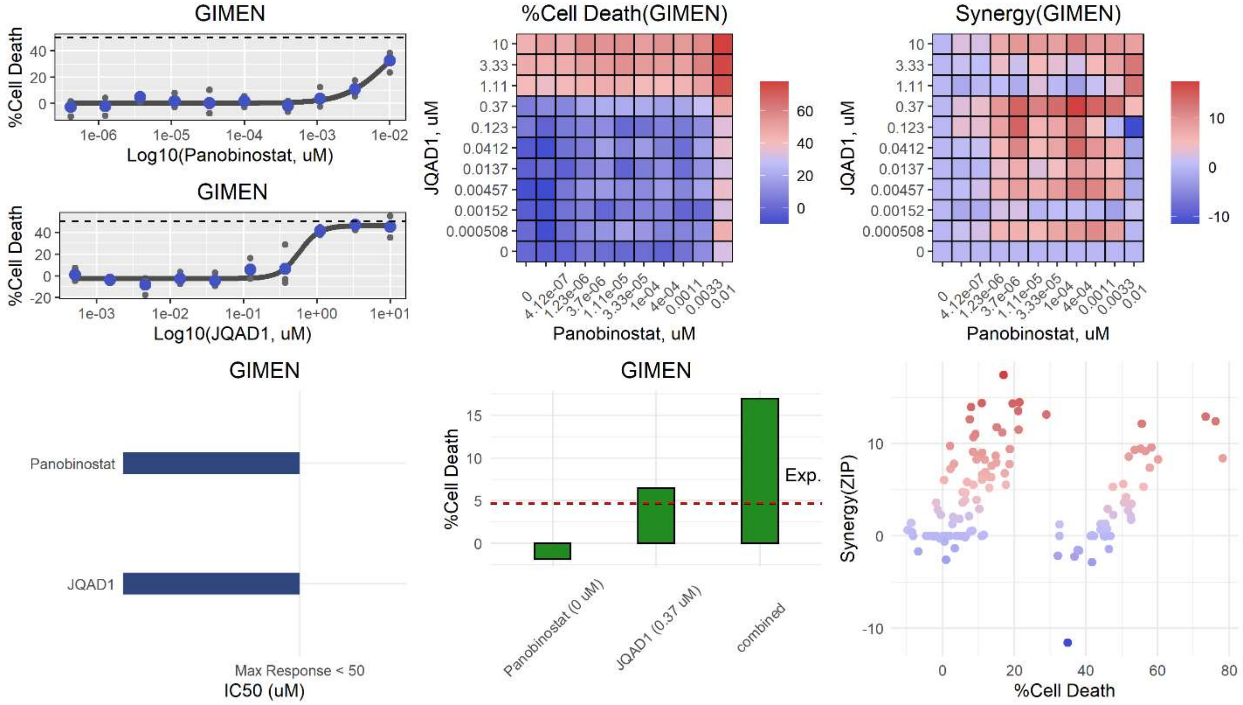
GIMEN
%Cell Death
Log10(Panobinostat, uM)
GIMEN
%Cell Death
Log10(JQAD1, uM)
%Cell Death(GIMEN)
JQAD1, uM
Panobinostat, uM
Synergy(GIMEN)
JQAD1, uM
Panobinostat, uM
GIMEN
Panobinostat
JQAD1
Max Response < 50
IC50 (uM)
GIMEN
%Cell Death
Exp.
Panobinostat (0 uM)
JQAD1 (0.37 uM)
combined
Synergy(ZIP)
%Cell Death

**ap**

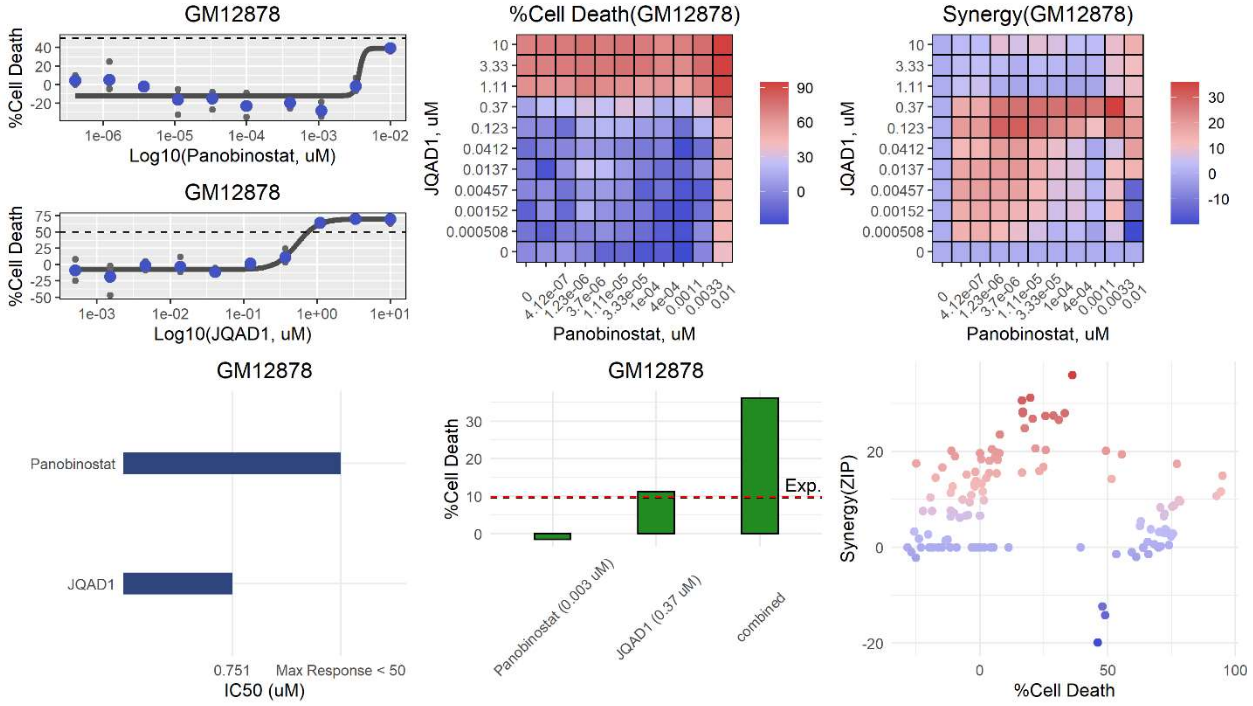
GM12878
%Cell Death
Log10(Panobinostat, uM)
GM12878
%Cell Death
Log10(JQAD1, uM)
%Cell Death(GM12878)
JQAD1, uM
Panobinostat, uM
Synergy(GM12878)
JQAD1, uM
Panobinostat, uM
GM12878
Panobinostat
JQAD1
0.751
Max Response < 50
IC50 (uM)
GM12878
%Cell Death
Exp.
Panobinostat (0.003 uM)
JQAD1 (0.37 uM)
combined
Synergy(ZIP)
%Cell Death

**aq**

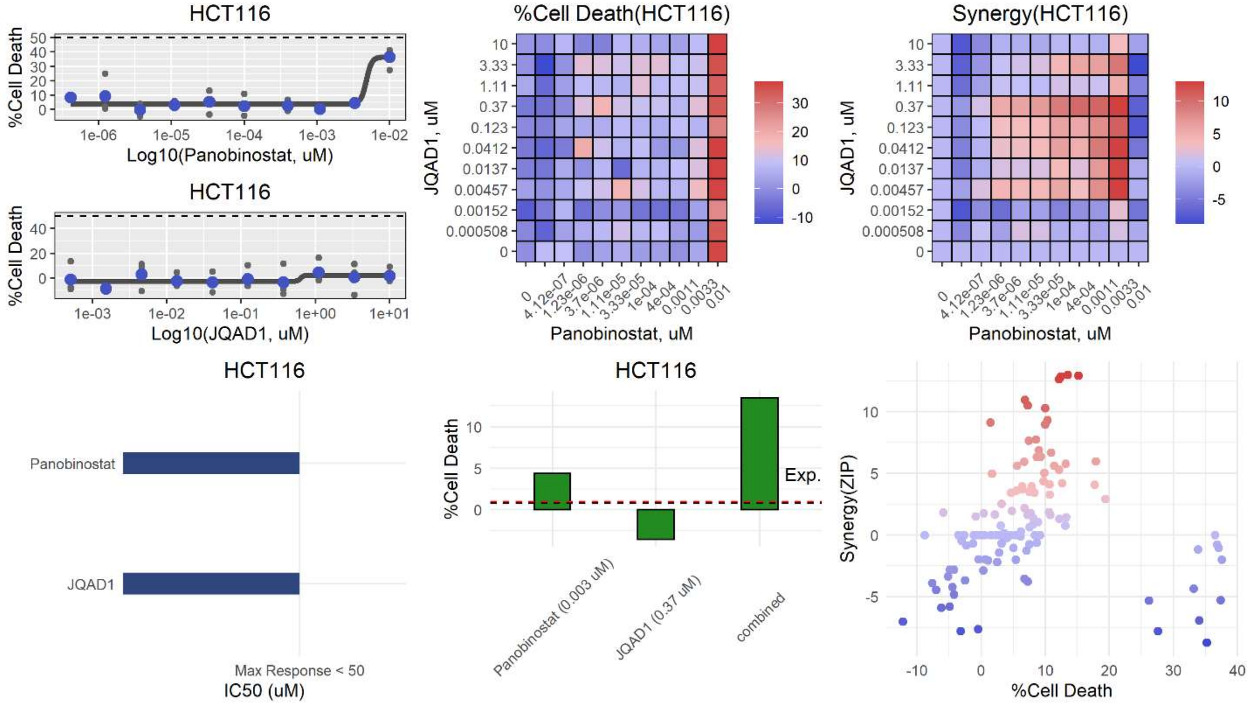
HCT116
%Cell Death
Log10(Panobinostat, uM)
HCT116
Log10(JQAD1, uM)
%Cell Death(HCT116)
JQAD1, uM
Panobinostat, uM
Synergy(HCT116)
HCT116
Panobinostat
JQAD1
Max Response < 50
IC50 (uM)
HCT116
Exp.
Panobinostat (0.003 uM)
JQAD1 (0.37 uM)
combined
Synergy(ZIP)
%Cell Death

**ar**

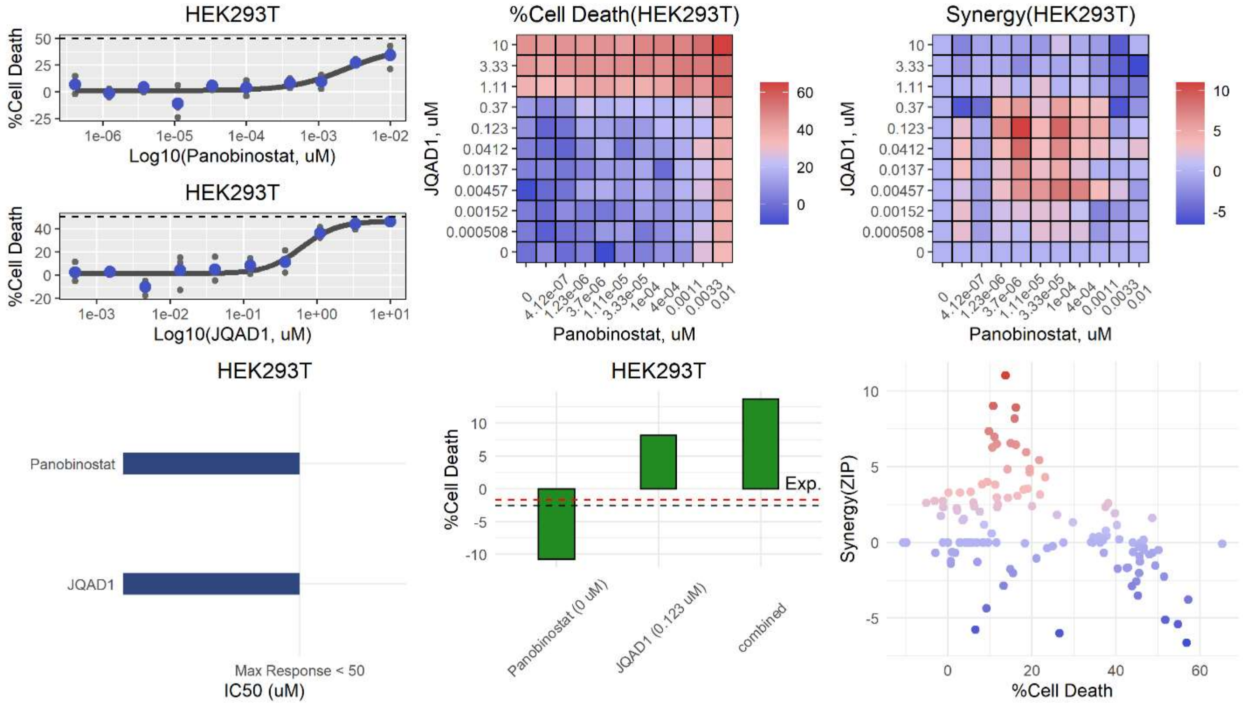
HEK293T
%Cell Death
Log10(Panobinostat, uM)
HEK293T
Log10(JQAD1, uM)
%Cell Death(HEK293T)
JQAD1, uM
Panobinostat, uM
Synergy(HEK293T)
HEK293T
Panobinostat
JQAD1
Max Response < 50
IC50 (uM)
HEK293T
Exp.
Panobinostat (0 uM)
JQAD1 (0.123 uM)
combined
Synergy(ZIP)
%Cell Death

**as**

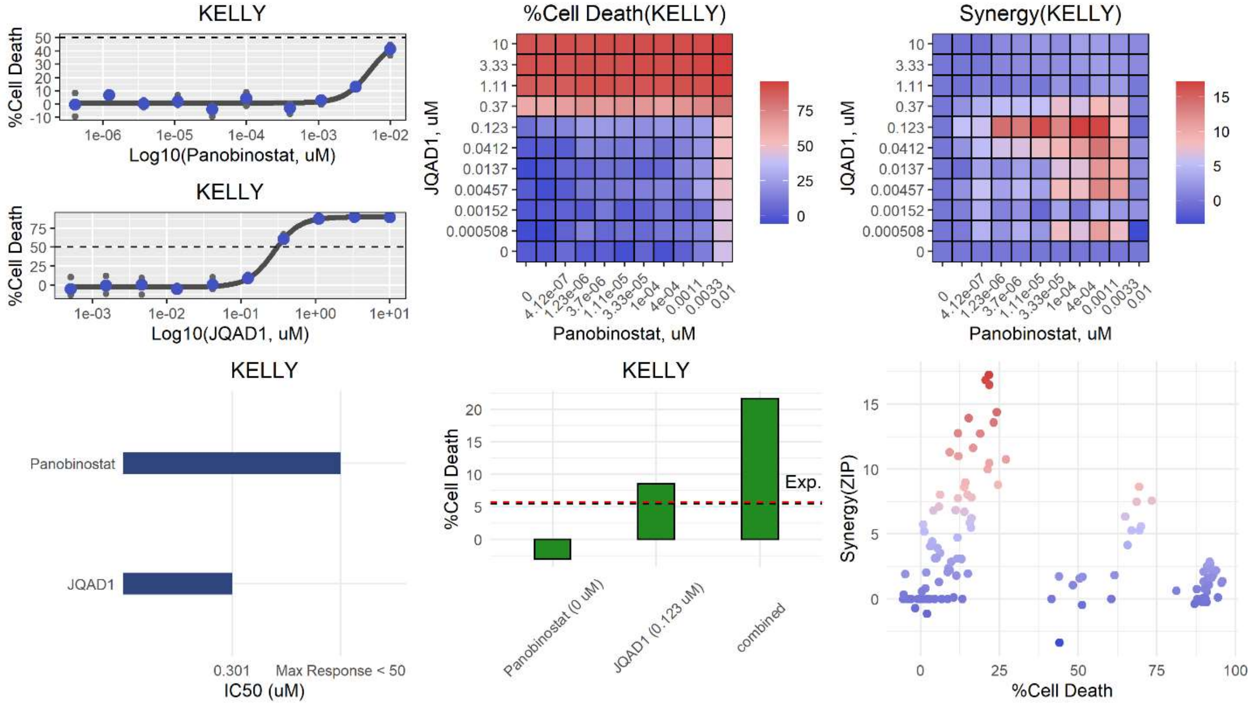

KELLY
%Cell Death
Log10(Panobinostat, uM)
KELLY
%Cell Death
Log10(JQAD1, uM)
%Cell Death(KELLY)
JQAD1, uM
Panobinostat, uM
Synergy(KELLY)
JQAD1, uM
Panobinostat, uM
KELLY
Panobinostat
JQAD1
0.301
Max Response < 50
IC50 (uM)
KELLY
%Cell Death
Exp.
Panobinostat (0 uM)
JQAD1 (0.123 uM)
combined
Synergy(ZIP)
%Cell Death


**at**

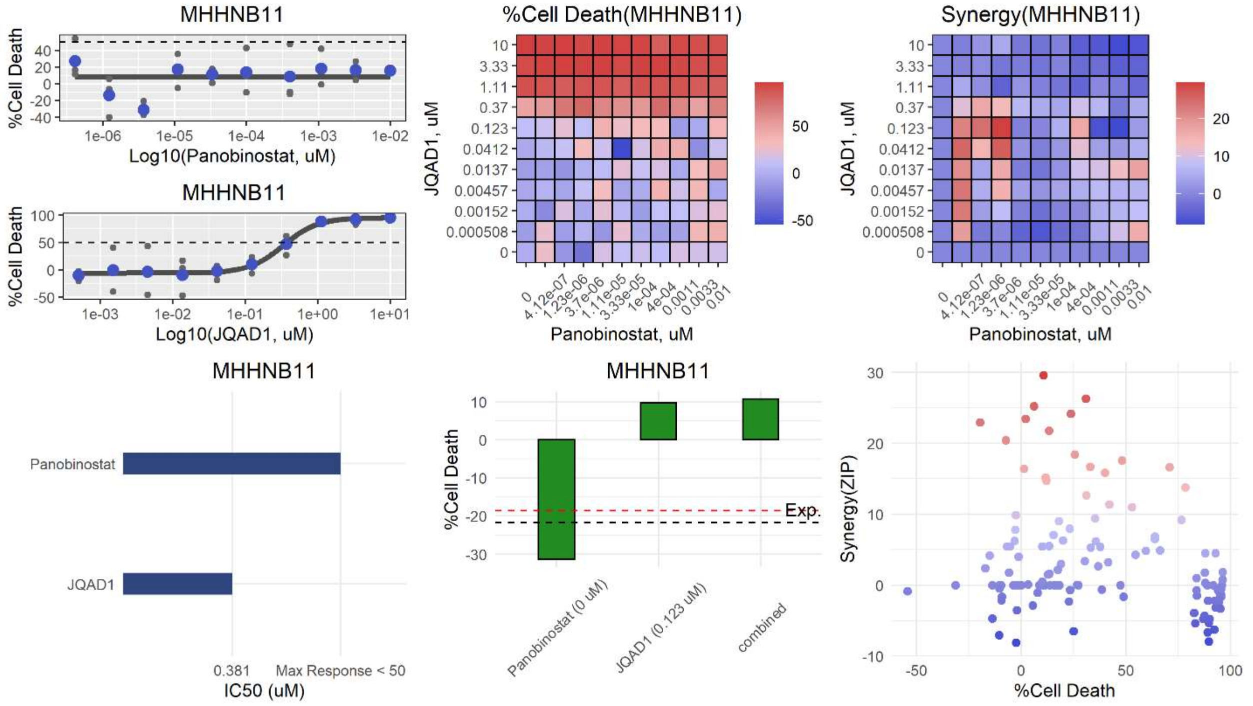

MHHNB11
%Cell Death
Log10(Panobinostat, uM)
MHHNB11
%Cell Death
Log10(JQAD1, uM)
%Cell Death(MHHNB11)
JQAD1, uM
Panobinostat, uM
Synergy(MHHNB11)
JQAD1, uM
Panobinostat, uM
MHHNB11
Panobinostat
JQAD1
0.381
Max Response < 50
IC50 (uM)
MHHNB11
%Cell Death
Exp.
Panobinostat (0 uM)
JQAD1 (0.123 uM)
combined
Synergy(ZIP)
%Cell Death

**au**

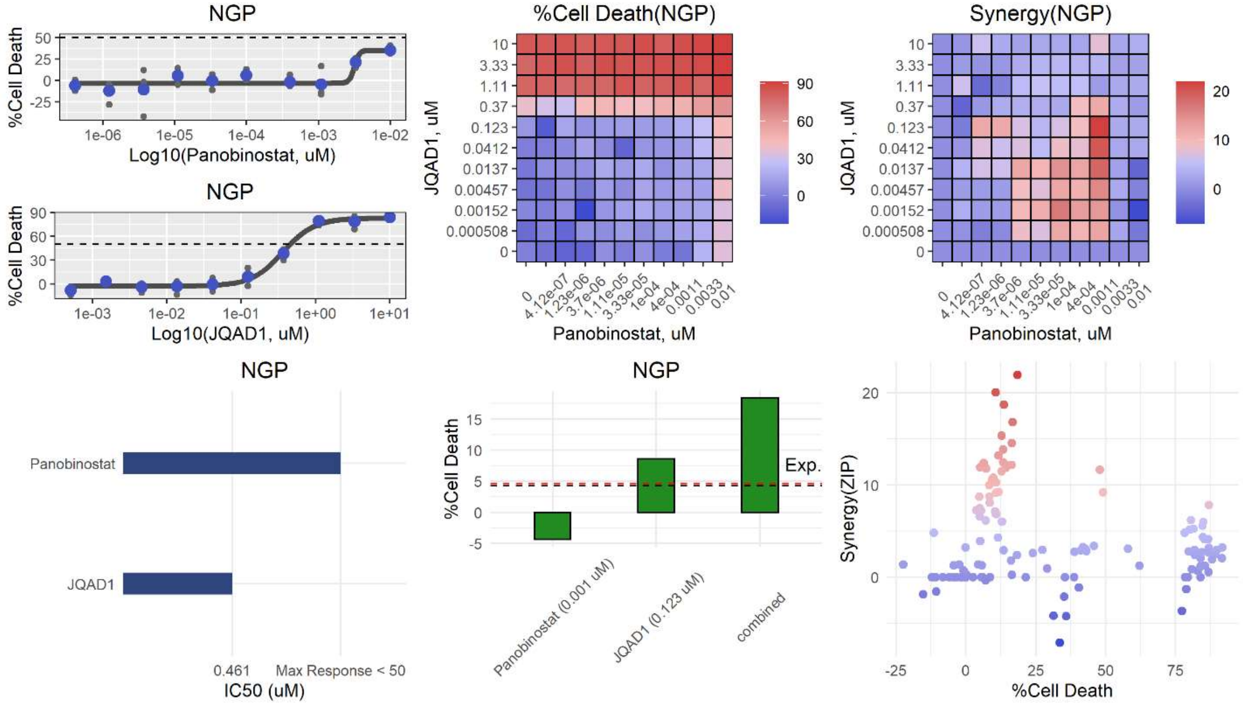
NGP
%Cell Death
Log10(Panobinostat, uM)
NGP
Log10(JQAD1, uM)
%Cell Death(NGP)
JQAD1, uM
Panobinostat, uM
Synergy(NGP)
NGP
Panobinostat
JQAD1
0.461
Max Response < 50
IC50 (uM)
NGP
Exp.
Panobinostat (0.001 uM)
JQAD1 (0.123 uM)
combined
Synergy(ZIP)
%Cell Death

**av**

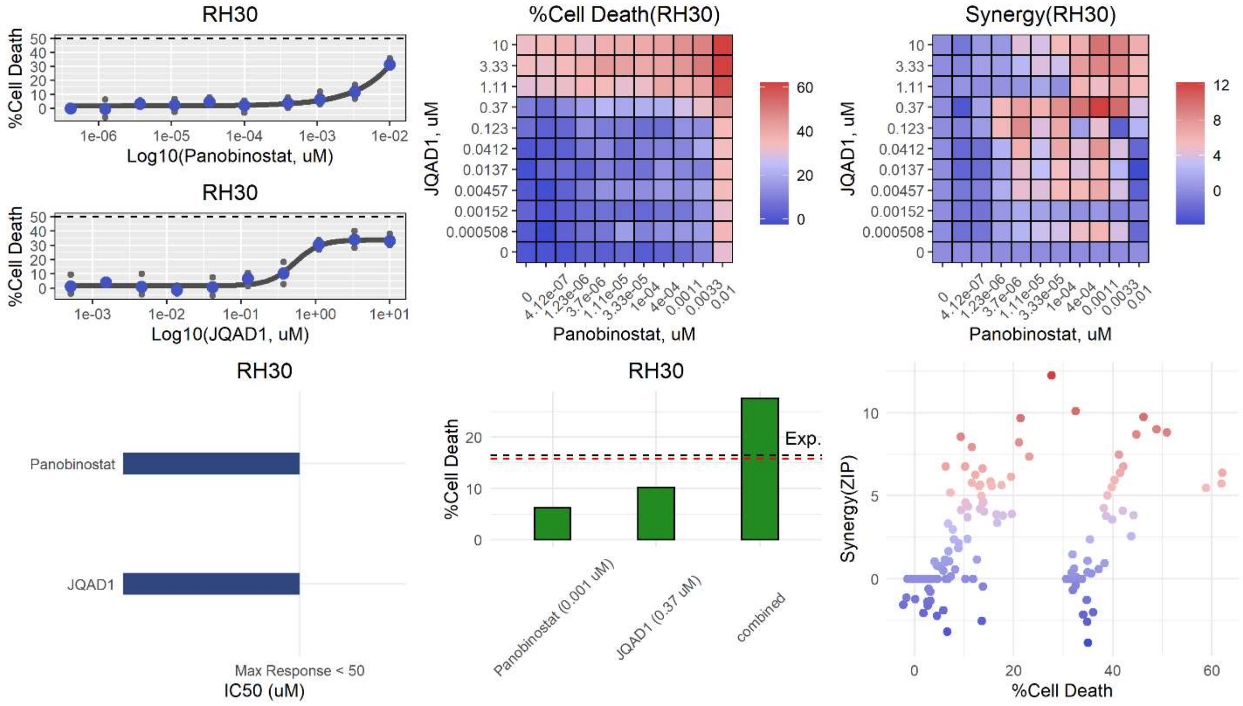
RH30
%Cell Death
Log10(Panobinostat, uM)
RH30
Log10(JQAD1, uM)
%Cell Death(RH30)
JQAD1, uM
Panobinostat, uM
Synergy(RH30)
RH30
Panobinostat
JQAD1
Max Response < 50
IC50 (uM)
RH30
Exp.
Panobinostat (0.001 uM)
JQAD1 (0.37 uM)
combined
Synergy(ZIP)
%Cell Death

**aw**

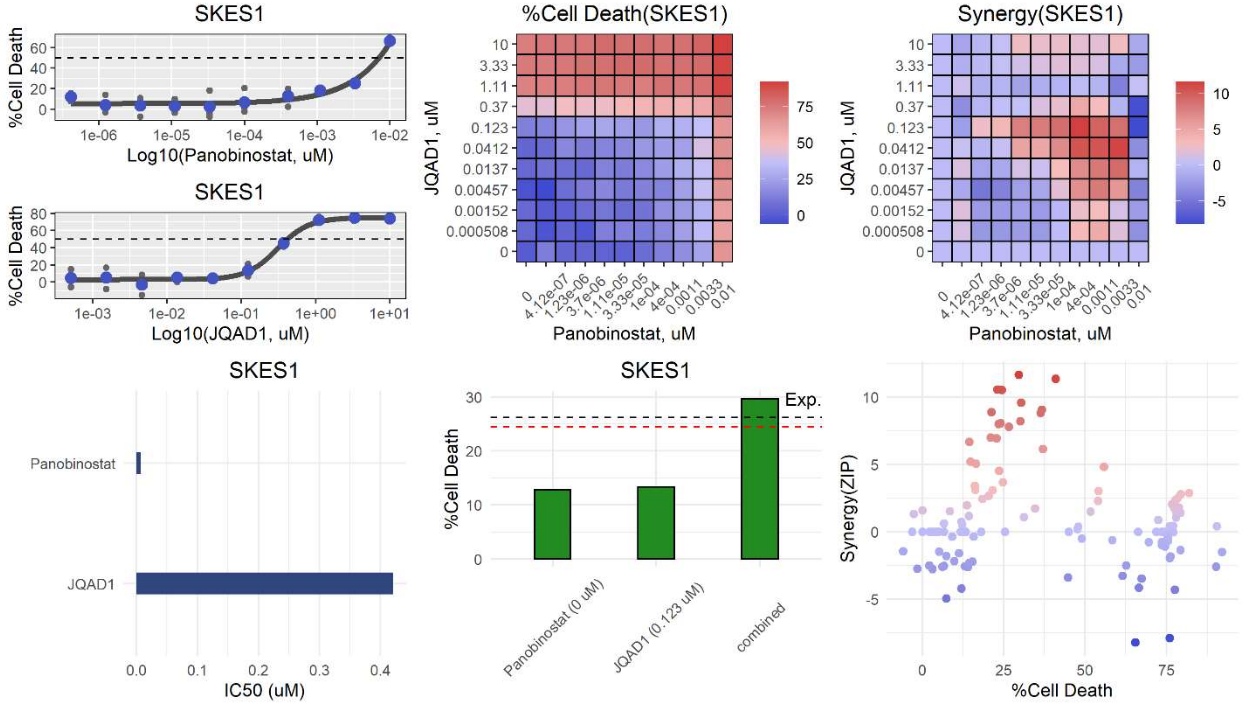
SKES1
%Cell Death
Log10(Panobinostat, uM)
SKES1
Log10(JQAD1, uM)
%Cell Death(SKES1)
JQAD1, uM
Panobinostat, uM
Synergy(SKES1)
SKES1
Panobinostat
JQAD1
IC50 (uM)
SKES1
Exp.
Panobinostat (0 uM)
JQAD1 (0.123 uM)
combined
Synergy(ZIP)
%Cell Death

**ax**

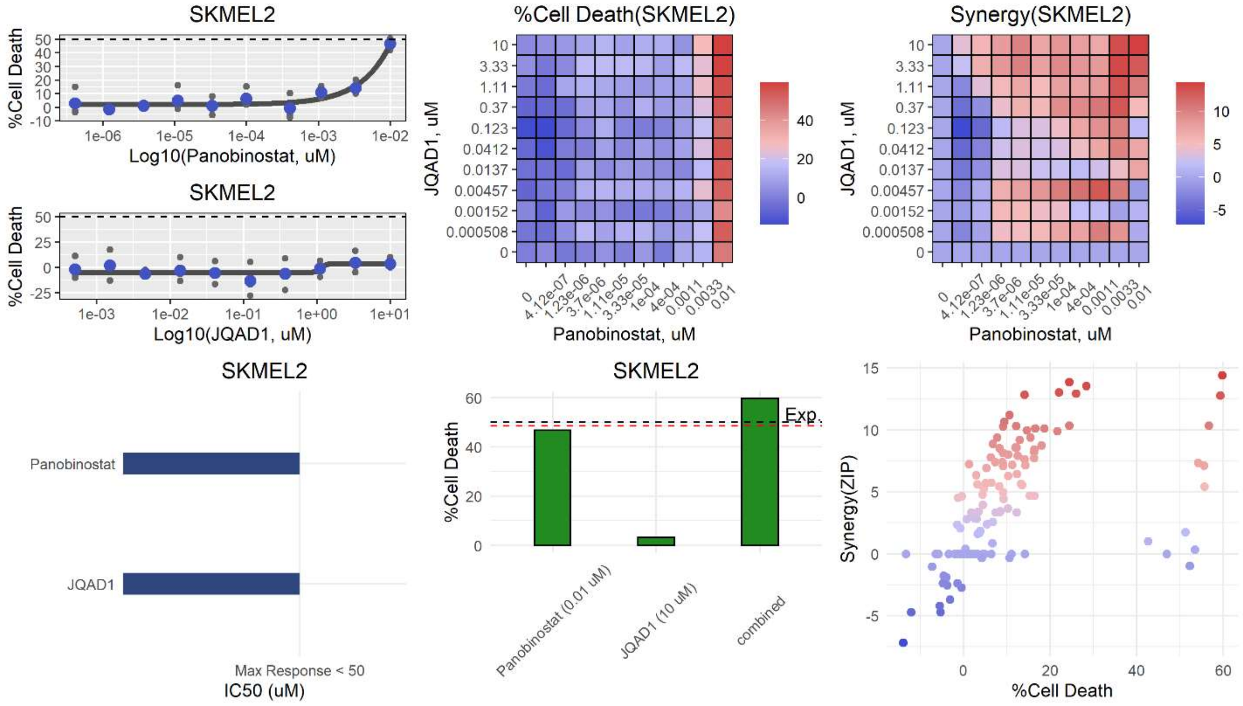
SKMEL2
%Cell Death
Log10(Panobinostat, uM)
SKMEL2
Log10(JQAD1, uM)
%Cell Death(SKMEL2)
JQAD1, uM
Panobinostat, uM
Synergy(SKMEL2)
SKMEL2
Panobinostat
JQAD1
Max Response < 50
IC50 (uM)
SKMEL2
Exp.
Panobinostat (0.01 uM)
JQAD1 (10 uM)
combined
Synergy(ZIP)
%Cell Death

**ay**

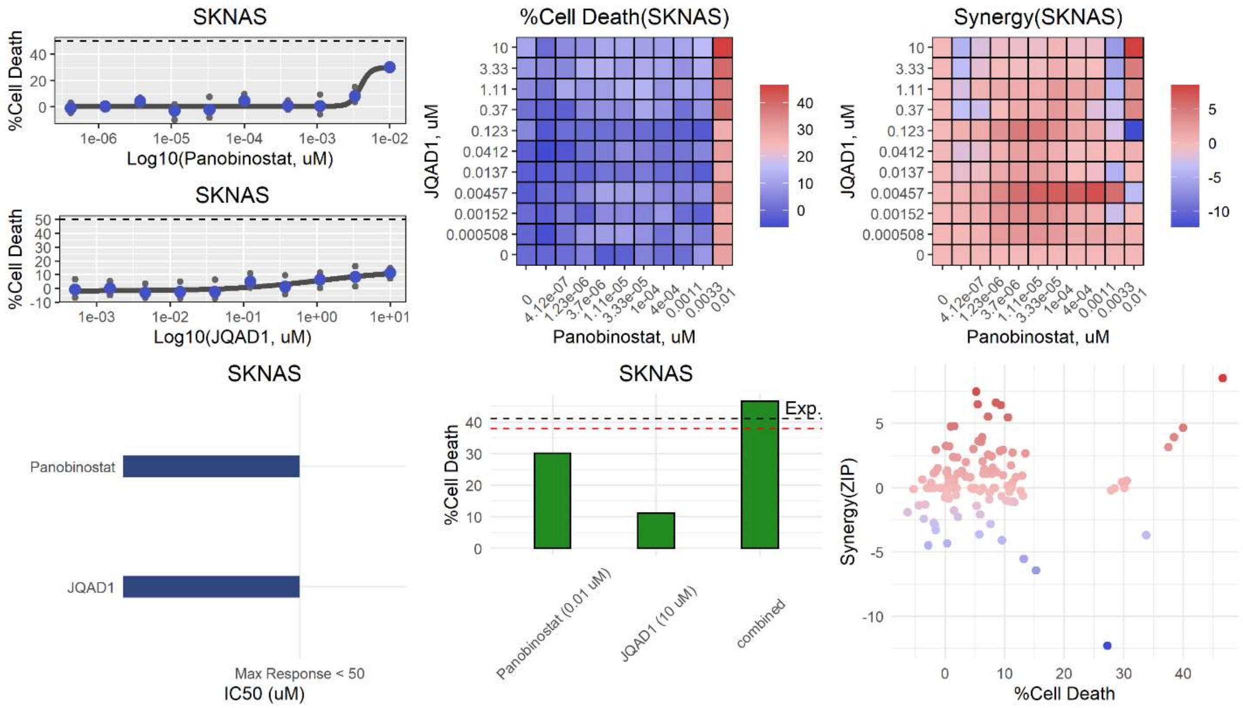
SKNAS
%Cell Death
Log10(Panobinostat, uM)
SKNAS
%Cell Death
Log10(JQAD1, uM)
%Cell Death(SKNAS)
JQAD1, uM
Panobinostat, uM
Synergy(SKNAS)
JQAD1, uM
Panobinostat, uM
SKNAS
Panobinostat
JQAD1
Max Response < 50
IC50 (uM)
SKNAS
%Cell Death
Exp.
Panobinostat (0.01 uM)
JQAD1 (10 uM)
combined
Synergy(ZIP)
%Cell Death

**az**

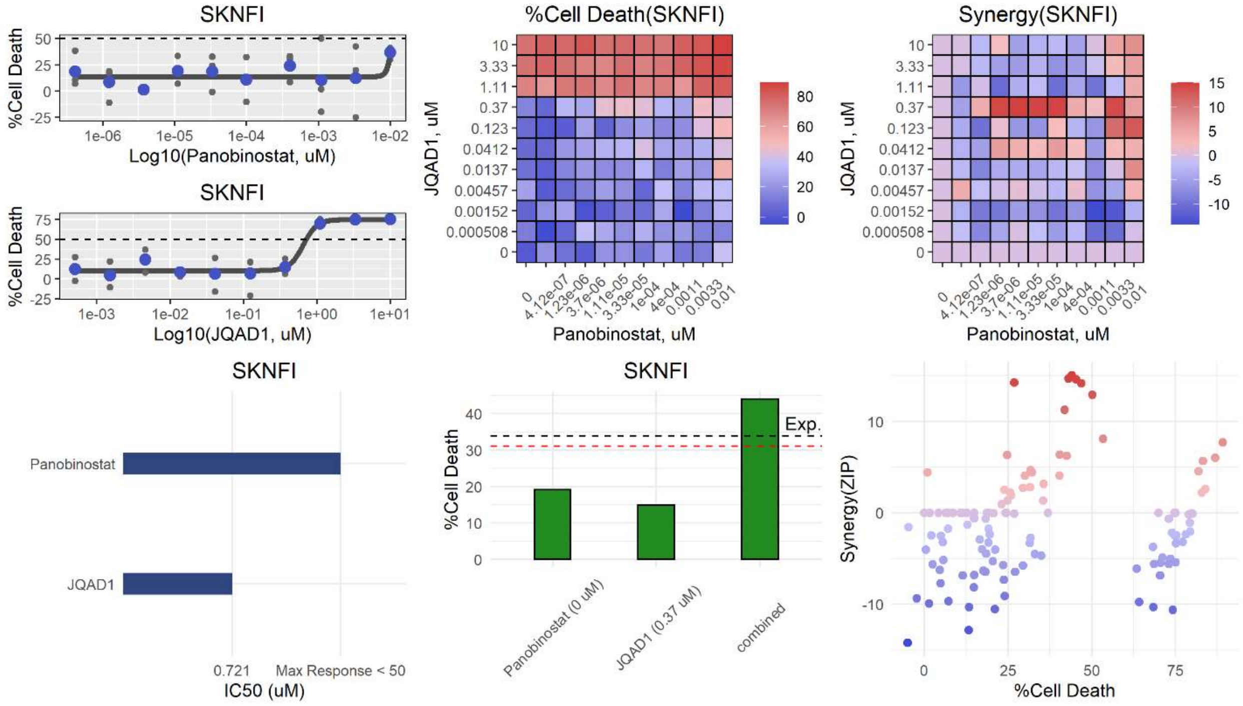
SKNFI
%Cell Death
Log10(Panobinostat, uM)
SKNFI
%Cell Death
Log10(JQAD1, uM)
%Cell Death(SKNFI)
JQAD1, uM
Panobinostat, uM
Synergy(SKNFI)
JQAD1, uM
Panobinostat, uM
SKNFI
Panobinostat
JQAD1
0.721
Max Response < 50
IC50 (uM)
SKNFI
%Cell Death
Exp.
Panobinostat (0 uM)
JQAD1 (0.37 uM)
combined
Synergy(ZIP)
%Cell Death

ba

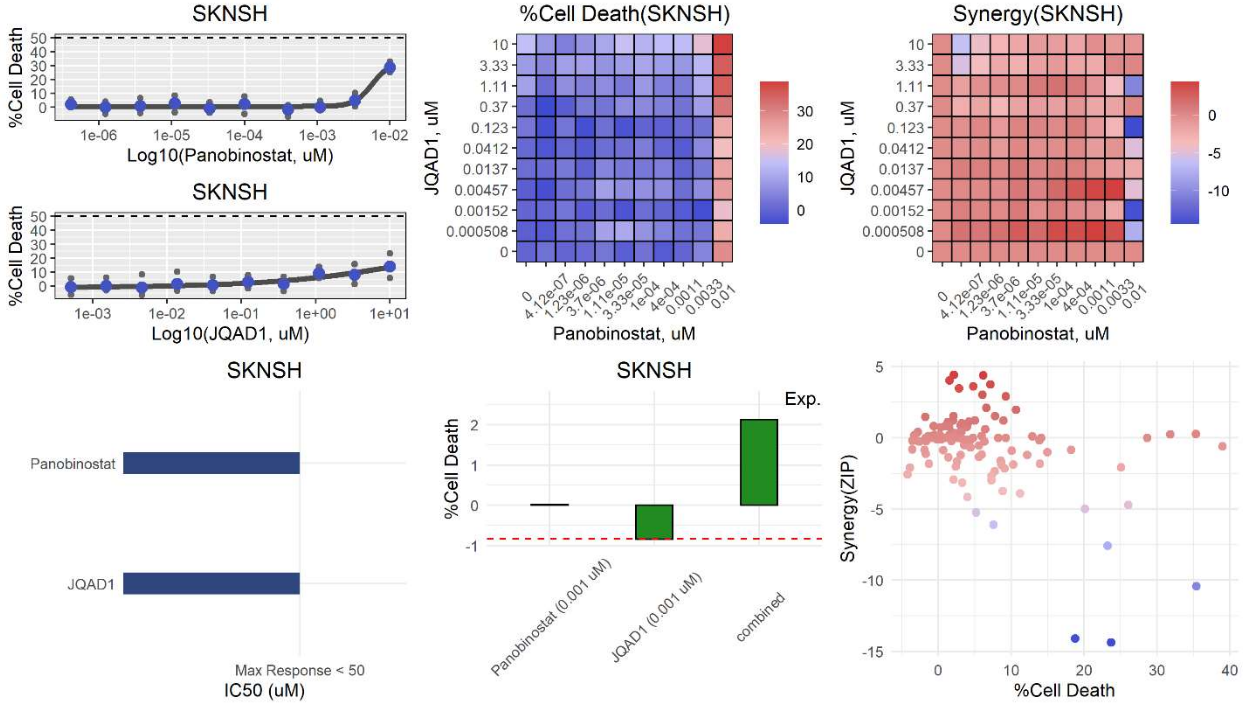
SKNSH
%Cell Death
Log10(Panobinostat, uM)
SKNSH
Log10(JQAD1, uM)
%Cell Death(SKNSH)
JQAD1, uM
Panobinostat, uM
Synergy(SKNSH)
SKNSH
Panobinostat
JQAD1
Max Response < 50
IC50 (uM)
SKNSH
Exp.
Panobinostat (0.001 uM)
JQAD1 (0.001 uM)
combined
Synergy(ZIP)
%Cell Death

bc

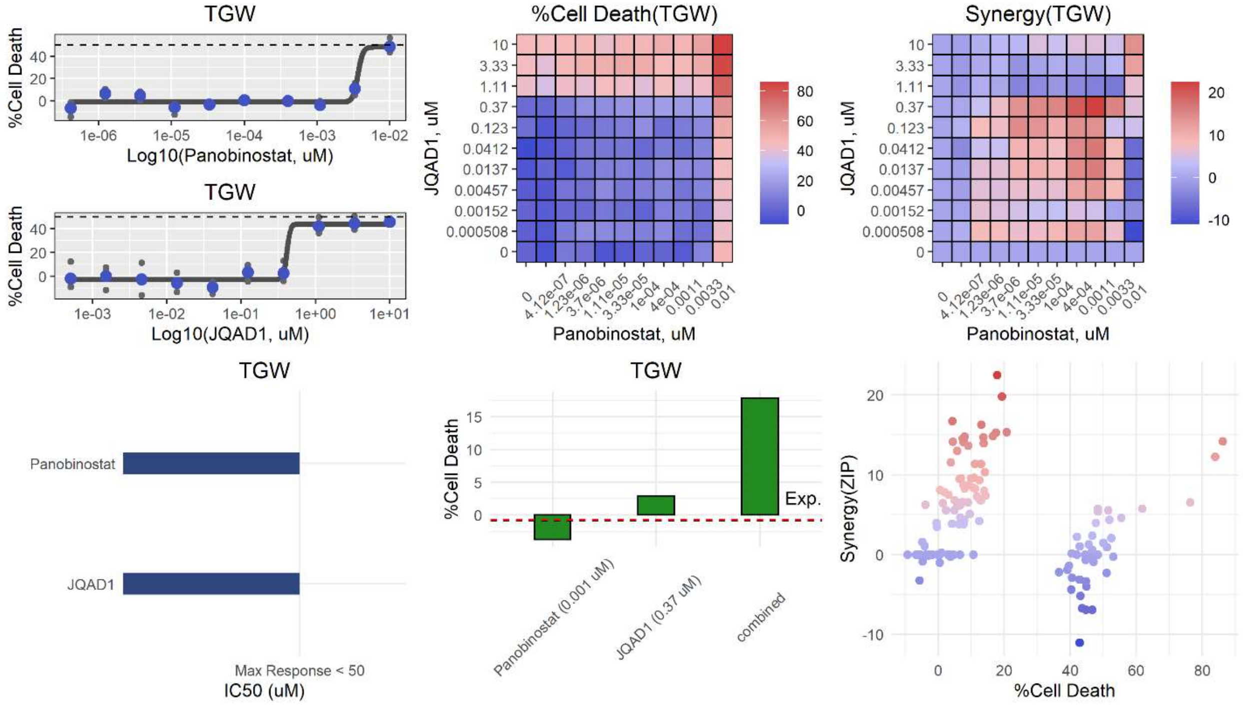
TGW
%Cell Death
Log10(Panobinostat, uM)
TGW
Log10(JQAD1, uM)
%Cell Death(TGW)
JQAD1, uM
Panobinostat, uM
Synergy(TGW)
TGW
Panobinostat
JQAD1
Max Response < 50
IC50 (uM)
TGW
Exp.
Panobinostat (0.001 uM)
JQAD1 (0.37 uM)
combined
Synergy(ZIP)
%Cell Death

**bd**

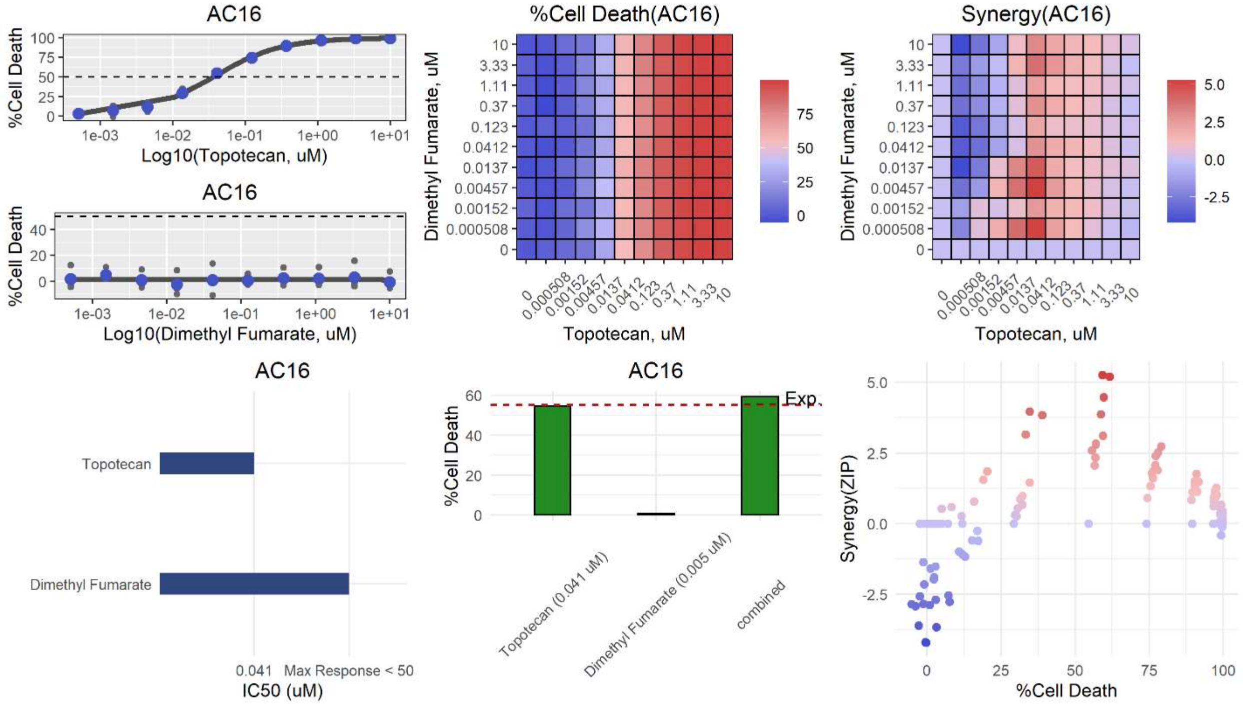

AC16
%Cell Death
Log10(Topotecan, uM)
AC16
%Cell Death
Log10(Dimethyl Fumarate, uM)
%Cell Death(AC16)
Dimethyl Fumarate, uM
Topotecan, uM
Synergy(AC16)
Dimethyl Fumarate, uM
Topotecan, uM
AC16
Topotecan
Dimethyl Fumarate
0.041 Max Response < 50
IC50 (uM)
AC16
%Cell Death
Exp.
Topotecan (0.041 uM)
Dimethyl Fumarate (0.005 uM)
combined
Synergy(ZIP)
%Cell Death


**be**

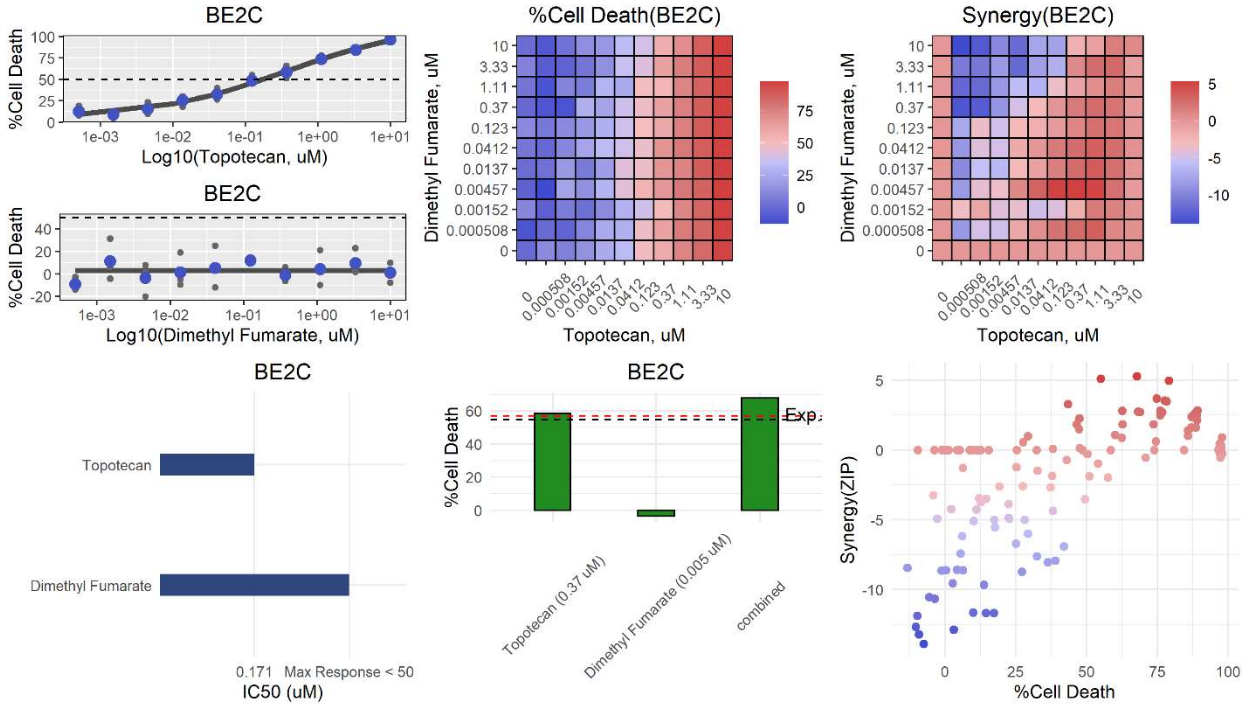

BE2C
%Cell Death
Log10(Topotecan, uM)
BE2C
%Cell Death
Log10(Dimethyl Fumarate, uM)
%Cell Death(BE2C)
Dimethyl Fumarate, uM
Topotecan, uM
Synergy(BE2C)
Dimethyl Fumarate, uM
Topotecan, uM
BE2C
Topotecan
Dimethyl Fumarate
0.171 Max Response < 50
IC50 (uM)
BE2C
%Cell Death
Exp.
Topotecan (0.37 uM)
Dimethyl Fumarate (0.005 uM)
combined
Synergy(ZIP)
%Cell Death

**bf**

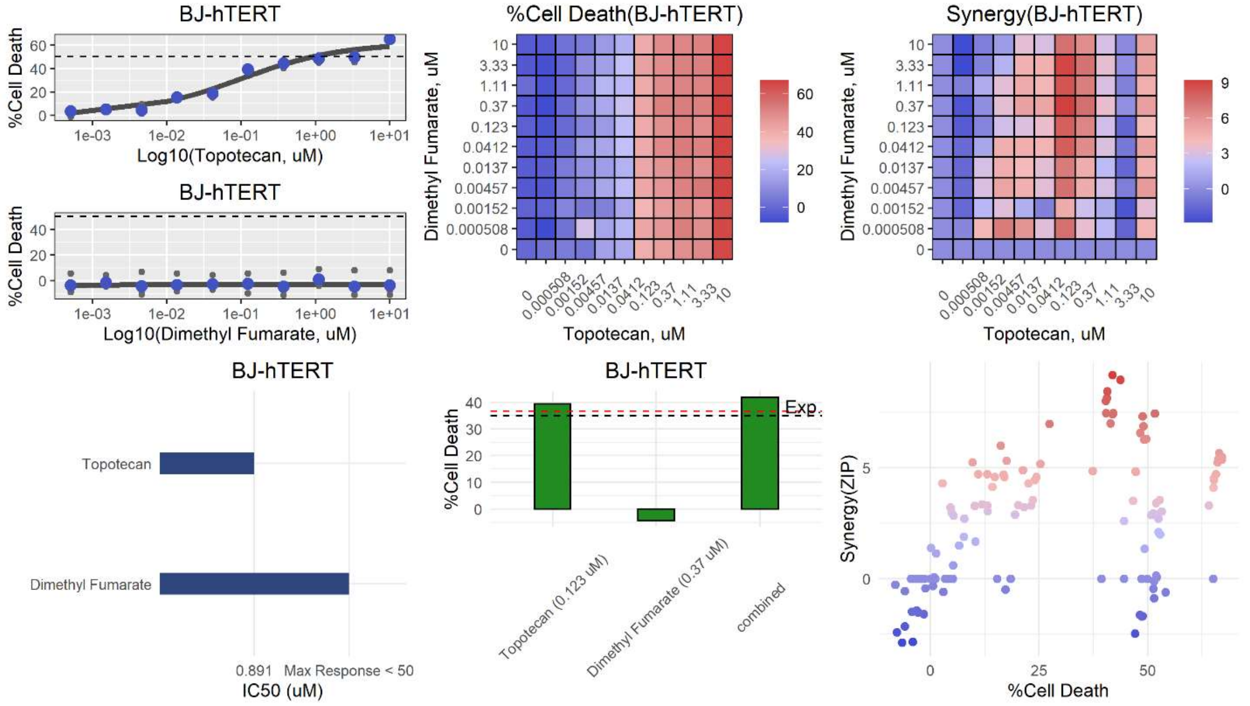
BJ-hTERT
%Cell Death
Log10(Topotecan, uM)
BJ-hTERT
%Cell Death
Log10(Dimethyl Fumarate, uM)
%Cell Death(BJ-hTERT)
Dimethyl Fumarate, uM
Topotecan, uM
Synergy(BJ-hTERT)
Dimethyl Fumarate, uM
Topotecan, uM
BJ-hTERT
Topotecan
Dimethyl Fumarate
0.891 Max Response < 50
IC50 (uM)
BJ-hTERT
%Cell Death
Exp.
Topotecan (0.123 uM)
Dimethyl Fumarate (0.37 uM)
combined
Synergy(ZIP)
%Cell Death

**bg**

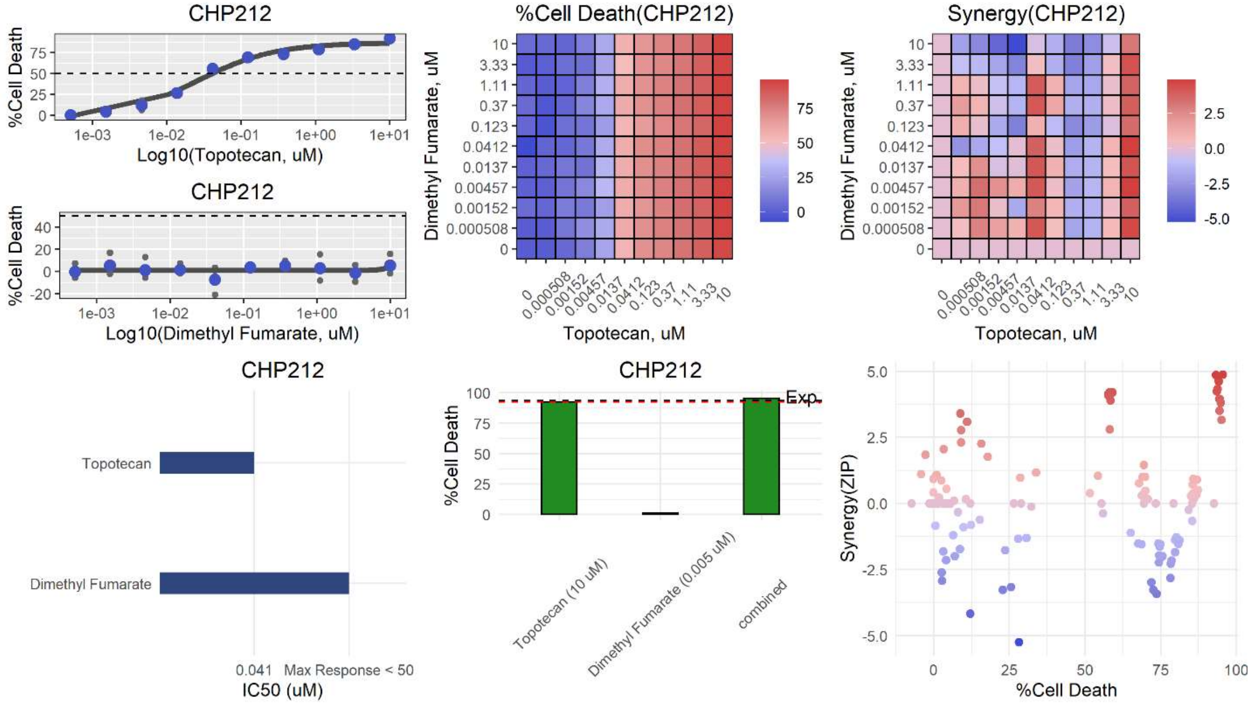
CHP212
%Cell Death
Log10(Topotecan, uM)
CHP212
%Cell Death
Log10(Dimethyl Fumarate, uM)
%Cell Death(CHP212)
Dimethyl Fumarate, uM
Topotecan, uM
Synergy(CHP212)
Dimethyl Fumarate, uM
Topotecan, uM
CHP212
Topotecan
Dimethyl Fumarate
0.041 Max Response < 50
IC50 (uM)
CHP212
%Cell Death
Exp.
Topotecan (10 uM)
Dimethyl Fumarate (0.005 uM)
combined
Synergy(ZIP)
%Cell Death

**bh**

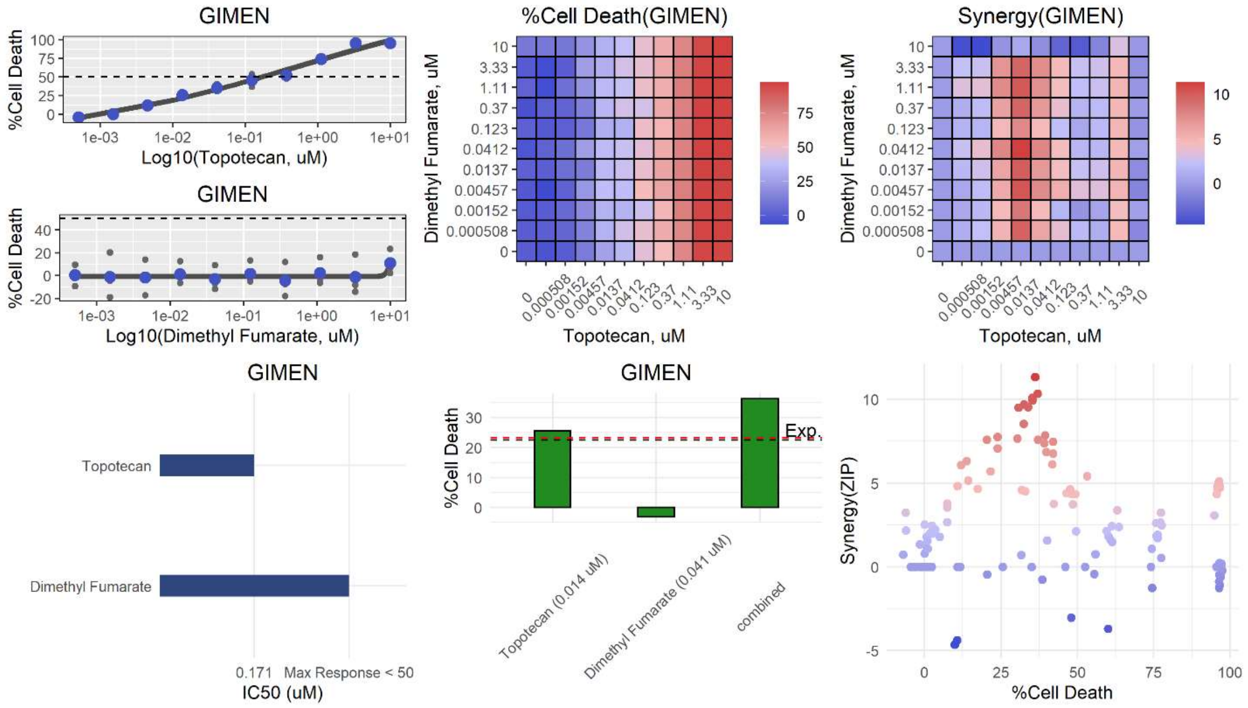
GIMEN
%Cell Death
Log10(Topotecan, uM)
GIMEN
%Cell Death
Log10(Dimethyl Fumarate, uM)
%Cell Death(GIMEN)
Dimethyl Fumarate, uM
Topotecan, uM
Synergy(GIMEN)
Dimethyl Fumarate, uM
Topotecan, uM
GIMEN
Topotecan
Dimethyl Fumarate
0.171 Max Response < 50
IC50 (uM)
GIMEN
%Cell Death
Exp.
Topotecan (0.014 uM)
Dimethyl Fumarate (0.041 uM)
combined
Synergy(ZIP)
%Cell Death

**bi**

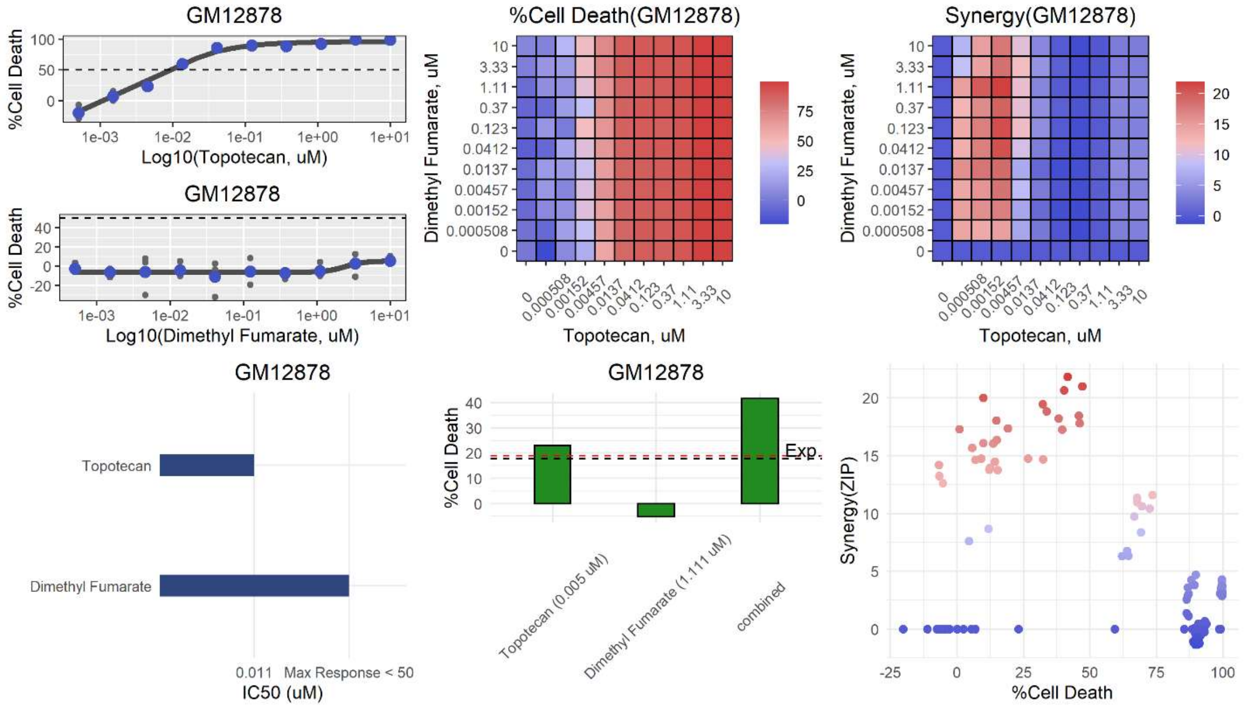
GM12878
%Cell Death
Log10(Topotecan, uM)
GM12878
%Cell Death
Log10(Dimethyl Fumarate, uM)
%Cell Death(GM12878)
Dimethyl Fumarate, uM
Topotecan, uM
Synergy(GM12878)
Dimethyl Fumarate, uM
Topotecan, uM
GM12878
Topotecan
Dimethyl Fumarate
0.011 Max Response < 50
IC50 (uM)
GM12878
%Cell Death
Exp.
Topotecan (0.005 uM)
Dimethyl Fumarate (1.111 uM)
combined
Synergy(ZIP)
%Cell Death

**bj**

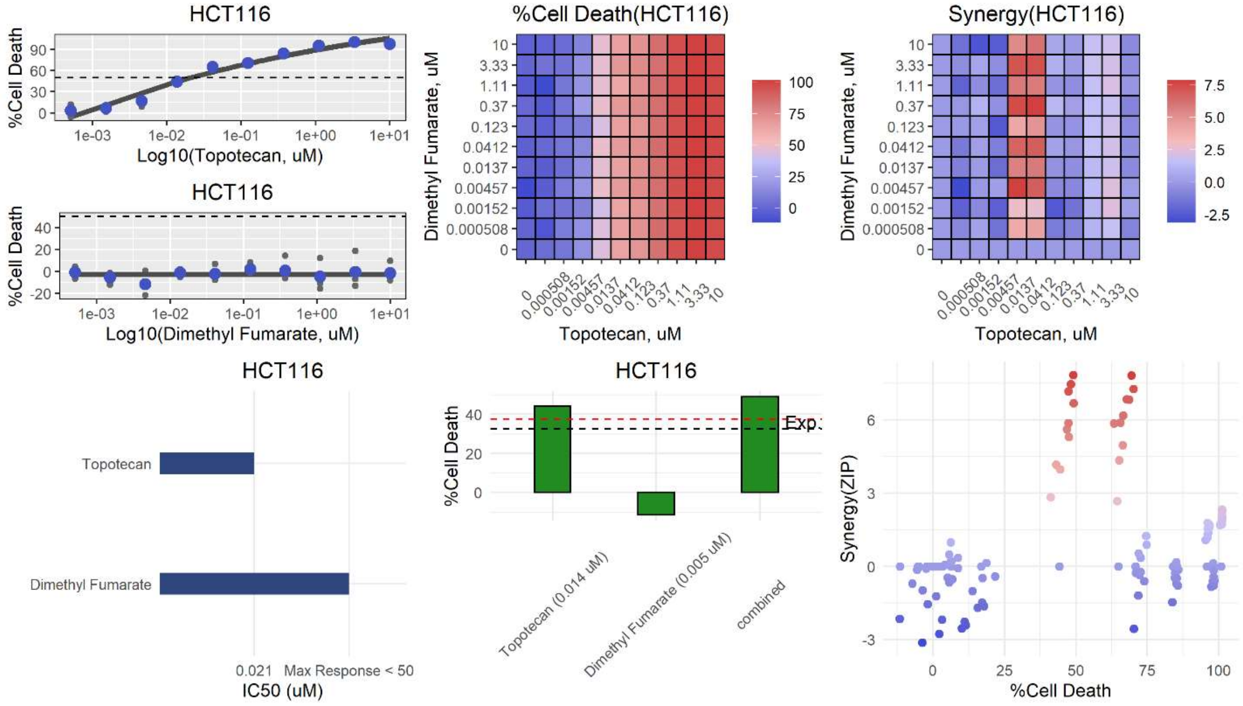
HCT116
%Cell Death
Log10(Topotecan, uM)
HCT116
%Cell Death
Log10(Dimethyl Fumarate, uM)
%Cell Death(HCT116)
Dimethyl Fumarate, uM
Topotecan, uM
Synergy(HCT116)
Dimethyl Fumarate, uM
Topotecan, uM
HCT116
Topotecan
Dimethyl Fumarate
0.021 Max Response < 50
IC50 (uM)
HCT116
%Cell Death
Exp.
Topotecan (0.014 uM)
Dimethyl Fumarate (0.005 uM)
combined
Synergy(ZIP)
%Cell Death

**bk**

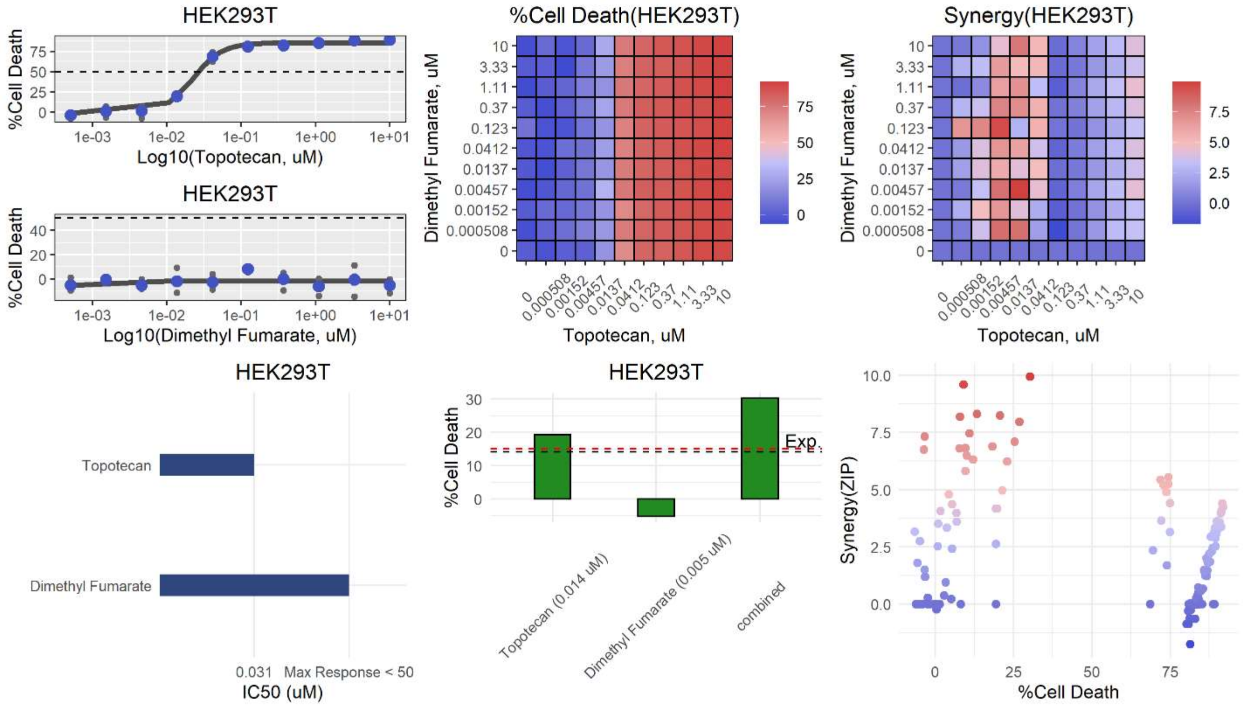
HEK293T
%Cell Death
Log10(Topotecan, uM)
HEK293T
%Cell Death
Log10(Dimethyl Fumarate, uM)
%Cell Death(HEK293T)
Dimethyl Fumarate, uM
Topotecan, uM
Synergy(HEK293T)
Dimethyl Fumarate, uM
Topotecan, uM
HEK293T
Topotecan
Dimethyl Fumarate
0.031 Max Response < 50
IC50 (uM)
HEK293T
%Cell Death
Exp.
Topotecan (0.014 uM)
Dimethyl Fumarate (0.005 uM)
combined
Synergy(ZIP)
%Cell Death

**bl**

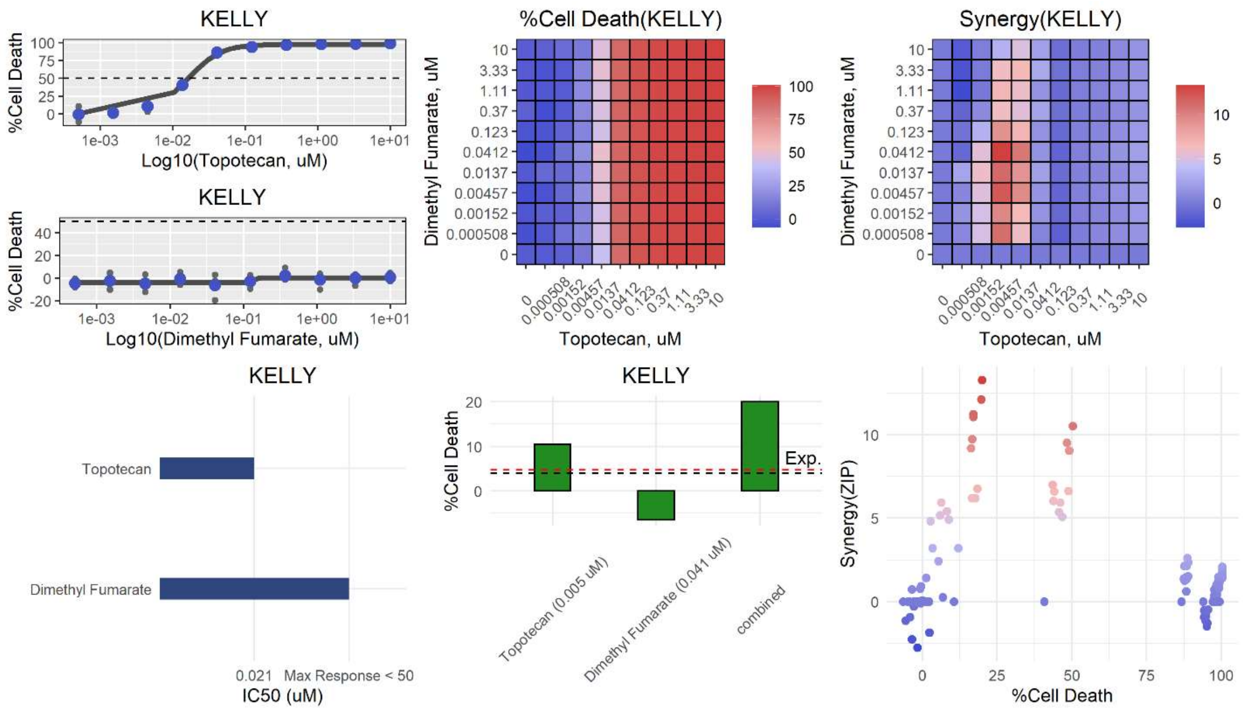
KELLY
%Cell Death
Log10(Topotecan, uM)
KELLY
%Cell Death
Log10(Dimethyl Fumarate, uM)
%Cell Death(KELLY)
Dimethyl Fumarate, uM
Topotecan, uM
Synergy(KELLY)
Dimethyl Fumarate, uM
Topotecan, uM
KELLY
Topotecan
Dimethyl Fumarate
0.021
Max Response < 50
IC50 (uM)
KELLY
%Cell Death
Exp.
Topotecan (0.005 uM)
Dimethyl Fumarate (0.041 uM)
combined
Synergy(ZIP)
%Cell Death

**bm**

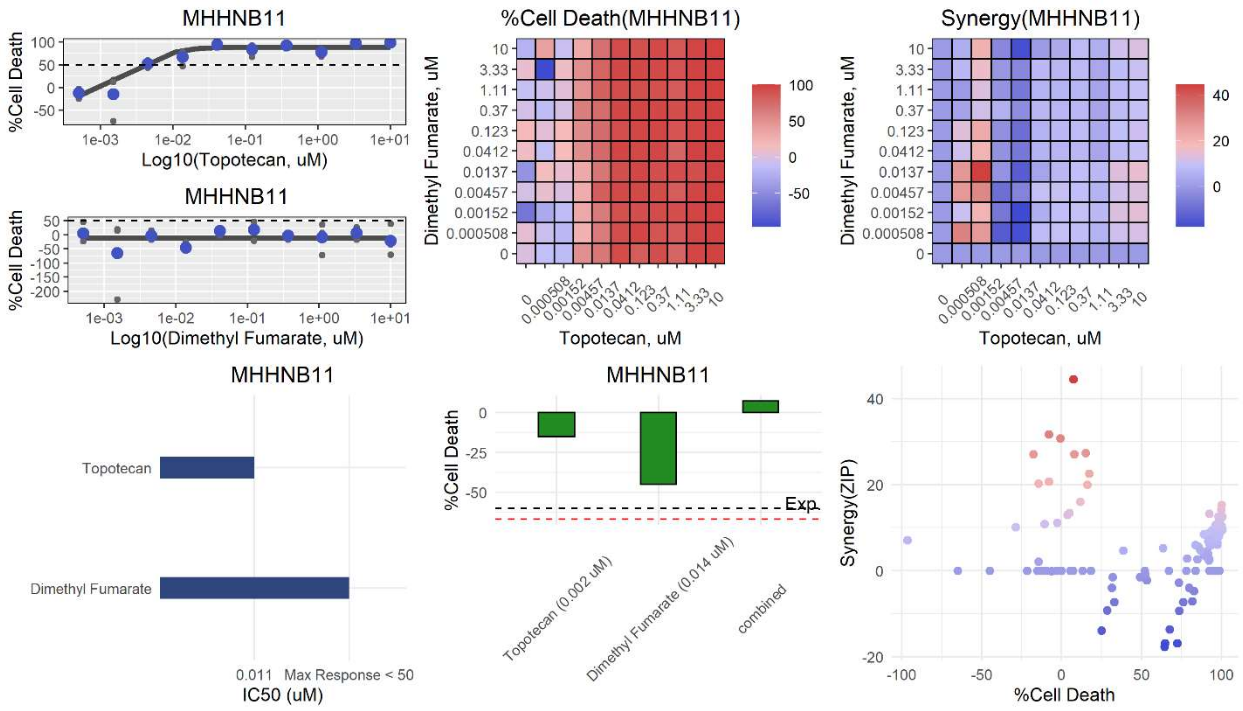
MHHNB11
%Cell Death
Log10(Topotecan, uM)
MHHNB11
%Cell Death
Log10(Dimethyl Fumarate, uM)
%Cell Death(MHHNB11)
Dimethyl Fumarate, uM
Topotecan, uM
Synergy(MHHNB11)
Dimethyl Fumarate, uM
Topotecan, uM
MHHNB11
Topotecan
Dimethyl Fumarate
0.011
Max Response < 50
IC50 (uM)
MHHNB11
%Cell Death
Exp.
Topotecan (0.002 uM)
Dimethyl Fumarate (0.014 uM)
combined
Synergy(ZIP)
%Cell Death

**bn**

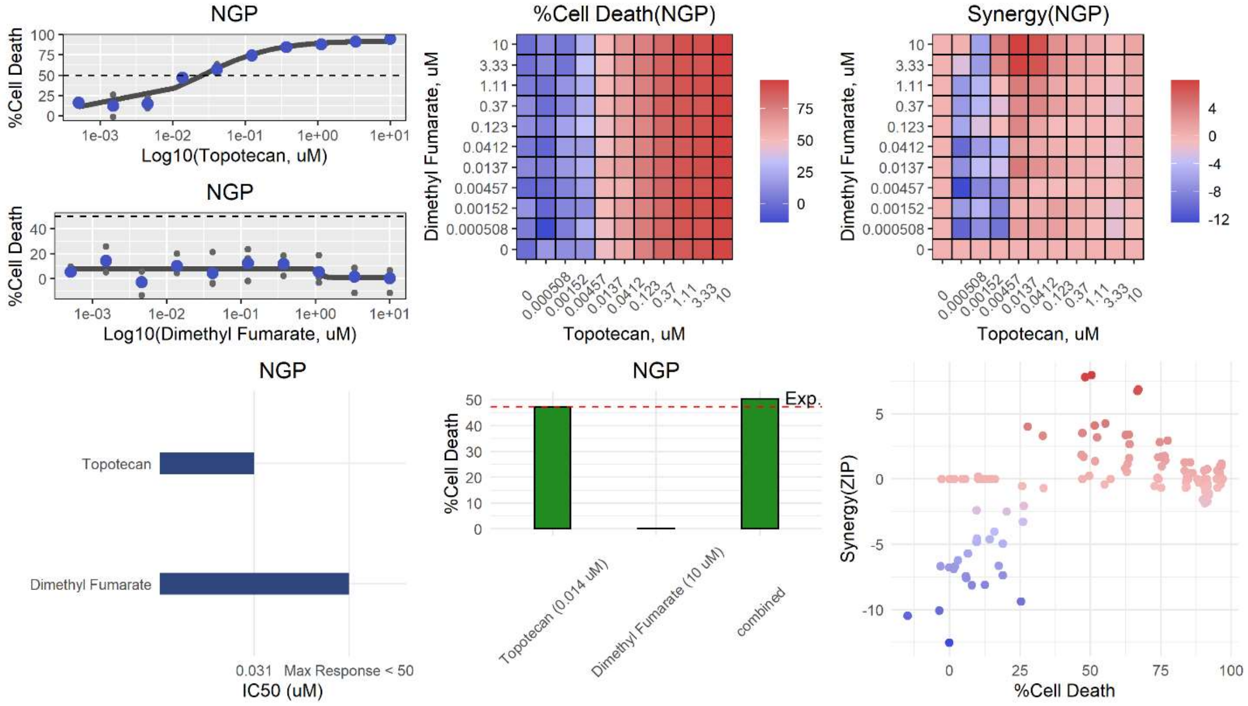
NGP
%Cell Death
Log10(Topotecan, uM)
NGP
%Cell Death
Log10(Dimethyl Fumarate, uM)
%Cell Death(NGP)
Dimethyl Fumarate, uM
Topotecan, uM
Synergy(NGP)
Dimethyl Fumarate, uM
Topotecan, uM
NGP
Topotecan
Dimethyl Fumarate
0.031
Max Response < 50
IC50 (uM)
NGP
%Cell Death
Exp.
Topotecan (0.014 uM)
Dimethyl Fumarate (10 uM)
combined
Synergy(ZIP)
%Cell Death

**bo**

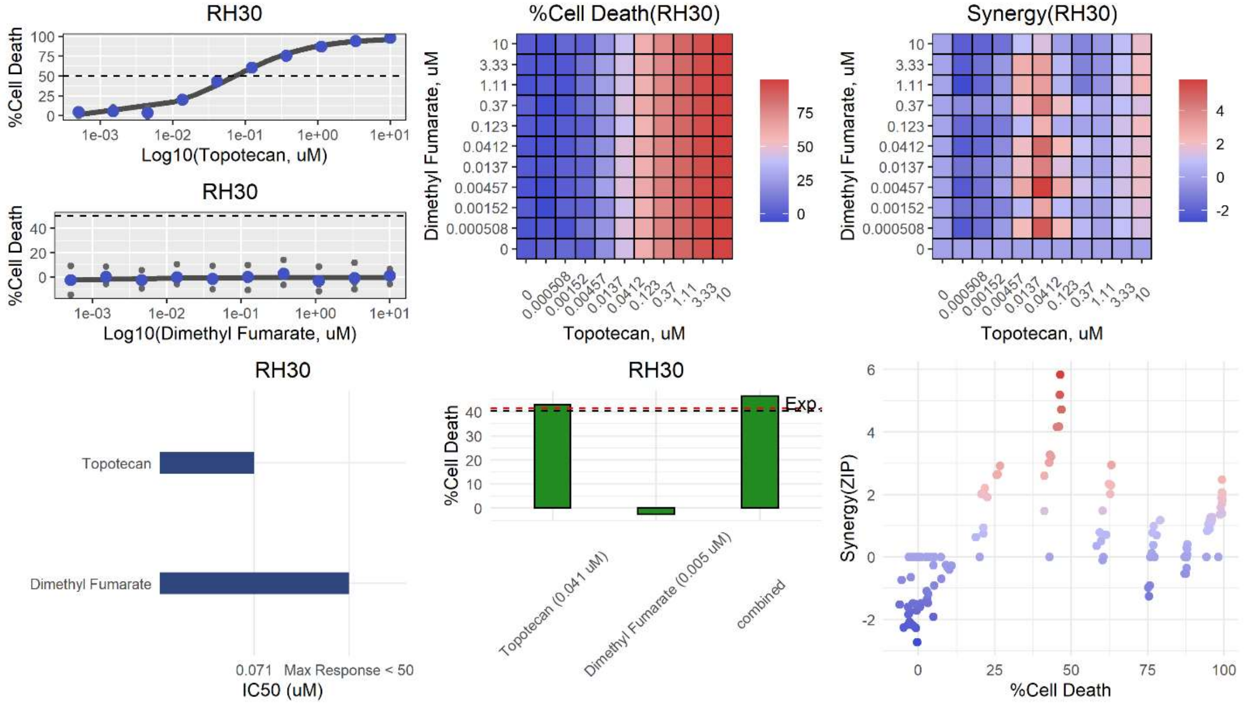
RH30
%Cell Death
Log10(Topotecan, uM)
RH30
%Cell Death
Log10(Dimethyl Fumarate, uM)
%Cell Death(RH30)
Dimethyl Fumarate, uM
Topotecan, uM
Synergy(RH30)
Dimethyl Fumarate, uM
Topotecan, uM
RH30
Topotecan
Dimethyl Fumarate
0.071
Max Response < 50
IC50 (uM)
RH30
%Cell Death
Exp.
Topotecan (0.041 uM)
Dimethyl Fumarate (0.005 uM)
combined
Synergy(ZIP)
%Cell Death

**bp**

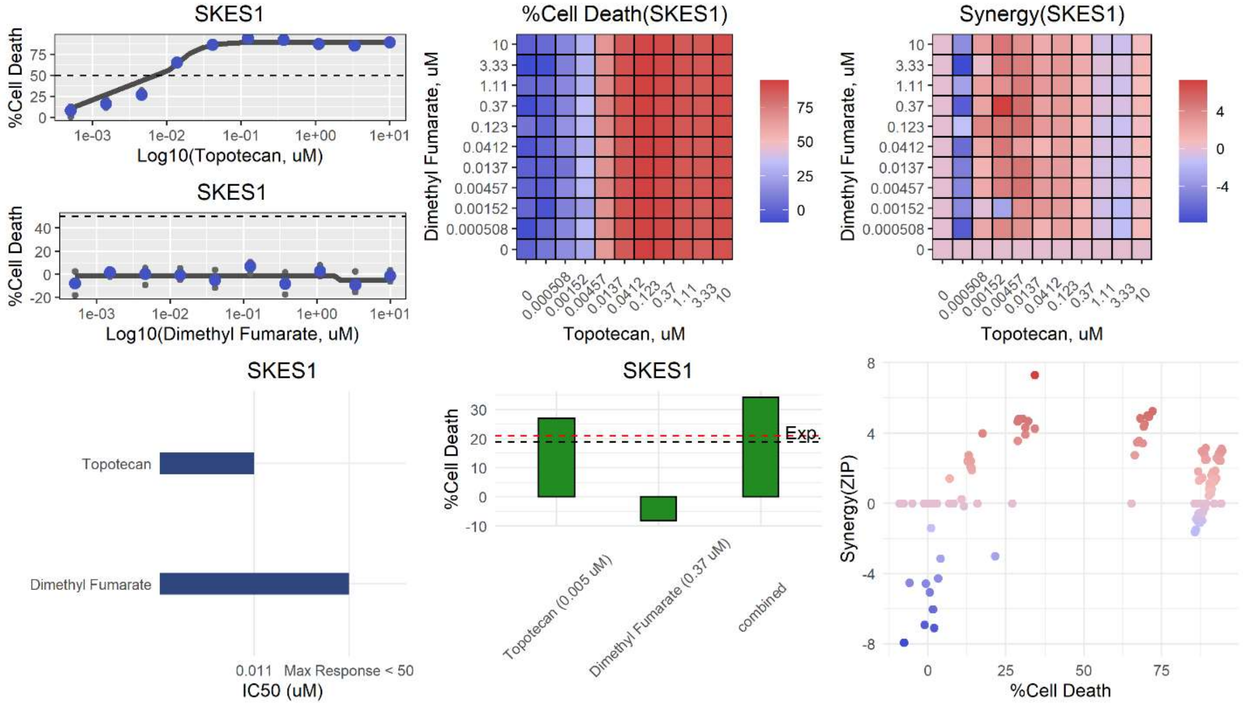

SKES1
%Cell Death
Log10(Topotecan, uM)
SKES1
%Cell Death
Log10(Dimethyl Fumarate, uM)
%Cell Death(SKES1)
Dimethyl Fumarate, uM
Topotecan, uM
Synergy(SKES1)
Dimethyl Fumarate, uM
Topotecan, uM
SKES1
Topotecan
Dimethyl Fumarate
0.011 Max Response < 50
IC50 (uM)
SKES1
%Cell Death
Exp.
Topotecan (0.005 uM)
Dimethyl Fumarate (0.37 uM)
combined
Synergy(ZIP)
%Cell Death


**bq**

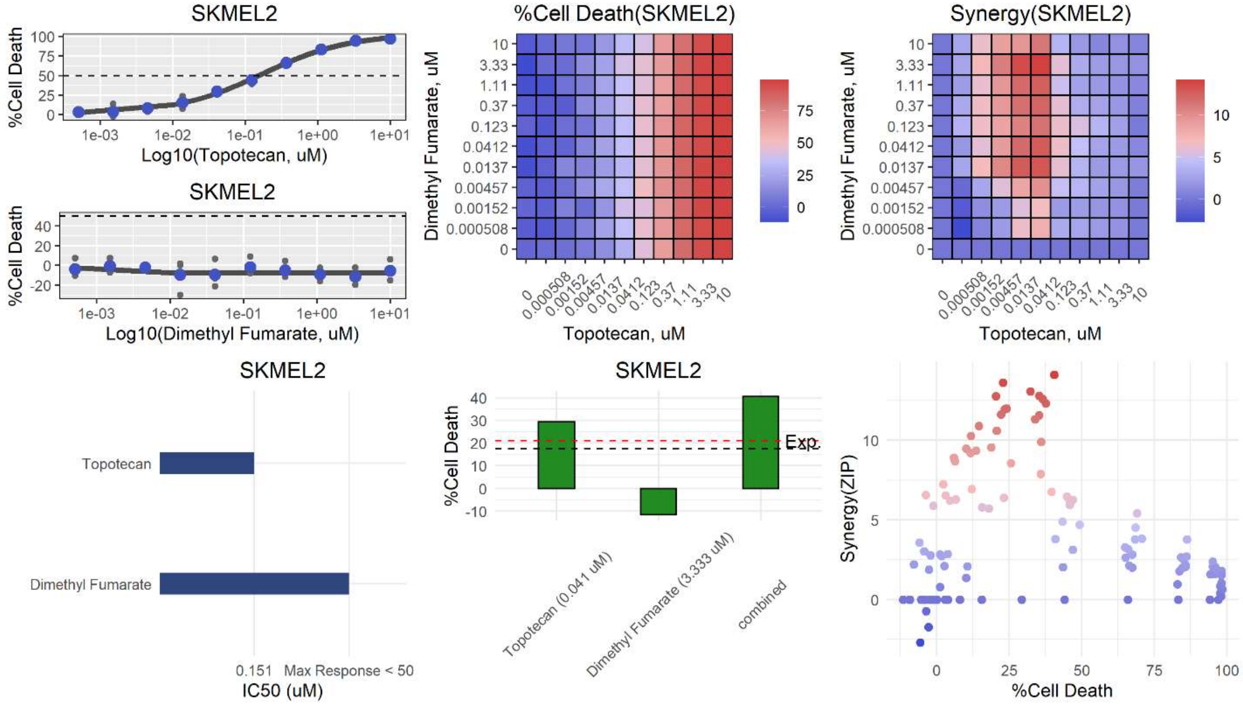

SKMEL2
%Cell Death
Log10(Topotecan, uM)
SKMEL2
%Cell Death
Log10(Dimethyl Fumarate, uM)
%Cell Death(SKMEL2)
Dimethyl Fumarate, uM
Topotecan, uM
Synergy(SKMEL2)
Dimethyl Fumarate, uM
Topotecan, uM
SKMEL2
Topotecan
Dimethyl Fumarate
0.151 Max Response < 50
IC50 (uM)
SKMEL2
%Cell Death
Exp.
Topotecan (0.041 uM)
Dimethyl Fumarate (3.333 uM)
combined
Synergy(ZIP)
%Cell Death

**br**

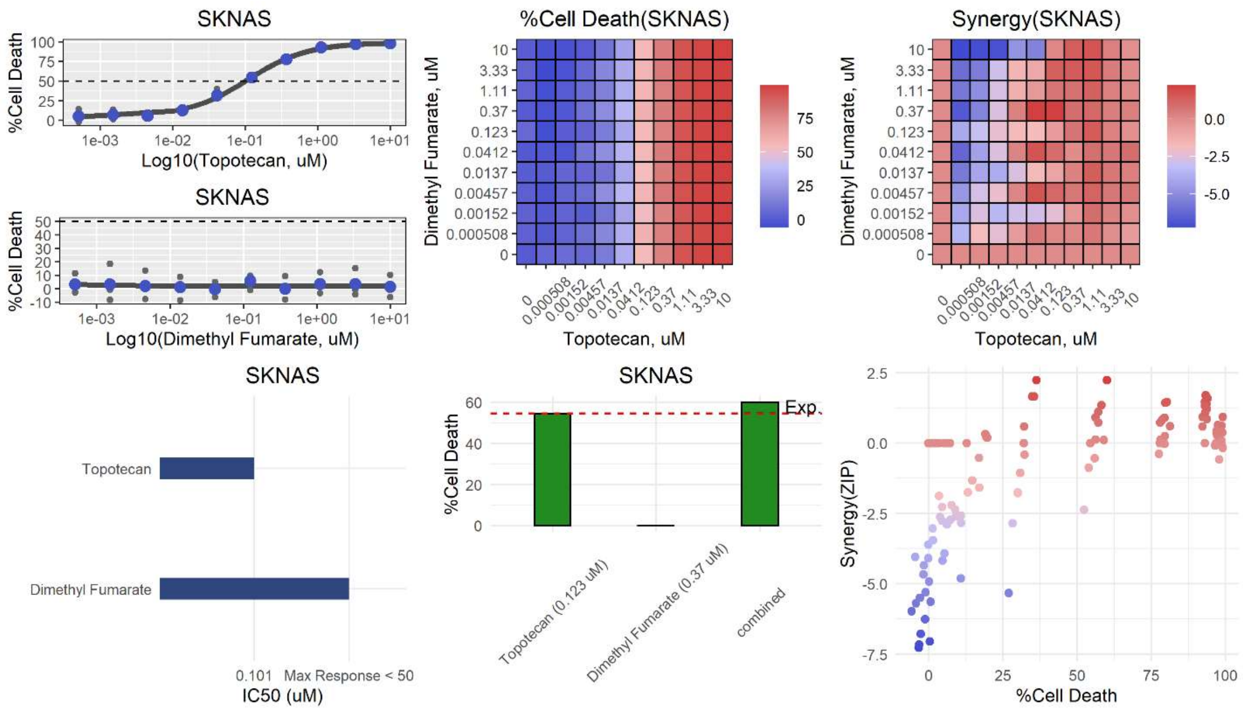
SKNAS
%Cell Death
Log10(Topotecan, uM)
SKNAS
%Cell Death
Log10(Dimethyl Fumarate, uM)
%Cell Death(SKNAS)
Dimethyl Fumarate, uM
Topotecan, uM
Synergy(SKNAS)
Dimethyl Fumarate, uM
Topotecan, uM
SKNAS
Topotecan
Dimethyl Fumarate
0.101
Max Response < 50
IC50 (uM)
SKNAS
%Cell Death
Exp.
Topotecan (0.123 uM)
Dimethyl Fumarate (0.37 uM)
combined
Synergy(ZIP)
%Cell Death

**bs**

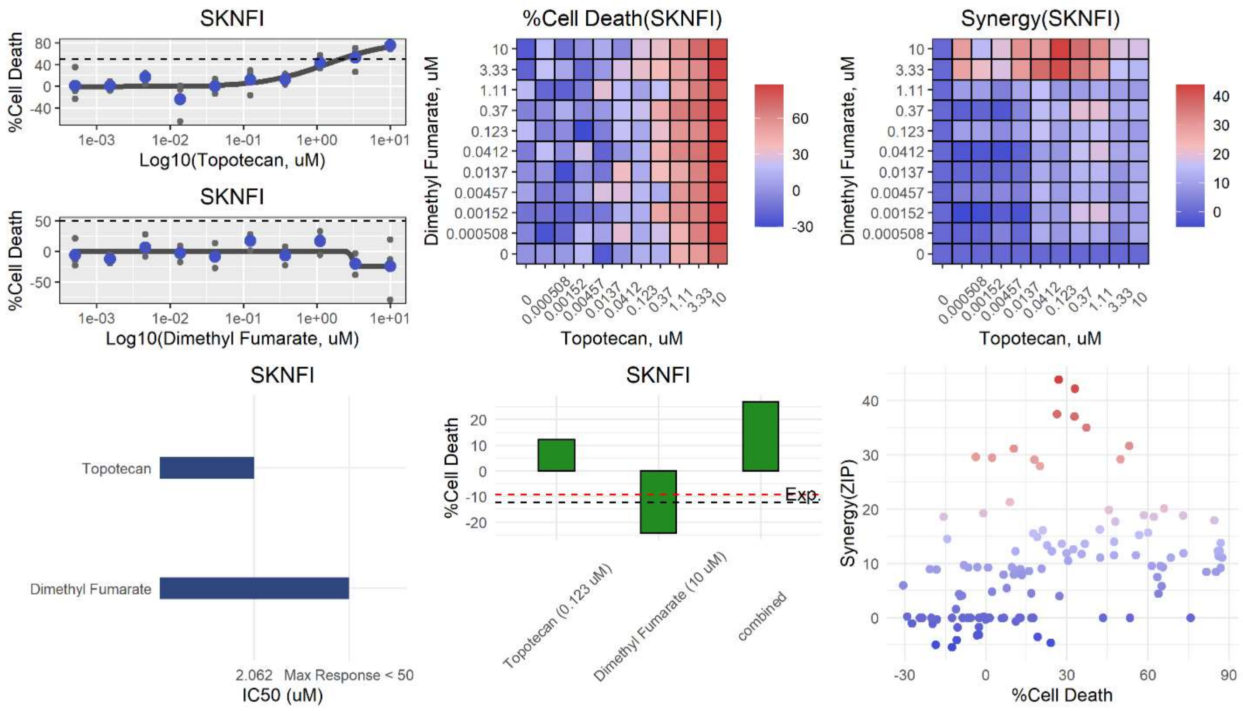
SKNFI
%Cell Death
Log10(Topotecan, uM)
SKNFI
%Cell Death
Log10(Dimethyl Fumarate, uM)
%Cell Death(SKNFI)
Dimethyl Fumarate, uM
Topotecan, uM
Synergy(SKNFI)
Dimethyl Fumarate, uM
Topotecan, uM
SKNFI
Topotecan
Dimethyl Fumarate
2.062
Max Response < 50
IC50 (uM)
SKNFI
%Cell Death
Exp.
Topotecan (0.123 uM)
Dimethyl Fumarate (10 uM)
combined
Synergy(ZIP)
%Cell Death

**bt**

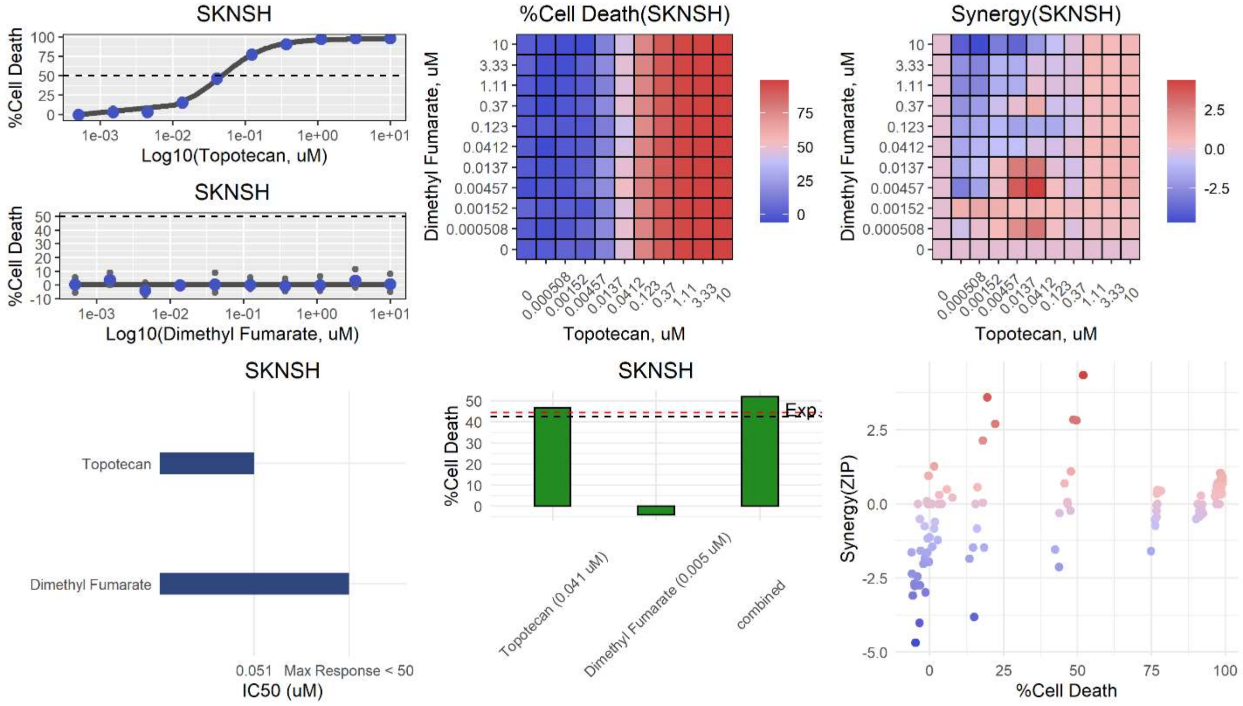
SKNSH
%Cell Death
Log10(Topotecan, uM)
SKNSH
%Cell Death
Log10(Dimethyl Fumarate, uM)
%Cell Death(SKNSH)
Dimethyl Fumarate, uM
Topotecan, uM
Synergy(SKNSH)
Dimethyl Fumarate, uM
Topotecan, uM
SKNSH
Topotecan
Dimethyl Fumarate
0.051
Max Response < 50
IC50 (uM)
SKNSH
%Cell Death
Exp.
Topotecan (0.041 uM)
Dimethyl Fumarate (0.005 uM)
combined
Synergy(ZIP)
%Cell Death

**bu**

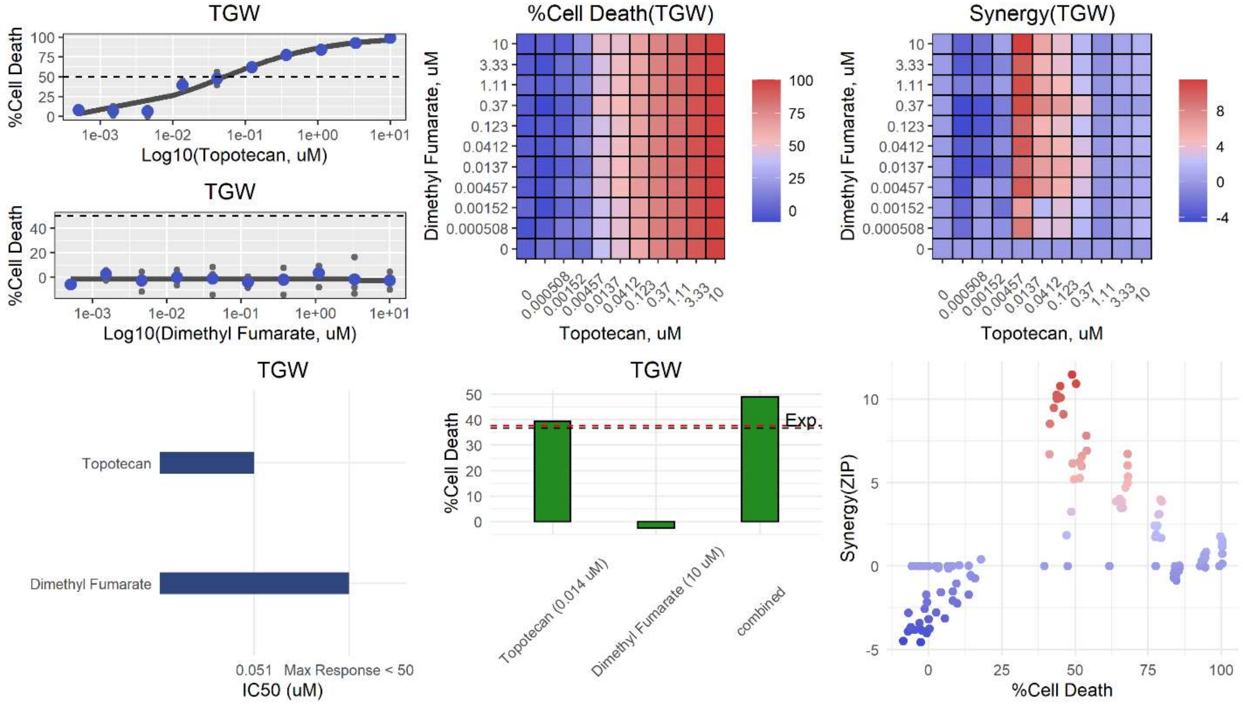
TGW
%Cell Death
Log10(Topotecan, uM)
TGW
%Cell Death
Log10(Dimethyl Fumarate, uM)
%Cell Death(TGW)
Dimethyl Fumarate, uM
Topotecan, uM
Synergy(TGW)
Dimethyl Fumarate, uM
Topotecan, uM
TGW
Topotecan
Dimethyl Fumarate
0.051
Max Response < 50
IC50 (uM)
TGW
%Cell Death
Exp.
Topotecan (0.014 uM)
Dimethyl Fumarate (10 uM)
combined
Synergy(ZIP)
%Cell Death

**Extended Data Figure 5. Summary of dense drug-drug combinations in all cell lines.**

72 panels for each combination of 4 sets in all 18 cell lines (a-bu) Dose-response curves for each single-agent (top left), normalized cell death matrix (top middle), synergy matrix (top right; Note: units are δZIP), $IC_{50}$ values of each single-agent (lower left), barplots corresponding to maximum synergy (lower middle; Note: dashed lines indicate expected values under additivity (black) or Bliss independence (red)), and scatterplot of cell death vs. synergy (lower right) for all 18 cell lines used in dense drug-drug screens.

**a**

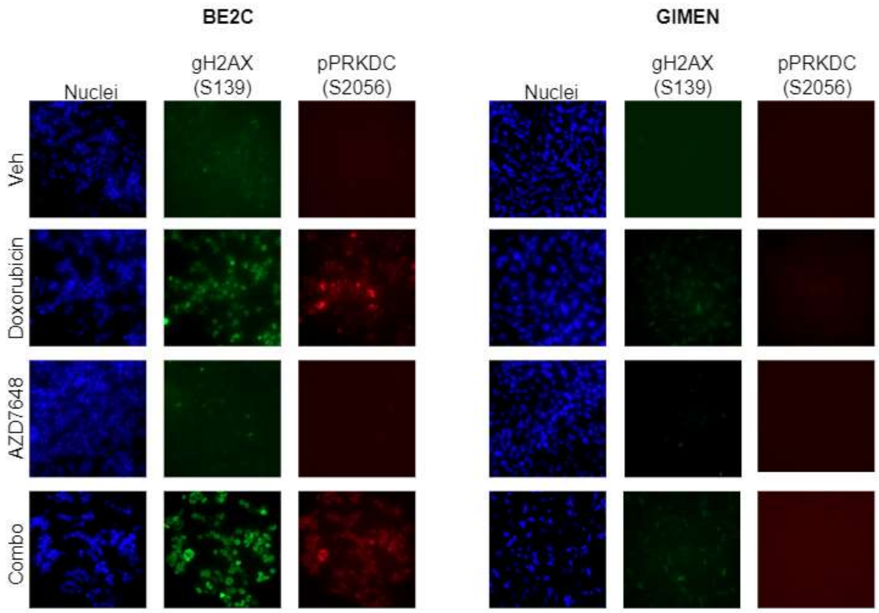


**b**

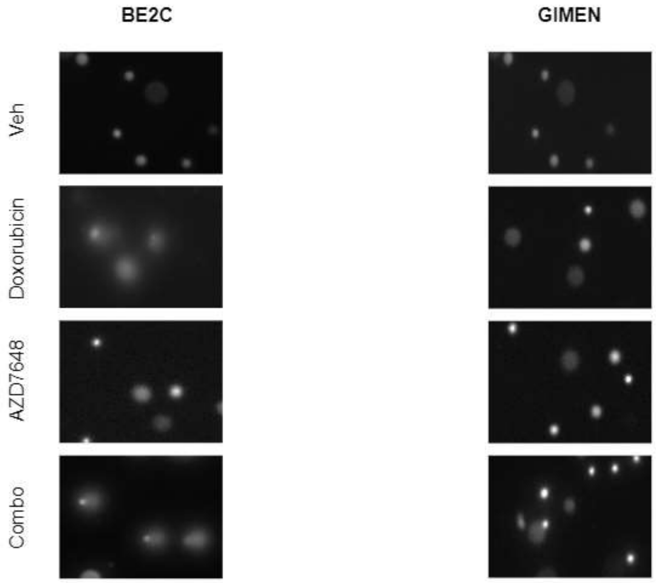


**Extended Data Figure 6. DNA damages in BE2C and GIMEN**

After 72 hr vehicle or drug treatment, γH2AX and phosphorylated PRKDC were visualized (a) and tail moment was visualized (b) a

b

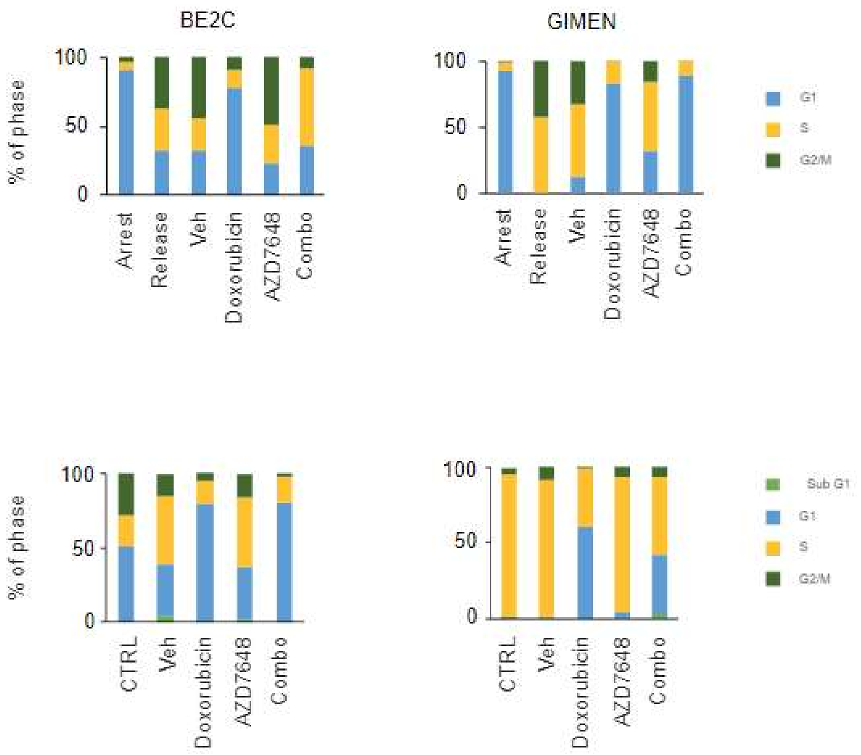


**Extended Data Figure 7. Cell cycle analysis in BE2C and GIMEN**

After 48 hr serum starvation, the cells were arrested at G0/G1 (a) and without cell synchronization (b), followed by vehicle or drug treatment for 72 hr.  After 72 hr, the cells were analyzed by FACS.

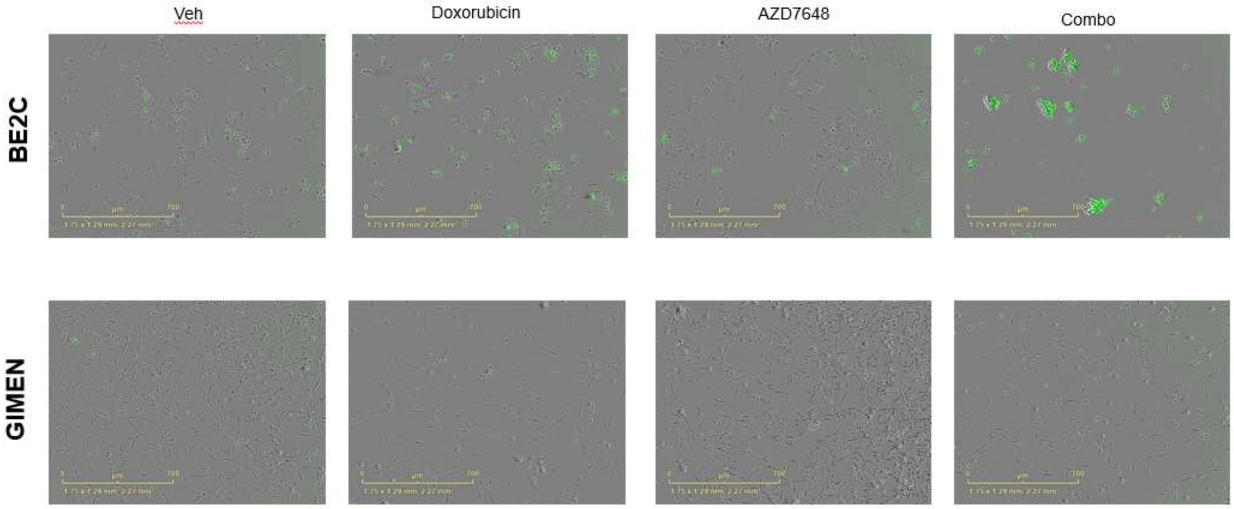


**Extended Data Figure 8. NHEJ activity in BE2C and GIMEN**

Images showing the changes in i-GFP, an indicator of NHEJ 72 hr after vehicle or drug treatment.

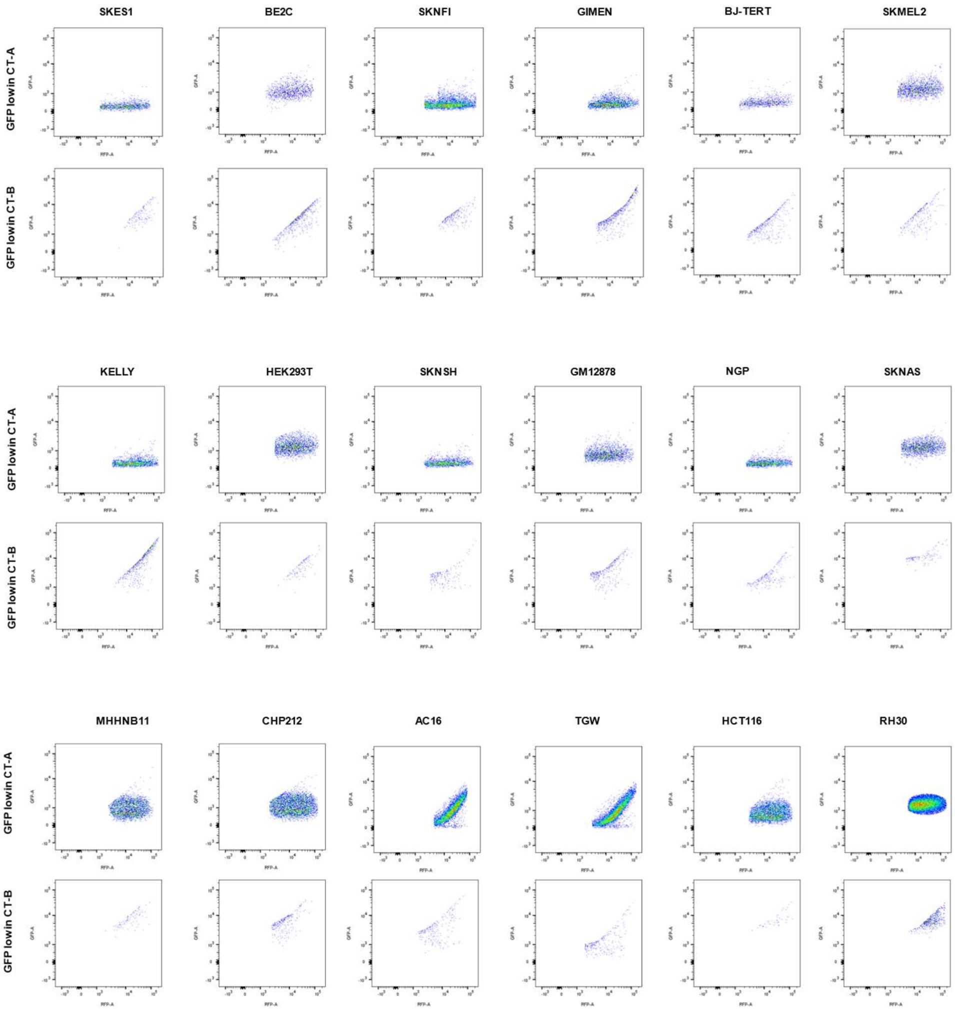


**Extended Data Figure 9. Flow cytometry of GFP negative cells in Cas9 expressing cell lines**

Comparing cell population with GFP low or negative in CT-A and CT-B directly provides Cas9 activity (see Methods). Associated data sets are "Extended Data Table 13 Cas9 activity.xlsx"

a

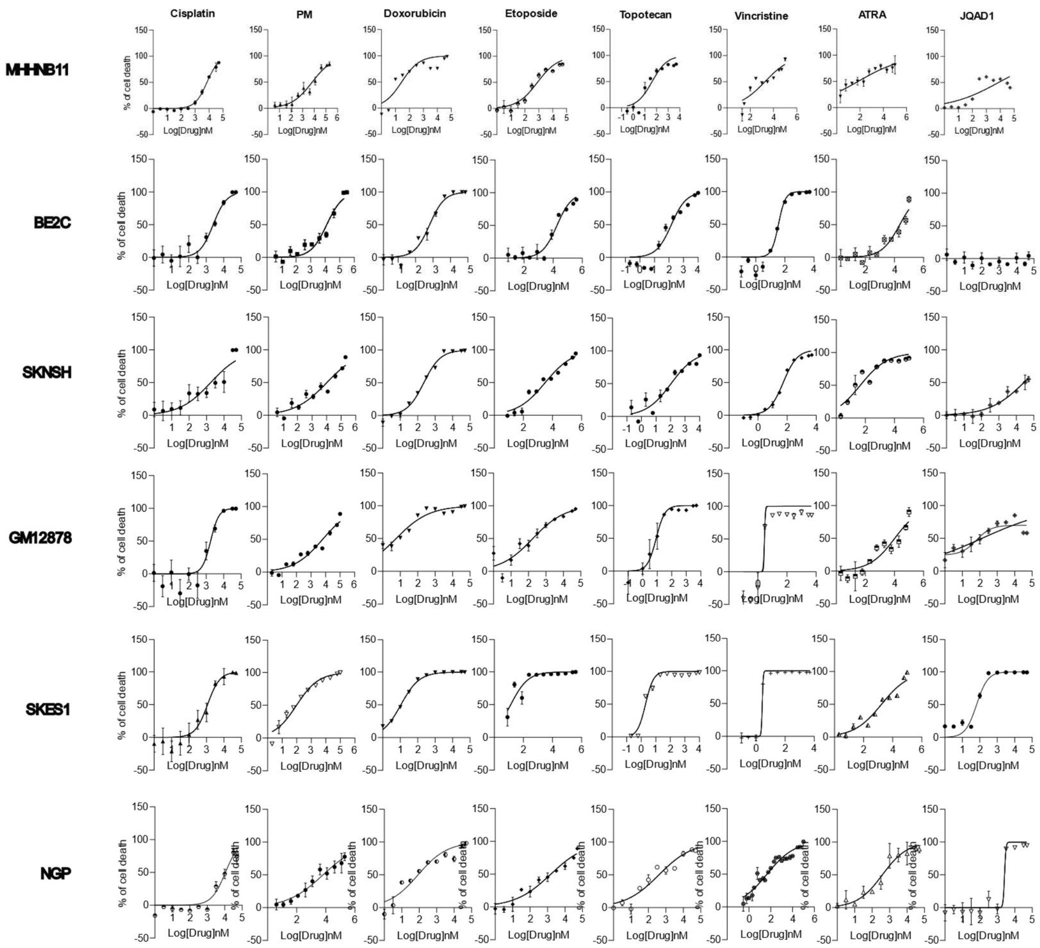
Cisplatin
PM
Doxorubicin
Etoposide
Topotecan
Vincristine
ATRA
JQAD1
MHHNB11
BE2C
SKNSH
GM12878
SKES1
NGP
% of cell death
Log[Drug]nM

**b**

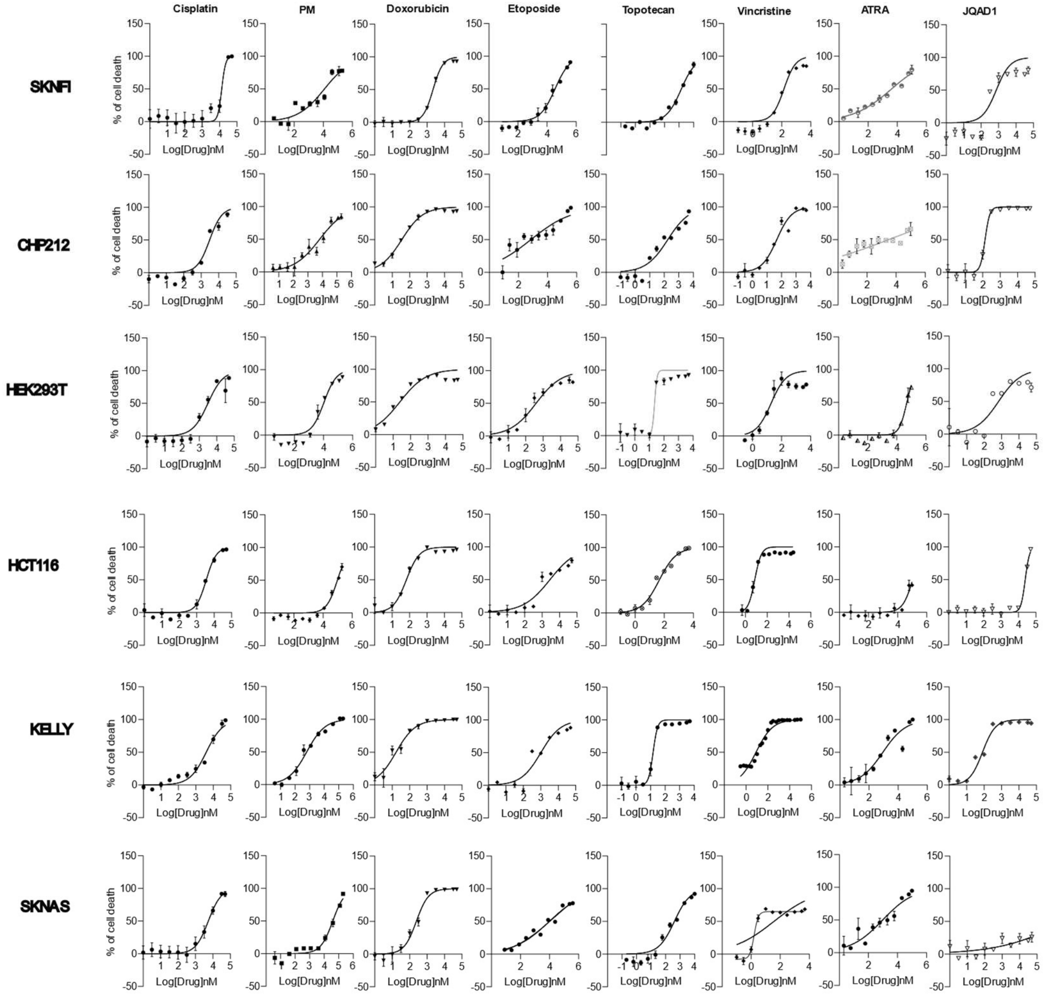
Cisplatin
PM
Doxorubicin
Etoposide
Topotecan
Vincristine
ATRA
JQAD1
SKNFI
CHP212
HEK293T
HCT116
KELLY
SKNAS
% of cell death
Log[Drug]nM

**c**

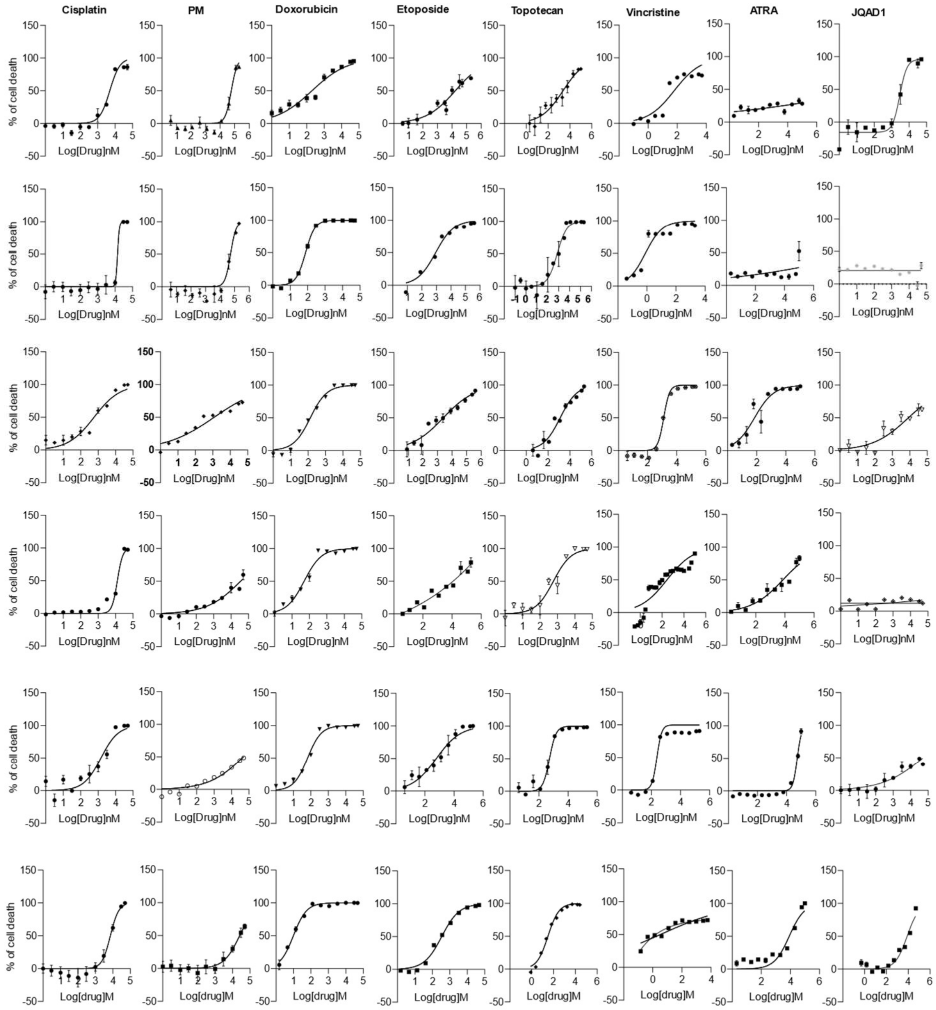


**Extended Data Figure 10. Dose response of 8 drugs in 18 Cas9 expressing cell lines**

(a-c) Dose dependent curves of 8 drugs and 18 Cas9 expressing cell lines to determine $IC_{20}$-$IC_{30}$ for CRISPR screen.

a b

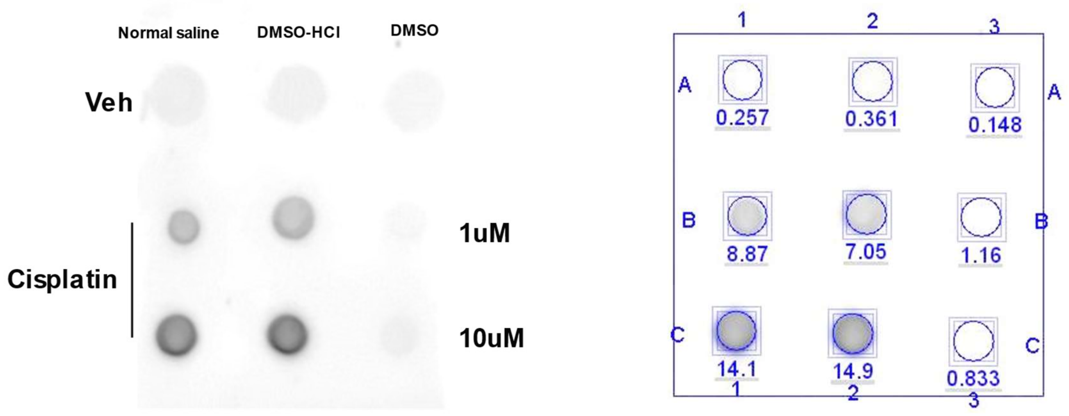


**Extended Data Figure 11. Cisplatin-DNA adduct**

(a) Dot blot of cisplatin-DNA adduct. (b) Measured intensity of each dot. DMSO-HCl overcomes instability of cisplatin in normal saline and activity loss of cisplatin in DMSO alone.
